# First 3D-Structural Data of Full-length Rod-Outer-Segment Guanylyl Cyclase 1 in Bovine Retina by Cross-linking/Mass Spectrometry

**DOI:** 10.1101/2020.09.25.312835

**Authors:** Anne Rehkamp, Dirk Tänzler, Christian Tüting, Panagiotis L. Kastritis, Claudio Iacobucci, Christian H. Ihling, Marc Kipping, Karl-Wilhelm Koch, Andrea Sinz

**Author notes:** Address correspondence to: Prof. Dr. Andrea Sinz, Department of Pharmaceutical Chemistry & Bioanalytics, Institute of Pharmacy, Charles Tanford Protein Center, Martin Luther University Halle-Wittenberg, Kurt-Mothes-Str. 3a, 06120 Halle/Saale, Germany.

## Abstract

The rod-outer-segment guanylyl cyclase 1 (ROS-GC1) is a key transmembrane protein for retinal phototransduction. Mutations of ROS-GC1 correlate with different retinal diseases that often lead to blindness. No structural data are available for ROS-GC1 so far. We performed a 3D-structural analysis of native ROS-GC1 from bovine retina by cross-linking/mass spectrometry (XL-MS) and computational modeling. Absolute quantification and activity measurements of native ROS-GC1 were performed by MS-based assays directly in bovine retina samples. Our data present the first 3D-structural analysis of active, full-length ROS-GC1 in bovine retina. We propose a novel domain organization for the intracellular domain ROS-GC1. Our XL-MS data reveal that the α-helical domain connecting the kinase homology and catalytic domains can acquire different conformations. Also, the XL-MS data of native ROS-GC1 in bovine retina agree with a dimeric architecture. Our integrated approach can serve as a blueprint for conducting 3D-structural studies of membrane proteins in their native environment.

## Introduction

Phototransduction is a process converting the light signal into an electrical response using a signal cascade in the outer segment of the rods in the retina^1–3^. After the activation of rhodopsin via photon absorption, cGMP is hydrolyzed by phosphodiesterases. Subsequently, the decreasing cGMP concentration triggers the closing of the cGMP-dependent cyclic nucleotide-gated channels and the reduction of the intracellular calcium concentration. The rod-outer-segment guanylyl cyclase 1 (ROS-GC1)^4–6^ plays a major role in the calcium feedback mechanism regulated via guanylyl cyclase-activating proteins 1 and 2 (GCAP-1 and −2)^7–10^. In order to restore the calcium concentration to the dark-adapted state, ROS-GC1 raises the initial cGMP level, when calcium levels drop from ~ 500 nM to ~ 50 nM^5,11–13^. In 1991, ROS-GC1, possessing a molecular mass of 110-115 kDa, was purified from different vertebrates^4,14^, but the 3D-structure of full-length ROS-GCs is still elusive.

ROS-GC1 is divided into two main parts, the extracellular (ExtD) and intracellular domains (IcD), separated by the transmembrane domain (TMD) (**Fig. 1**). A large portion of the intracellular domain (IcD) is a kinase-like domain, also known as kinase homology domain (KHD), which is *C*-terminally followed by the catalytic domain (CD)^6,15^. The identity of the ROS-GC1 IcD compared to other members of the membrane guanylyl cyclase family is ~ 34% for the KHD and ~ 57% for the CD, as exemplified for natriuretic peptide receptors and the heat-stable enterotoxin receptor, NPR-A, NPR-B, and STaR^16^.

**Fig. 1.**
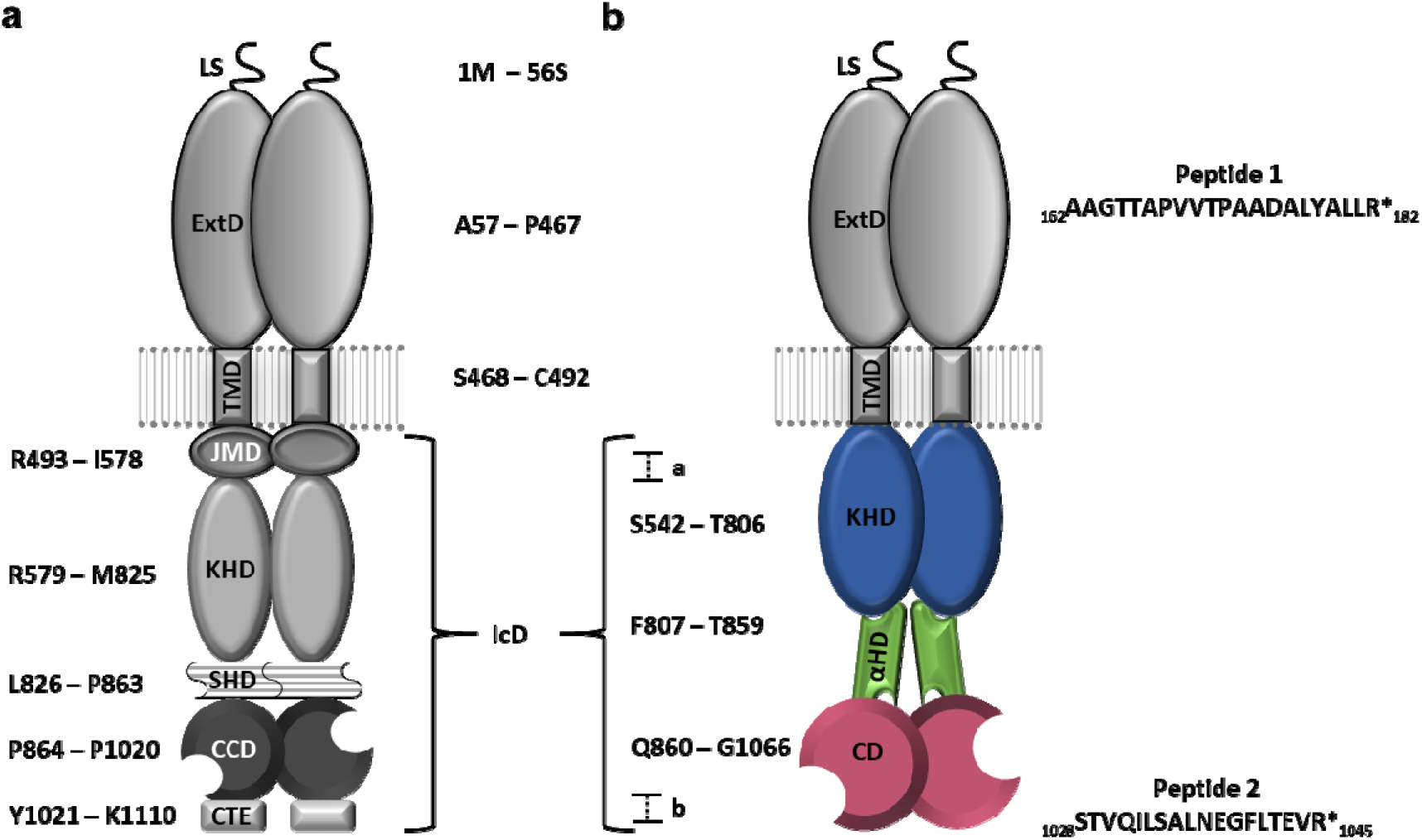
Schematic Presentation of ROS-GC1 Dimer. ROS-GC1 comprises of an extracellular domain (ExtD), a transmembrane domain (TMD), and an intracellular domain (IcD). **(a)** According to Duda *et al*.^28^ the numbering includes the leader sequence (LS). The IcD is subdivided into a juxtamembrane domain (JMD), a kinase homology domain (KHD), a signaling helix domain (SHD) and a core catalytic domain (CCD) with a *C*-terminal extension (CTE). **(b)** Based on the present study, we now divide the IcD into KHD (blue), α-helical domain (αHD, green) and catalytic domain (CD, purple). Boundaries of IcD (marked a and b) are not covered in our current structural working model of the ROS-GC1 IcD. Figure is adapted from ^82^. Isotope-labeled peptides 1 and 2 were used for absolute quantification of bovine ROS-GC1 in ROS preparations.

The classification, organization and overall molecular description of IcD subdomains in ROS-GC1 have evolved during the last years (**Fig. 1a**)^17^: The *N*-terminal part of the KHD was renamed juxtamembrane domain (JMD) as its sequence is highly conserved in ROS-GC1 and 2, but differs from other membrane guanylyl cyclase members, such as NPR-A, NPR-B and STaR^12,18^. The JMD is followed by a kinase-like domain, also known as kinase homology domain (KHD), and the catalytic domain (CD). KHD and CD are separated by a linker region, which exhibits 71% sequence homology to the dimerization domain of the NPR-A receptor^19,20^. Several studies suggest that the linker region is essential for ROS-GC1 dimerization and regulation by GCAPs^21–23^ and adopts either a coiled-coil^23^ or α-helical^24,25^ structure. Depending on its function, the linker region has been termed dimerization domain (DD)^20^ or signal helix domain (SHD)^26,27^. The linker region is followed by the CD that catalyzes the conversion of GTP into cGMP^11^ and the *C*-terminal extension (CTE, aa 1021 - 1110) that is characteristic for ROS-GC1^20^, but is not required for catalytic activity^28,29^. Based on bioinformatics analysis and our present 3D-structural studies, we now propose a novel and simplified domain organization for the IcD of ROS-GC1 that is divided into KHD, *“α-helical domain”* (αHD) and CD (**Fig. 1b**).

A number of retinal dysfunctions and diseases, such as Leber’s congenital amaurosis (type LCA1) and cone-rod dystrophies (CORD), correlate with more than 140 mutations in the *GUCY2D* gene encoding ROS-GC1 (alternatively dubbed GC-E), which cause blindness or severe visual impairment^30–33^. As such, the LCA1 mutation F565S in the *GUCY2D* gene^30^ results in a loss of sensitivity for GCAP-1 regulation when tested *in vitro* with the bovine ortholog^34^. CORD-related mutations can cause different functional impairments as a complete loss of activity, a shift in Ca^2+^-sensitive regulation by GCAPs or constitutive activity^35^. To design novel therapies against these retinal diseases, 3D-structural information on full-length ROS-GC1 is urgently needed.

Chemical cross-linking in combination with mass spectrometry (XL-MS)^36–38^ presents a powerful technique to derive information on the elusive 3D-structure of ROS-GC1 (**Fig. 2**). Chemical cross-linkers react with specific functional groups of amino acids to fix the 3Dstructures of proteins and protein complexes. The major benefit of XL-MS consists in its outstanding potential to derive structural information of proteins and protein networks directly from cell lysates, intact cells, organelles, and tissues^39–42^. The constraints derived from the cross-links then serve as basis for a computational modeling of the protein 3D-structures^43–46^. For our XL-MS studies of ROS-GC1, we applied the MS-cleavable cross-linker disuccinimidyl dibutyric urea (DSBU) that connects mainly primary amine groups of lysines within Cα-Cα distances up to 30 Å^47–49^. Conducting the cross-linking reaction directly in bovine ROS preparations allowed mapping the 3D-structure of ROS-GC1 in its native cellular environment. In this paper, we present the first 3D-structural model of full-length ROS-GC1 by integrating XL-MS and molecular modeling. Our cross-linking data allow refining the domain organization of ROS-GC1 and confirming that ROS-GC1 exists in a native dimeric state.

**Fig. 2.**
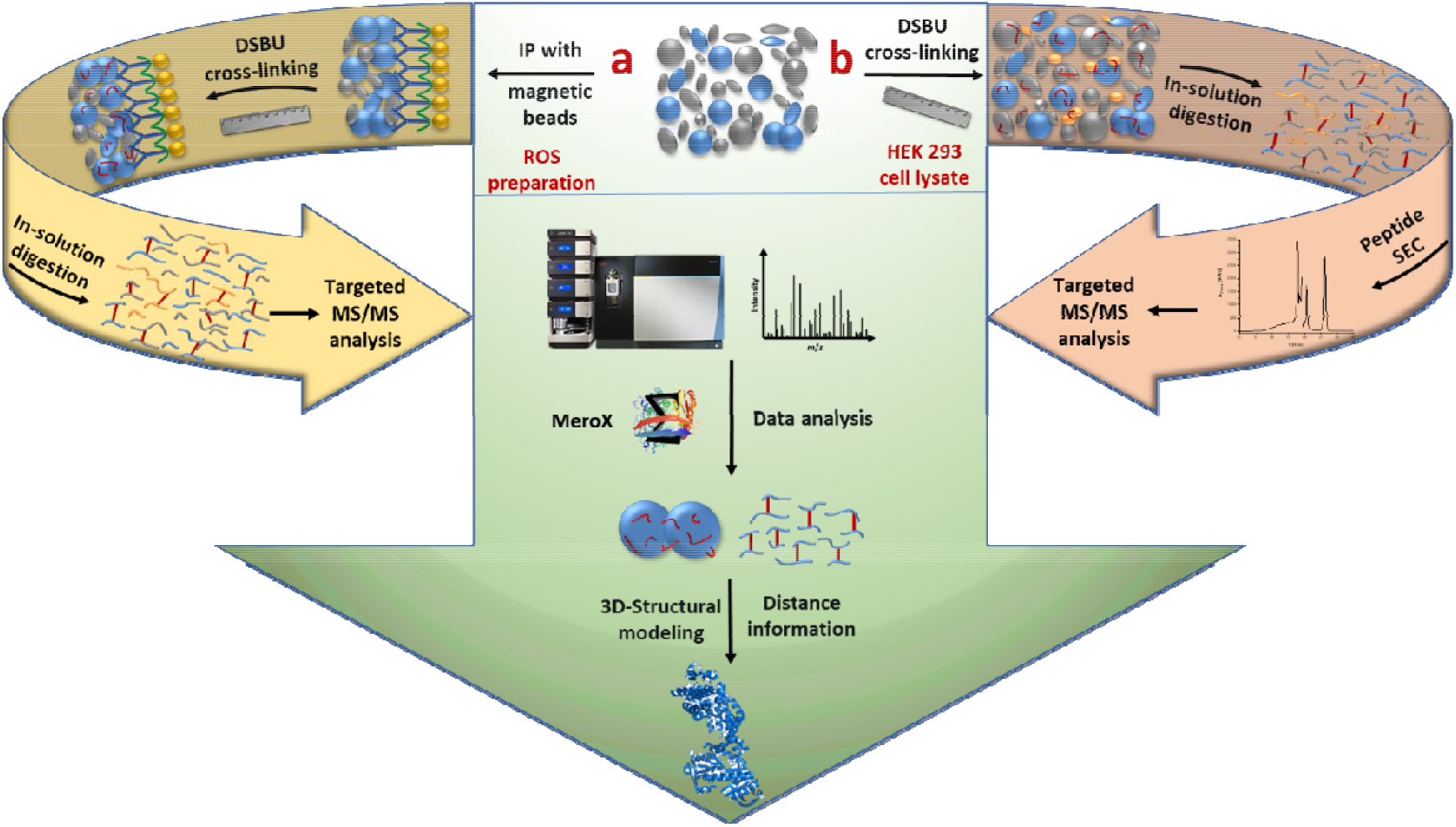
XL-MS Workflow. **(a)** ROS-GC1 was enriched from the bovine rod outer segment (ROS) preparation by immunoprecipitation (IP). Directly on the magnetic beads, the crosslinking with DSBU was performed and subsequently the digestion with trypsin. The supernatant containing the cross-linked peptides was analyzed by LC/ESI-MS/MS. **(b)** After disruption of HEK293 cells transfected with full-length, human ROS-GC, the cell lysate was cross-linked with DSBU and the insoluble fraction was subjected to enzymatic digestion. Afterwards, cross-linked peptides were separated by SEC and the resulting fractions were analyzed by LC/MS/MS. Data analysis was performed with MeroX 2.0. The identified crosslinked amino acids provided distance information for computational modeling of the 3Dstructure of ROS-GC1.

## Results

### Quantification and Activity Measurements of Native Bovine ROS-GC1 in ROS Preparations

As a first step, we characterized the native full-length, bovine ROS-GC in the ROS preparations regarding its absolute concentration and catalytic activity. MS-based quantification of ROS-GC1 was performed using isotope-labeled peptides (*see Methods*)^50–54^. (**Fig. 1b**). The ROS-GC1 concentration was determined to be 102.1 ± 6.7 nmol/l and 129.9 ± 11.5 nmol/l for the two ROS-GC1 peptides by applying a mass spectrometric MRM approach (**Fig. 3**), while it was found to be 100.6 ± 8.0 nmol/l and 130.6 ± 8.1 nmol/l by MS-based analysis (**Supplementary Fig. 1**). Therefore, the ROS-GC1 concentration in the ROS preparations was ca. 115 nmol/l, which is comparable to previous estimations obtained by other methods yielding between 167 and 418 nmol/l^4,55^.

**Fig. 3.**
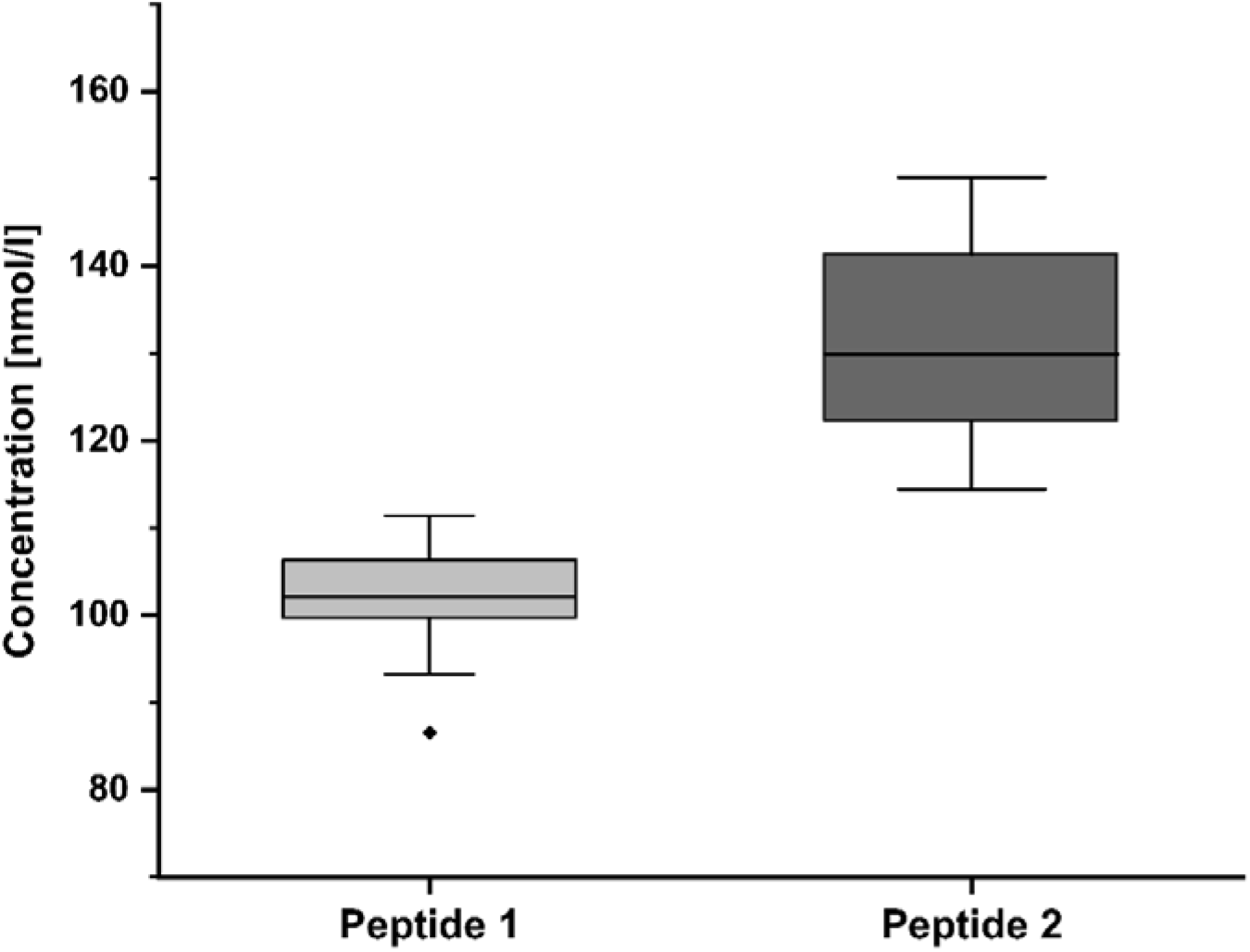
Absolute quantification of Bovine ROS-GC1 in ROS Preparation. Peptides with isotope-labeled Arg at the *C*-terminus, peptide 1 _162_AAGTTAPVVTPAADALYALLR_182_ and peptide 2 _1028_STVQILSALNEGFLTEVR_1045_, were used for absolute quantification of ROS-GC1 (nomenclature of ROS-GC1 peptides, please see **Fig. 1**). Intensity ratios were determined between the peptide that had been generated during digestion and the isotope-labeled peptide that had been externally added. Multiple reaction monitoring (MRM) analysis was performed by the software Skyline 20.1.0.155^52,53^. Concentrations of 102.1 ± 6.7 nmol/l (peptide 1) and 129.9 ± 11.5 nmol/l (peptide 2) were determined.

To ensure that ROS-GC1 derived from ROS preparations and HEK293 cell membranes is catalytically active, activity measurements were conducted by MS-based assays. Quantification of the ROS-GC1 reaction product cGMP, as well as GMP and guanosine, was performed by a mass spectrometric MRM approach (**Supplementary Fig. 2**)^56^. The activity of human ROS-GC1 contained in the HEK293 cell membranes, calculated based on cGMP levels generated, were found to be comparable with previous studies^25,57,58^ (**Supplementary Fig. 3**). ROS preparations were exposed to ambient light to activate phototransduction and to measure GC activity under physiologically relevant light conditions; cGMP generated by ROS-GC1 was degraded via phosphodiesterases (PDEs) to GMP, followed by the conversion of GMP to guanosine (**Supplementary Fig. 4a**). Addition of the PDE inhibitor IBMX could only decelerate, but not abolish, cGMP degradation. Therefore, cGMP, GMP, and guanosine levels were added to yield the total ROS-GC1 activity. To calculate the amount of GMP and guanosine formation, which were generated independently of the ROS-GC1 activity, but directly from GTP via residual enzymatic background activity in the ROS preparations, several control experiments were performed (**Supplementary Fig. 4**). The reaction products derived from isotope-labeled GTP were analyzed in the presence of excess non-labeled cGMP. These high cGMP levels induce a product inhibition of ROS-GC1 and present the major substrate source for PDEs up to reaction times of 10 min (**Supplementary Fig. 4b**). Under these conditions, no labeled cGMP was produced. The activity of full-length, bovine ROS-GC1 was determined to be 7.6 ± 2.2 nmol cGMP x min^−1^ x mg rhodopsin^−1^ (Rh) at a reaction time of 5 minutes (**Fig. 4**), which is in good agreement with published data for complete ROS preparations measured by other methods (~ 7-11 nmol x min^−1^ x mg Rh^−1^)^11,58^. Our careful MS-based analysis allowed us confirming the catalytic activity of ROS-GC1 in ROS preparations and made us confident to perform subsequent XL-MS studies for deriving 3D-structural information of full-length, native ROS-GC1.

**Fig. 4.**
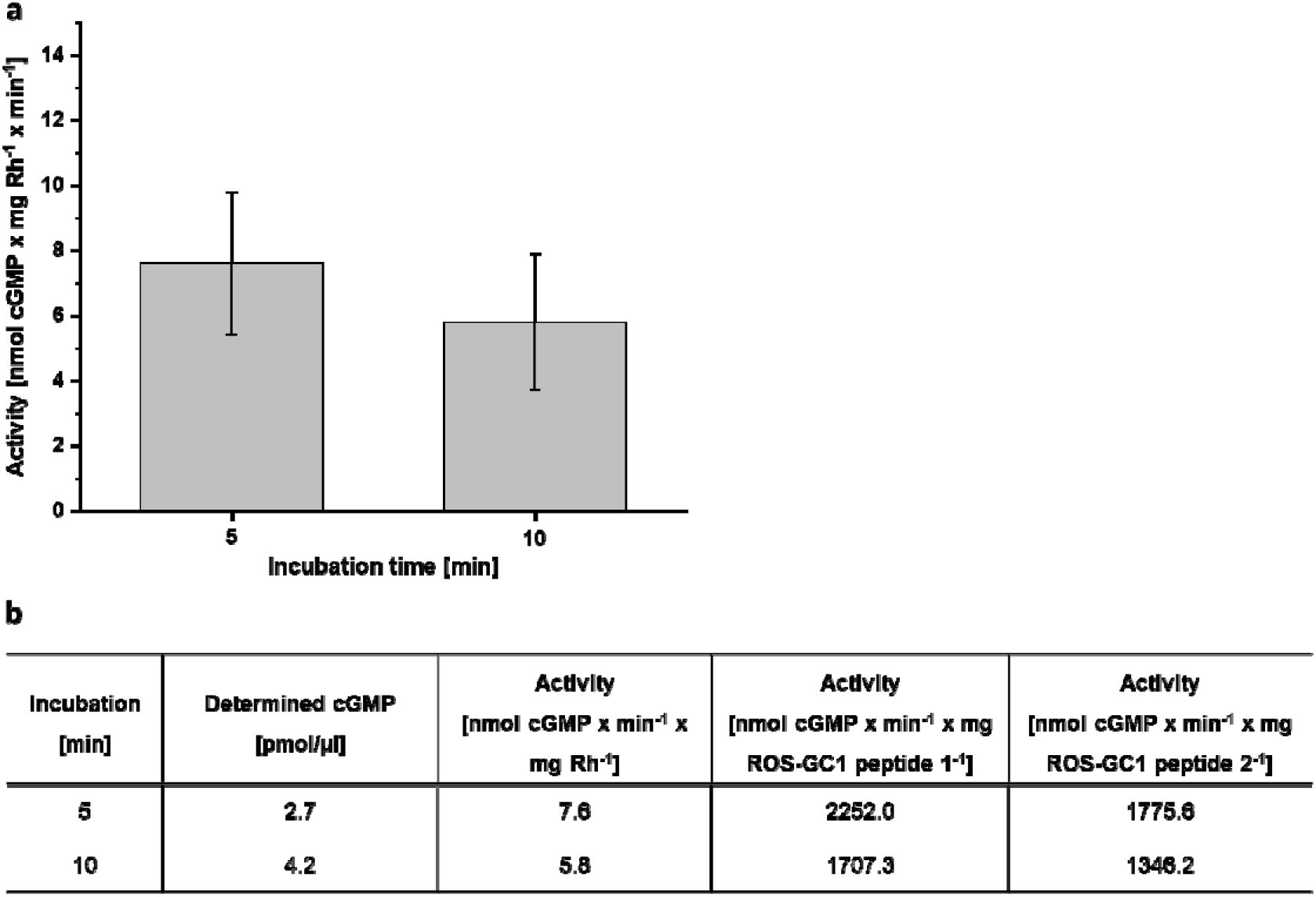
Determination of the Bovine ROS-GC1 Activity. The ROS preparation was incubated for 5 and 10 minutes in the absence of calcium. Quantification of cGMP, GMP and guanosine was performed based on the specific transition of the selected precursor to a product ion by MRM analysis. Peak areas were determined with Skyline 20.1.0.155^52,75^. For the determination of the activity the produced amount of cGMP, GMP and guanosine were added together. As a control, the amount of GMP and guanosine, generated by residual enzymatic activity in the ROS preparation, was subtracted (**Supplementary Fig. 4**). **(a)** ROS-GC1 activities of 7.6 ± 2.2 nmol cGMP x mg Rh^−1^ x min^−1^ (5 min) and 5.8 ± 2.1 cGMP x mg Rh^−1^ x min^−1^ (10 min) were calculated. **(b)** Overview of determined cGMP concentration (pmol/μl), specific activity based on rhodopsin (Rh), ROS-GC peptide 1 and 2.

### In-Tissue Cross-linking Allows the 3D-Structural Study of Native Bovine ROS-GC1

XL-MS experiments were performed with the amine-reactive, MS-cleavable cross-linker DSBU (**Supplementary Fig. 5**) to investigate the conformation of native and active, fulllength ROS-GC1 in the ROS preparations (**Fig. 1**). Conducting the XL-MS experiments with ROC-GC1 in a highly complex mixture without preceding purification of the protein is challenging, but exhibits the distinct advantage of gaining insights into ROS-GC1’s 3Dstructure in its native cellular environment of the bovine retina. The cross-links identified in native, bovine ROC-GC1 served as distance constraints for molecular modeling of ROS-GC1. As only one cross-link was detected in the ExtD of ROS-GC1, we focused on the 3D-structural characterization of the IcD of ROS-GC1.

The XL-MS workflow for analyzing the 3D-structure of native, bovine ROS-GC1 is presented in **Fig. 2**. To adapt the XL-MS strategy for identifying ROS-GC1 cross-linked peptides directly from the highly complex ROS preparations, we optimized our original XL-MS protocol that had originally been developed for studying purified proteins and protein assemblies^49^. The digestion protocol for ROS-GC1 was optimized in HEK 293 cell lysates and then applied to bovine ROS-GC1 from ROS preparations (*see Supplementary Information*).

In principle, two different strategies were applied: The bovine ROS-GC1 from ROS preparations was enriched via IP before XL-MS was performed. For human ROS-GC1 from HEK293 cells, the workflow was executed in the opposite manner, conducting the crosslinking reaction first and the SEC enrichment step afterwards.

Briefly, cross-linking with the MS-cleavable cross-linker DSBU facilitates the MS-based identification of cross-links from highly complex mixtures, such as tissue samples or whole cell lysates (**Supplementary Fig. 5**). DSBU reacts preferably with lysine side chains, but also possesses a significant reactivity towards serine, threonine, and tyrosine residues. The unique feature of DBSU to be cleaved during MS/MS experiments yields characteristic patterns that are recognized by the MeroX software allowing a fully automated and unbiased assignment of cross-links^59–61^.

All unique cross-linking sites identified in ROS-GC1 from the ROS preparations are summarized in **Table 1, Supplementary Tables 1** and **2** and displayed in **Fig. 5**. Each cross-link is assigned to its respective ROS-GC1 domain where the cross-linking site is located, applying the nomenclature and numbering as displayed in **Fig. 1b**. According to our novel domain organization, the majority of cross-links are now located in the KHD and αHD (**Fig. 5** and **Table 1**). One additional cross-link was found in the ExtD. The main cross-linking sites in the KHD and αHD are Lys-527 (involved in four cross-links) and Lys-686/Thr-688 as well as Lys-815/818 (involved in eight cross-links). The cross-links indicate that KHD and CD domain are apparently connected in a flexible manner.

**Fig. 5.**
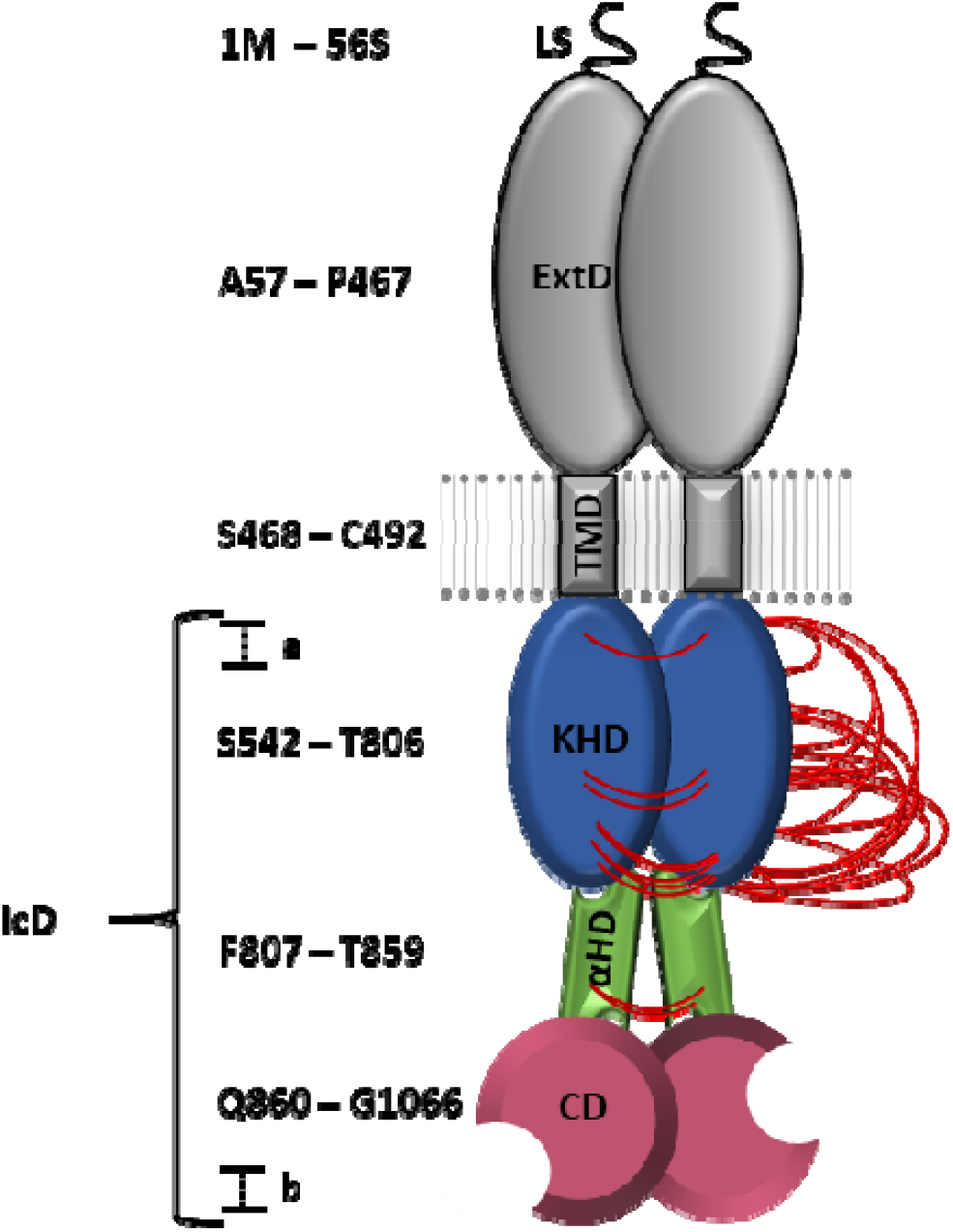
DSBU Cross-links in Native, Bovine ROS-GC1 Dimer. The domain organization was adapted based on the current XL-MS and computational modeling results. Cross-links are shown in red. Figure is adapted from^82^.

**Table 1.**
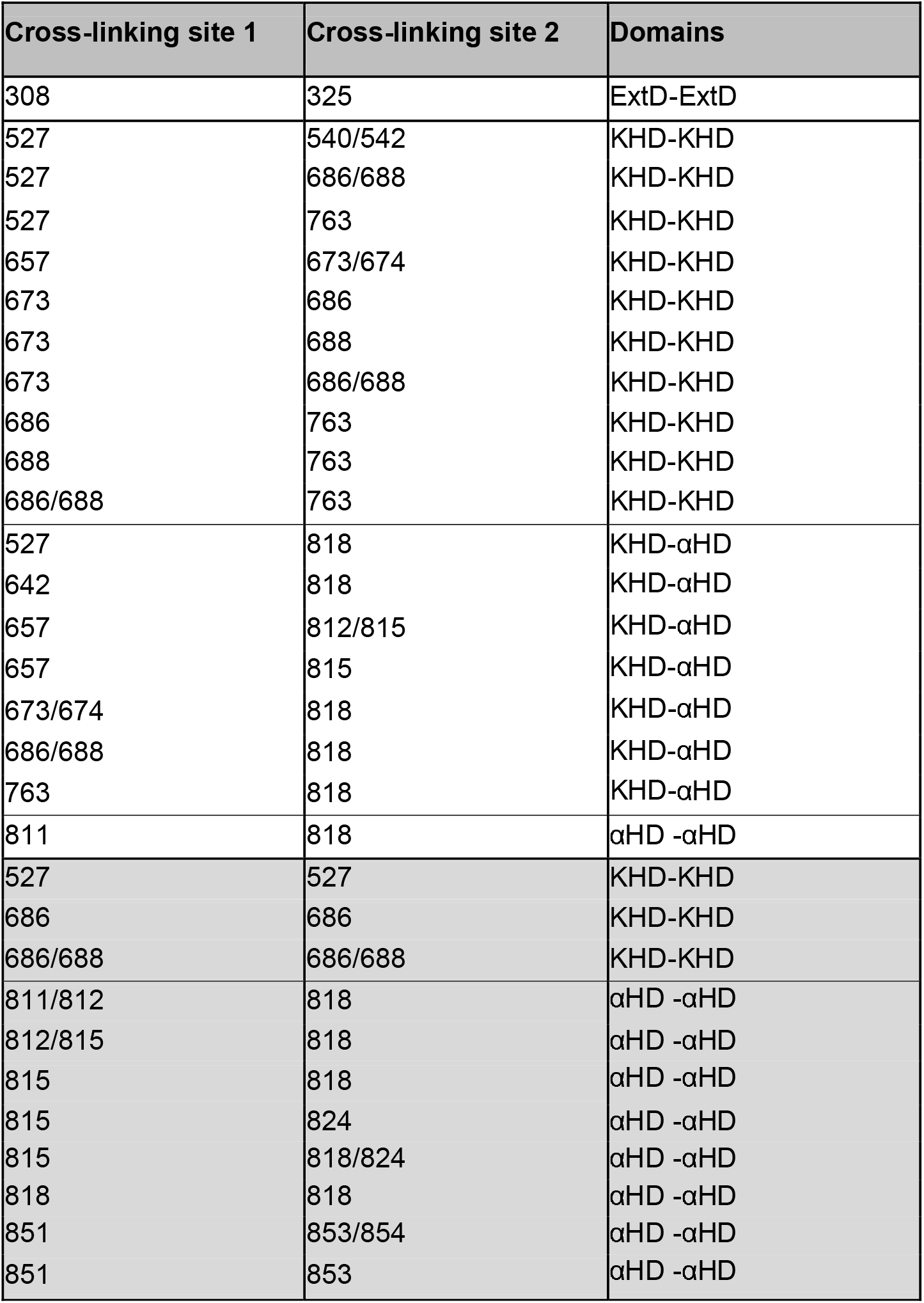
Cross-links in Bovine, Full-length ROS-GC1 from Bovine Retina Preparation. Cross-links between identical or overlapping sequences are considered as interprotein cross-links between two monomers and are shaded in grey. If the cross-linking site is ambiguous, all potential cross-linked amino acids are given. For the nomenclature of ROS-GC1 domains, please see **Fig. 1b**.

| Cross-linking site 1 | Cross-linking site 2 | Domains |
| --- | --- | --- |
| 308 | 325 | ExtD-ExtD |
| 527 | 540/542 | KHD-KHD |
| 527 | 686/688 | KHD-KHD |
| 527 | 763 | KHD-KHD |
| 657 | 673/674 | KHD-KHD |
| 673 | 686 | KHD-KHD |
| 673 | 688 | KHD-KHD |
| 673 | 686/688 | KHD-KHD |
| 686 | 763 | KHD-KHD |
| 688 | 763 | KHD-KHD |
| 686/688 | 763 | KHD-KHD |
| 527 | 818 | KHD-αHD |
| 642 | 818 | KHD-αHD |
| 657 | 812/815 | KHD-αHD |
| 657 | 815 | KHD-αHD |
| 673/674 | 818 | KHD-αHD |
| 686/688 | 818 | KHD-αHD |
| 763 | 818 | KHD-αHD |
| 811 | 818 | αHD -αHD |
| 527 | 527 | KHD-KHD |
| 686 | 686 | KHD-KHD |
| 686/688 | 686/688 | KHD-KHD |
| 811/812 | 818 | αHD -αHD |
| 812/815 | 818 | αHD -αHD |
| 815 | 818 | αHD -αHD |
| 815 | 824 | αHD -αHD |
| 815 | 818/824 | αHD -αHD |
| 818 | 818 | αHD -αHD |
| 851 | 853/854 | αHD -αHD |
| 851 | 853 | αHD -αHD |

Furthermore, 11 unique cross-links were identified between identical or overlapping amino acid sequences, located in the KHD and αHD, and can therefore only by explained by the existence of a ROS-GC1 homodimer in the bovine retina (**Table 1**). These findings are in agreement with previously reported, initial XL-MS data in combination with Western blot analysis where the homodimer was hypothesized to be the active form of native GC in ROS preparations^21^.

### Full-Length Human ROS-GC1 from HEK293 Cells Recapitulates In-Tissue Cross-links

Additionally, we performed XL-MS of ROS-GC1 contained in HEK293 cell membranes as a comparison if this HEK cell overexpressing system will yield data comparable to native ROS-GC1 from bovine retina. This will offer the chance to use HEK293 cell membranes instead of ROS preparations for future structural studies of ROS-GC1.

Apparently, the cross-linking reaction conducted in HEK293 cell lysates was successful (**Supplementary Fig. 6**). SDS-PAGE analysis of non-cross-linked and cross-linked samples revealed a shift towards high molecular weight bands in the DSBU-cross-linked HEK293 cell lysate.

All unique cross-linking sites identified for human ROS-GC1 from HEK293 cells are summarized in **Table 2**, **Supplementary Tables 3**, **4** and **5** and shown in **Supplementary Fig. 7**. As the numbering of human and bovine ROS-GC1 differs by five amino acids, we decided to present all cross-linked amino acids according to bovine numbering (**Supplementary Table 6**).

**Table 2.** Cross-links in Human, Full-length ROS-GC1. Cross-links between identical or overlapping sequences are considered as interprotein cross-links between two monomers and are shaded in grey. If the cross-linking site is ambiguous, all potential cross-linked amino acids are listed. For the nomenclature of ROS-GC1 domains, please see **Fig. 1b.**

| Cross-linking site 1 | Cross-linking site 2 | Domains |
| --- | --- | --- |
| 524/525 | 527 | KHD - KHD |
| 527 | 540 | KHD - KHD |
| 527 | 540/542 | KHD - KHD |
| 527 | 542 | KHD - KHD |
| 527 | 546 | KHD - KHD |
| 527 | 546/549 | KHD - KHD |
| 527 | 549 | KHD - KHD |
| 527 | 546/549/554 | KHD - KHD |
| 527 | 568 | KHD - KHD |
| 527 | 657 | KHD - KHD |
| 532 | 568 | KHD - KHD |
| 568 | 657 | KHD - KHD |
| 657 | 673 | KHD - KHD |
| 657 | 673/674 | KHD - KHD |
| 673/674 | 700 | KHD - KHD |
| 527 | 818 | KHD-αHD |
| 642 | 815 | KHD-αHD |
| 657 | 815 | KHD-αHD |
| 657 | 818 | KHD-αHD |
| 657 | 819 | KHD-αHD |
| 657 | 818/819 | KHD-αHD |
| 815 | 818 | αHD - αHD |
| 527 | 1098 | KHD - CD |
| 588 | 1098 | KHD - CD |
| 851 | 853 | αHD - αHD |
| 1051 | 1051 | CD - CD |
| 1048/1051 | 1053 | CD - CD |

The majority of cross-links were identified in the KHD (**Table 2, Supplementary Fig. 7**). One major cross-linking site is again Lys-527, with 10 cross-links being located in the KHD.

Moreover, Lys-657 (KHD) and Lys-815/818 (αHD) were found to be frequently cross-linked with residues in the *N*-terminal part of the IcD. Structurally relevant cross-links are located between Lys-527 or Lys-588 (KHD) and Lys-1098 (CD), which are located spatially apart from each other in the ROS-GC1 domain structure (**Fig. 1**). That specific cross-link was however not detected in native bovine ROS-GC1.

Our XL-MS results clearly indicate that in the cellular environment in HEK293 cells, the KHD and CD come very close to each other as otherwise they could not be cross-linked. Furthermore, three interprotein cross-links in CD were identified between identical or overlapping amino acid sequences, which again confirm the formation of a ROS-GC1 homodimer under cellular conditions (**Table 2**).

Impressively, the cross-link (Lys-851 x Lys-853) in CD and all cross-linking sites - with the exceptions of Lys-686/Thr-688, Lys-763, Lys-811/Ser-812 obtained from native bovine ROS-GC1 - were confirmed in the human ROS-GC1. Therefore, it can be concluded that fulllength, human ROS-GC1 overexpressed in HEK293 cells adopts a comparable 3D-structure to that of the native bovine ROS-GC1 from ROS preparations and might alternatively be used for future structural studies.

### *C*-Terminal Region of Bovine ROS-GC1 Confirms Higher-Order Architecture

To gain additional structural information of ROS-GC1, we selected a *C*-terminal fragment from bovine ROS-GC1, comprising the αHD and CD (aa 814-1110), for further XL-MS experiments (**Supplementary Fig. 8**). The catalytic activity (14 pmol x min^−1^ x mg^−1^ ROS-GC1) of this fragment was comparable to previously published results where the activity of soluble ROS-GC1 constructs was ~ 7 pmol x min^−1^ x mg^−1^ ROS-GC1^62,63^. The ROS-GC1 fragment was expressed in *E. coli*, purified by affinity chromatography and SEC, enzymatically digested, and subjected to XL-MS in the same manner as bovine and fulllength ROS-GC1 (see *Methods*). A total of 37 unique cross-linking sites were identified for this *C*-terminal ROS-GC1 region comprising the αHD and CD. The majority of cross-links were found in the CD (**Supplementary Tables 7** and **8** and **Supplementary Fig. 9**). Selected MS/MS spectra of DSBU cross-links are shown in **Supplementary Fig. 10**. Five cross-links derived from ROC-GC1 dimer (Lys-1051 x Lys-1051, Lys-1048/1051 x Lys-1053, Lys-1051 x Lys-1053, Lys-1098 x Lys-1101 and Lys-1110 x Lys-1110) contain identical sequence stretches and once again confirm the homodimeric structure of ROS-GC1.

### Molecular Modeling and Cross-link Validation for ROS-GC1 Reveal the Spatial Organization of a Flexible Homodimer

According to our novel domain organization (**Fig. 1b**), the IcD of ROS-GC1 consists of three domains: KHD (aa 542 – 806), CD (aa 860 – 1066), connected by αHD (aa 807-859). The αHD contains a helix-turn-helix motif with a hinge (**Fig. 6** and **Fig. 7**). Our XL-MS experiments identified ROS-GC1 to exist as a homodimer in its native form and the KHD and CD to be connected in a flexible manner.

**Fig. 6.**
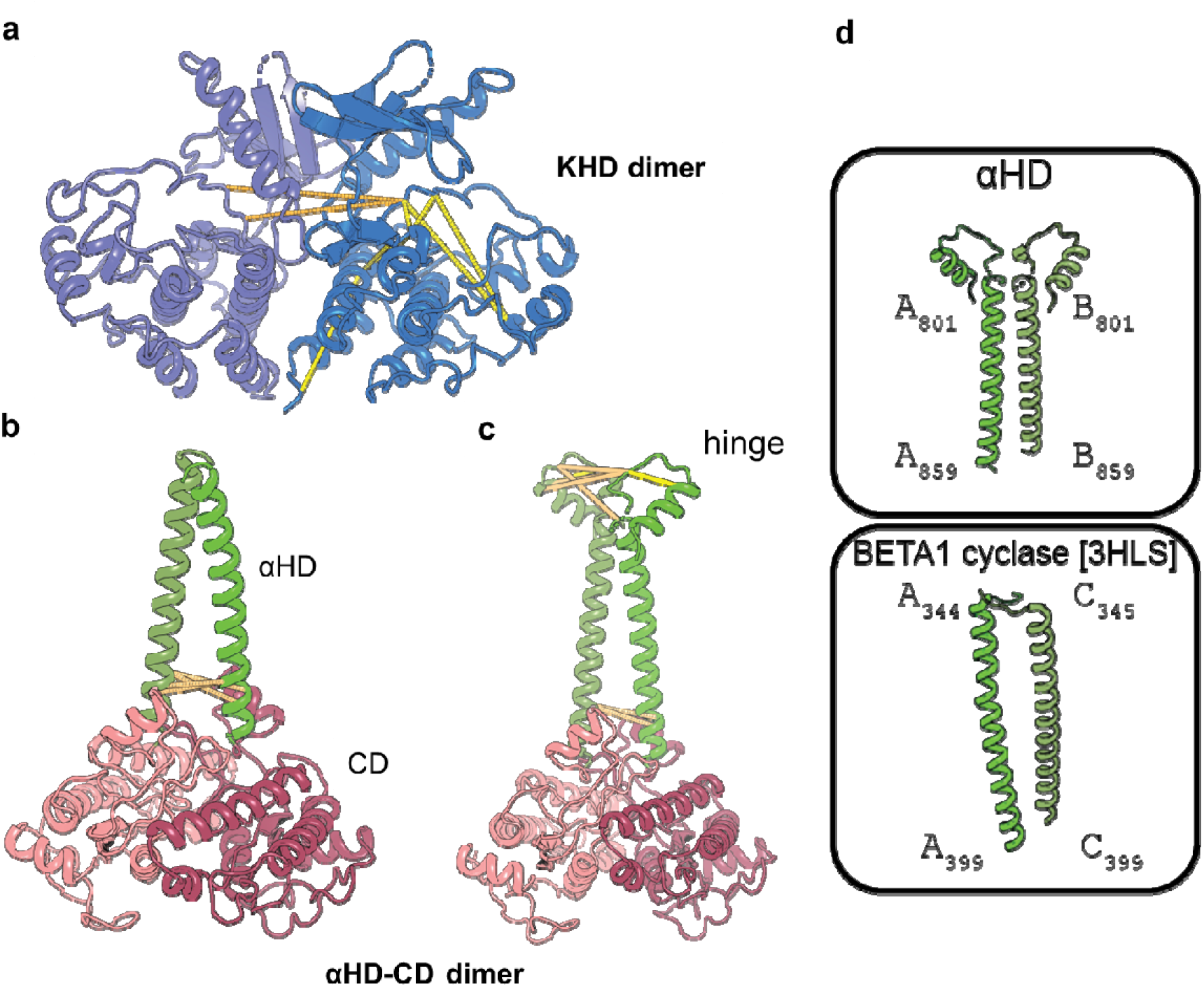
Dimeric models for the KHD, αHD and CD domains and satisfaction of all identified cross-links on the structural models. **(a)** Dimeric KHD domain, derived from binary HADDOCK docking by applying C2 symmetry. **(b)** Initial dimeric αHD-CD domain. **(c)** Extended dimeric αHD-CD, now including the molecular hinge. The initial structure was extended by adding a helix-turn-helix motif at the *N*-terminal region. The derived structure was validated by 6 cross-links. Cross-links are shown as dashed lines. Intra- and intermolecular cross-links are colored in yellow and orange, respectively. **(d)** Comparison of the αHD from ROS-GC1 and the β1 subunit of the soluble guanylyl cyclase (BETA1 cyclase, 3HLS). Both structures show a helical structure, with a turn at the *N*-terminal end.

**Fig. 7.**
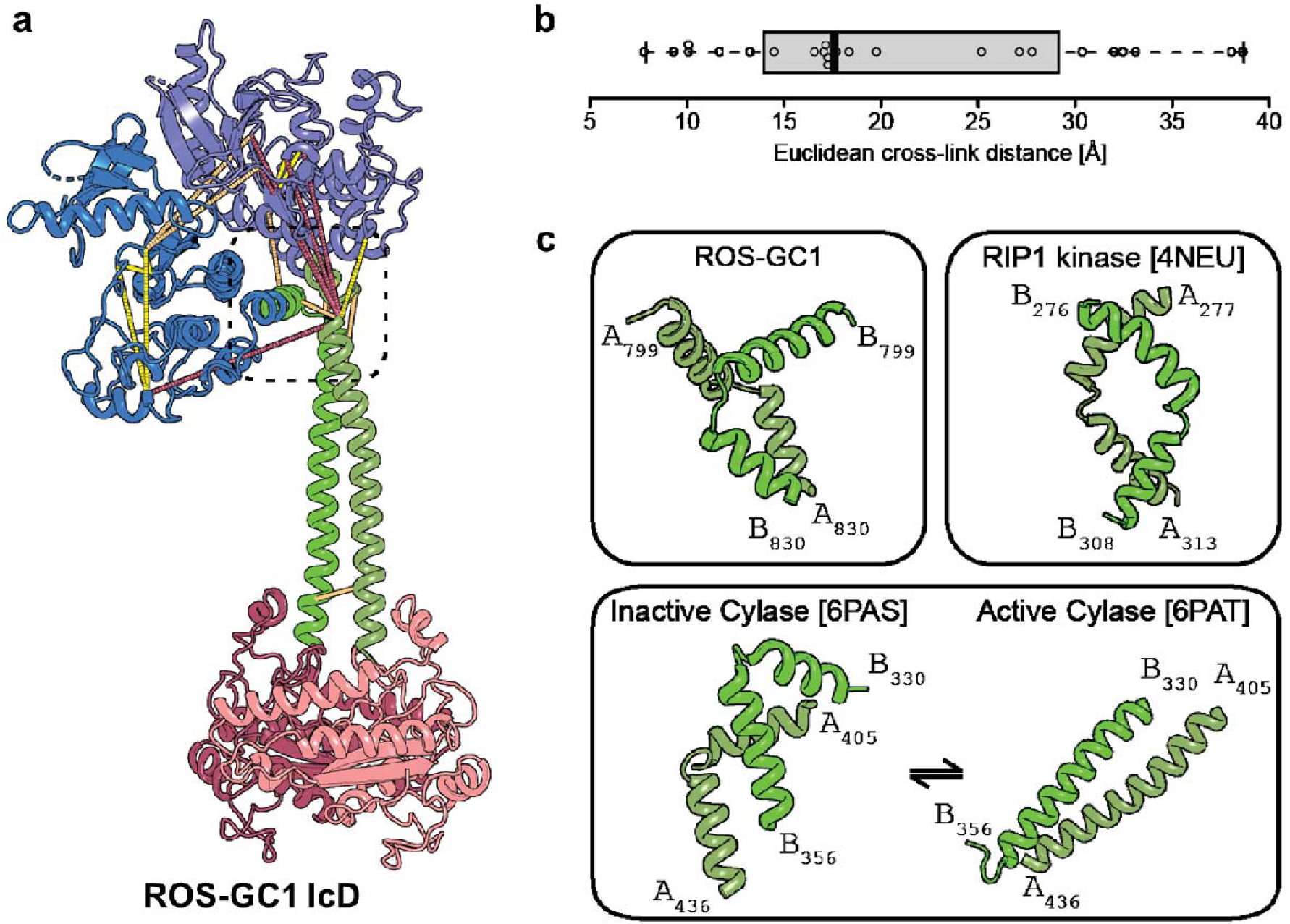
Model of the complete dimeric intracellular domain (IcD) of ROS-GC1. **(a)** Dimeric model of the ROS-GC1 IcD. Cross-links are shown in dashed lines and colored by their type: intra-molecular cross-links are shown in yellow, inter-molecular in orange and violated cross-links (both, intra- and intermolecular) in red. **(b)** Statistics of the Euclidean distance distribution. The median distance is indicated at 15.60 Å. **(c)** Representative conformations for the flexible helix-turn-helix motif in kinases and cyclases. On the top left, the motif included in the ROS-GC1 structure generated in this study is shown; on the top right, the motif from the RIP1 kinase (PDB-ID: 4NEU) is shown. On the bottom, this motif can undergo structural changes upon activation as shown in a cyclase (PBD-IDs: 6PAS, 6PAT). The helix-turn-helix manifests in the inactive conformation (6PAS) and the same sequence acquires an extended helical conformation in the active state. Residue numbers of *N-* and *C*-terminus of the regions are given.

The KHD is highly conserved in its monomeric state, but a variety of dimeric interfaces exist in the KHD. We identified various dimeric KHD interfaces of high homology, present in the B-raf kinase of the MAPK pathway complex (6Q0J)^64^, the Ser/Thr kinase PknA (4X3F)^65^, the Tyr kinase c-Abl (4XEY)^66^, and the Ser/Thr kinase PBL2 (6J5T)^67^. This implies that kinase dimers can acquire multiple higher-order states, which is also reflected by the XL-MS data. This is because, after superposition and cross-link mapping on the generated KHD dimer models using the templates above, violations of cross-links are observed. Therefore, an unbiased, data-driven binary docking of the monomeric structures using HADDOCK leads to a KHD dimer with a complete satisfaction of cross-linking constraints (**Fig. 6a**; **Supplementary Table 9**).

The CD is also highly conserved, as shown for a variety of homologous protein complexes. Additionally, the αHD is oriented in a perpendicular manner towards the CD and is conserved in multiple models of those homologous structures. In contrast to the KHD, the dimeric interface of the αHD-CD structure is highly conserved. Therefore, based on the dimeric structure of a soluble guanylate cyclase from *Manduca sexta* (6PAS)^68^, we generated the dimeric structure of the αHD and CD (aa 823 – 1065, **Fig. 6b**). Intermolecular cross-links identified (Lys-851 x Lys-853 or Lys-854) validate the αHD/CD dimeric model. The *N*-terminal region of the αHD extends in a helix-turn-helix motif to connect the KHD and CD. This helix-turn-helix motif (aa 800 - 825) is highly conserved in various structural models deposited in the PDB (4NEU, 6OKO, 4UYA, 6PAT, 6PAS), but acquires multiple conformations involving helical elements, forming a molecular hinge. The consequence of these structural changes is the subsequent spatial and conformational alteration of the interconnected domains. A similar conformational change might occur in ROS-GC1, defining the orientation of the CD, present at the *C*-terminus of the IcD, while the *N*-terminus is anchored in the membrane. The model of the αHD has therefore been extended by this helix-turn-helix or hinge motif (aa 800 – 1066; **Fig. 6c**). The derived structure satisfied one intra- and five intermolecular cross-links (**Fig. 6c**). The overall structure of the complete αHD is in a good agreement (RMSD 7.06 Å) with the known αHD from the β1 subunit of the soluble guanylyl cyclase (**Fig. 6d**). We successfully completed the ROS-GC1 IcD structure after structural superposition of the derived KHD and αHD/CD dimers after subsequent refinement using HADDOCK (**Fig. 7a**). In addition, the final ROS-GC1 IcD model satisfies 79% (19/24, **Supplementary Table 10**) of identified cross-links and recapitulates a lognormal distance distribution (**Fig. 7b**). Interestingly, the five violated cross-links are exclusively located in the hinge motif of the αHD and can be rationalized by the intrinsic flexibility of this region. In fact, different conformations have been observed for other cyclase and kinase domains (**Fig. 7c**). Additionally, a conformational change is required for activation as seen in the soluble guanylate cyclase from *Manduca sexta*, which is captured in an active (6PAT) and inactive state (6PAS)^68^. Here, a conformational change from a 90° kinked helix-turn-helix motif to a straight helix motif is apparent (**Fig. 7c**).

The model generated for the complete IcD was further used to validate the cross-links, identified in human ROS-GC1 from HEK293 cells and the recombinant *C*-terminal bovine ROS-GC1 fragment produced in *E*. *coli*. All cross-links (14/26) that can be mapped for ROS-GC1 from HEK293 cells are within the allowed distance range (**Supplementary Table 11**). The majority of cross-links in the *C*-terminal bovine ROS-GC1 fragment (aa 814 – 1110) are located within the *C*-terminal region of the CD (aa 1067 – 1110), which could not be modeled due to the absence of suitable template structures. Interestingly, this *C*-terminal region is predicted to be disordered. Conclusively, only 8 out of 37 cross-links could be mapped onto the native IcD structure with a cross-link satisfaction of 37.5 % (3/8; **Supplementary Table 12**). The violated cross-links are, again, located between the αHD and the CD.

## Discussion

So far, no high-resolution 3D-structure is available for native ROS-GC1. In particular, partition, domain arrangement and stoichiometry of the IcD are still under discussion. The two major domains of ROS-GC1 IcD are the KHD and the CD that are connected by the αHD (**Fig. 1b**)^28,62^. In particular, the dimerization and function of the αHD is currently discussed as GCAP binding interface^21–23^ or as a control switch/signaling domain (SHD)^24,25,27^. Based on our experiments, we propose a role of the αHD in regulating cyclase activity.

### Intracellular Domain of ROS-GC1

In this work, we investigated the 3D-structure of ROS-GC1 by XL-MS and computational structural biology methods and succeeded in deriving a structural model for the complete IcD of ROS-GC1. The cross-links that were independently derived from full-length bovine and human ROS-GC1 were in good agreement, delivering valuable insights into the enzyme’s 3D-structure and structural similarity across species (**Tables 1** and **2** and **Supplementary Fig. 11**). 10 cross-links identified in native bovine ROS-GC1 in ROS preparations conformed with 13 unique cross-links derived from human ROS-GC1 overexpressed in HEK293 cells. Furthermore, 14 cross-links in human ROS-GC1 provided additional structural information, of which 8 were not possible in bovine ROS-GC1 due to differences in the amino acid sequence. Three additional cross-links were not found in human ROS-GC1 caused by the absence of Lys-763 in the human sequence.

### Molecular Modeling Indicates Extensive Conformational Flexibility

The ROS-GC1 IcD model exhibits structural features of striking molecular complexity. In particular, the αHD (aa 807 - 859) that connects the KHD to the CD can acquire various conformations, having an effect on the orientation and proximity of the connected domains. The XL-MS data agree with a dimeric architecture and an extended conformation of the ROS-GC1 IcD, which has been shown to be the catalytically active conformation in other guanylate cyclases^68^ **(Fig. 7c**). This observation is also corroborated by our in-cell assays, which show a significantly higher activity of ROS-GC1 in bovine retina extracts as compared to those derived from ROS-GC1 overexpressed in HEK293 or *E. coli* cells.

Although we cannot exclude a conformation where the KHD and CD are in vicinity in native bovine ROS-GC1, this conformation is expected to be a minor population. No cross-link was observed for vicinal KHD and CD in the bovine ROS-GC1 from ROS preparation. Although the absence of cross-links between these domains does not imply the exclusion of structural states, activity assays corroborate the active conformation and not the closed, inactive state.

Only two cross-links in human ROS-GC1 from HEK293 cell membranes (Lys-527 x Lys-1098 and Lys-588 x Lys-1098) might however indicate the vicinity of KHD and CD. The cross-linking results from a *C*-terminal fragment of bovine ROS-GC1 (aa 814-1110) also suggest a compact structure where KHD and CD are in close neighborhood. We observed cross-links between KHD and CD (Lys-818 x Lys-1051, Lys-853/Thr-854 x Lys-1048/1051, Lys-818 x Lys-1098, and Lys-853/Thr-854 x Lys-1110). The DSBU cross-linker spans C_α_-C_α_ distances up to 30 Å^47–49^. The cross-links exhibit a distance of more than 30 Å in the model, indicating a high flexibility in the regions where both lysine residues are located. Structurally exploring the combined torsion angle space of the αHD and the predicted disordered C-terminal region of the CD does not yield satisfaction of the derived cross-links (**Supplementary Fig. 12a-d**). This points to even higher conformational flexibility underlying both regions. Indeed, violated cross-links in the bovine, full-length ROS-GC1 are exclusively located in the hinge motif and underlying the intrinsic flexibility and importance of this region for activity.

### Validating the Model of the Intracellular Domain

In order to verify our ROS-GC1 IcD model, C_α_–C_α_ distances bridged by the cross-linker were mapped into the KHD, αHD and CD. For the bovine ROS-GC1, the cross-links are in the range of 8 to 30 Å with four exceptions. These four cross-links exhibit distances of slightly higher than 30 Å but are readily explained by their location in the hinge region of the αHD (**Fig. 6** and **Supplementary Table 10**). Furthermore, the cross-links Lys-568 x Lys-657, Lys-642 x Lys-815, Lys-657 x Lys-673, Lys-657 x Ser-674, Lys-657 x Lys-815 and Lys-673 x Lys-700 of the human ROS-GC1 derived from HEK293 cell membranes fit perfectly with the KHD and αHD model showing distances of less than 30 Å (**Supplementary Table 11**). It has to be noted that currently we cannot exclude that these cross-links are intramolecular ones (within one monomer) and not intermolecular (between two monomers). The inter-protein cross-links Lys-851 x Lys-853, Lys-1051 x Lys-1051, Lys-1048 x Lys-1053 and Lys-1051 x Lys-1053 with C_α_-C_α_ distances of less than 30 Å confirm the dimeric ROS-GC1 structure (**Supplementary Table 11**). Strikingly, the majority of cross-links found for the native bovine ROS-GC1 dimer from ROS preparations were identical to those of full-length human ROS-GC1 and a *C*-terminal bovine ROS-GC1 fragment, additionally providing complementary 3D-structural information.

Conclusively, in this study, we established an integrated XL-MS workflow that enabled us to identify cross-linked products from full-length, enzymatically active ROS-GC1 directly in bovine ROS preparations. Cross-links were identified for the complete IcD of ROS-GC-1. 3D-structural information on the KHD and CD of ROS-GC1 was derived from the cross-link constraints serving as basis for subsequent molecular modeling studies. Apparently, the αHD that connects the KHD and the CD can acquire various conformations, having an effect on the orientation and proximity of the connected domains. Our XL-MS data agree with a dimeric architecture and an extended conformation of the ROS-GC1 IcD, which has been shown to be the catalytically active conformation in other guanylate cyclases. To refine our current structural model of full-length ROS-GC1, further experiments will be conducted using alternative cross-linking chemistry and combining complementary structural biology techniques, such as cryo-EM and XL-MS.

## Methods

### Chemicals and Reagents

Chemicals were obtained from Roth or Sigma-Aldrich, solvents were obtained from VWR. ROS-GC1 antibody (B7, monoclonal) was purchased from Santa Cruz Biotechnology, Dynabeads Protein G was obtained from Thermo Fisher Scientific (Invitrogen), and ROS-GC1 peptides (SpikeTides) were obtained from JPT Peptide Technologies.

### Cell Culture

HEK-flip 293 cells that had been permanently transfected with full-length, human ROS-GC1 gene were kindly provided by Prof. Daniele Dell’Orco, University of Verona, Italy. Cells were cultivated in Dulbecco’s modified Eagle’s medium (DMEM, high glucose, GlutaMAX, Gibco by Thermo Fisher Scientific), harvested, and centrifuged as previously described^25,69,70^. ROS-GC1 expression was controlled via fluorescence microscopy of the co-expressed eGFP and ROS-GC1 identification by peptide mass fingerprint analysis. Cell pellets were stored in liquid nitrogen at −80°C before XL-MS experiments or activity assays were conducted.

### Protein Expression and Purification

Additionally, the sequence from the *C*-terminal part of the IcD of bovine ROS-GC1 (aa 814-1110) was expressed with *N*-terminal (His)_8_ tag, lipoyl domain, and tobacco etch virus cleavage site (HLT tag) in *E*. *coli* BL21 (DE3) cells. Cells were grown at 37°C to an OD_600_ of 0.6. Afterwards, cells were incubated at 18°C for 1 h and gene expression was induced by adding 0.1 mM IPTG. Expression was performed at 18°C overnight. After harvesting, the cells were resuspended (50 mM HEPES, 150 mM NaCl, 2.5 mM TCEP, 10% glycerol, 0.05% DDM and 10 mM imidazole, pH 7.2) and disrupted by French press. The supernatant was loaded onto a Ni^2+^-NTA column (5 ml HisTrap FF column, GE Healthcare) and the protein was eluted with an imidazole gradient. Afterwards, size exclusion chromatography (SEC) (Superdex 200 pg 26/600, GE Healthcare) was performed (50 mM HEPES, 150 mM NaCl, 2.5 mM TCEP, 10% glycerol, 0.05% DDM and 1.5 mM MgCl_2_, pH 7.2), fractions of the protein monomer were combined, concentrated, and stored at −20°C.

### ROS Preparations

The ROS of bovine retina containing full-length ROS-GC1 were prepared from bovine eyes balls obtained from a local slaughterhouse according to an existing protocol^71,72^ or were bought from In Vision Biosciences (Seattle, USA). The rhodopsin concentration in the ROS preparations was determined to be 3.65 mg/ml^55^.

### Immunoprecipitation

For immunoprecipitation (IP) of the ROS preparations, magnetic beads (30 μl) with immobilized protein G (Dynabeads protein G, 30 mg/ml) were incubated with 15 μl ROS-GC1 antibody (monoclonal, 200 μg/ml) at room temperature for 60 min. After antibody immobilization, the beads were washed three times with 500 μl of reaction buffer (20 mM HEPES, 100 mM NaCl, 2 mM EGTA, 2 mM MgCl_2_, 2 mM DDM, 10% (*v*/*v*) glycerol, pH 7.5). 20 μl of ROS preparation were solubilized with 230 μl of the reaction buffer on ice for 60 min. Solubilized ROS suspension was incubated with the antibody beads for 60 min and washed three times with reaction buffer (500 μl each). Afterwards, ROS-GC1 bound to the magnetic beads was applied for XL-MS.

### ROS-GC1 Quantification

For absolute ROS-GC1 quantification in the ROS preparations, two representative tryptic ROS-GC1 peptides were employed (peptide 1: _16_2AAGTTAPVVTPAADALYALLR_182_, peptide 2: _1028_STVQILSALNEGFLTEVR_1045_, for nomenclature of ROS-GC1 peptides, please see **Fig. 1**). Both ROS-GC1 peptides were synthesized with stable isotope-labeled (^13^C_6_ and ^15^N_4_), C-terminal Arg, followed by a trypsin-cleavable Q-tag^50,51^. To 2.5 μl of ROS preparation, ProteaseMAX surfactant was added to a total volume of 90 μl. Digestion was performed according to the manufacturer’s protocol (Promega)^73,74^. Before adding 2 μl of trypsin (0.05 μg/μl, Promega), 500 fmol of isotope-labeled peptides 1 and 2 were added to spike the ROS preparation. The reaction was stopped by adding 10 μl of 10 % (*v/v*) TFA. Different sample volumes, corresponding to 5, 10, 25, 50 and 75 fmol of isotope-labeled peptides 1 and 2, were analyzed with the NanoElute UPLC system (Bruker Daltonik) coupled to a timsTOF Pro mass spectrometer equipped with CaptiveSpray Ion Source (Bruker Daltonik). The peptide mixtures were separated via a C18 column (75 μm x 150 mm, 1.9 μm, 120 Å, nanoElute Fifteen, Bruker Daltonik) with a linear 40-min gradient ranging from 2% to 50% solvent B: acetonitrile with 0.08% formic acid (FA); solvent A: 0.1% formic acid (FA) at a flow rate of 0.3 μl/min. MS data were recorded using a multiple reaction monitoring (MRM) approach. Triply charged ions of non-labeled and isotope-labeled peptides 1 and 2 were selected for fragmentation by collision-induced dissociation (CID) with a normalized collision energy of 35%. All MS measurements were performed in triplicate. For absolute ROS-GC1 quantification, intensity ratios were determined of the two intrinsic ROS-GC1 peptides 1 and 2, generated by tryptic digestion, and their externally added, isotope-labeled peptide counterparts. Data analysis was performed with Skyline 20.1.0.155 (MacCoss Lab, University of Washington)^52–54^.

### ROS-GC1 Activity

The catalytic activity of ROS-GC1 in ROS preparations and HEK293 cell membranes was determined according to existing protocols^11,25,58^. The conversion of GTP to cGMP, catalyzed by ROS-GC1, was induced by adding 1 μl of ROS preparation or 5 μl of HEK293 membrane suspension to a final volume of 25 μl of activity buffer (40 mM HEPES, 56 mM KCl, 8 mM NaCl, 10 mM MgCl_2_, 0.1 mM ATP, pH 7.5), containing 10 mM IBMX solution, 1 μM GCAP-2, and 2 mM GTP. After incubation at 37°C for 5 or 10 min, the reaction was stopped by adding 100 mM citric acid and 100 mM EDTA at a 1:1 (*v*/*v*) ratio. Calibration curves were generated by injection of 0.1, 0.25, 0.5, 1, 2, 5, 10 and 20 μM cGMP, GMP and guanosine in activity buffer (**Supplementary Fig. 2**). To determine the basal enzymatic activity in the ROS preparation, which might result in the production of GMP and guanosine, 2 mM cGMP was added to 2 mM ^13^C/^15^N-labeled GTP (**Supplementary Fig. 3**). ^13^C/^15^N-labeled GTP allowed discriminating whether the reaction products GMP and guanosine originally derive from GTP or cGMP.

Quantification of cGMP, GMP, and guanosine was performed by LC/MS using an MRM approach^56^ (nanoAcquity UPLC system coupled to a Xevo TQD mass spectrometer, Waters) using specific transitions of selected precursor ions to fragment ions (cGMP: *m/z* 346.0 → 152.0, *m/z* 361.0 → 162.0; GMP: *m/z* 364.0 → 152.0, *m/z* 379.0 → 162.0; guanosine: *m/z* 284.0 → 152.0, *m/z* 299 → 162.0). The mixtures were separated via a C18 column (100 μm x 1.0 mm, 2.6 μm, 100 Å, Kinetex XB-C18, Phenomenex) with a linear 5-min gradient ranging from 1% to 35% solvent B: acetonitrile, solvent A: 0.1% formic acid (FA) at a flow rate of 50 μl/min. All MS measurements were performed in triplicate. Peak areas were determined with Skyline 20.1.0.155 (MacCoss Lab, University of Washington)^52,75^.

The catalytic activity of a purified ROS-GC1 fragment (aa 814-1110) was performed by LC/MS/MS by an MRM approach using the specific transition of cGMP (*m/z* 346.0 → 152.0) (1200 HPLC system (Agilent Technologies) coupled to an LTQ-Orbitrap XL mass spectrometer (Thermo Fisher Scientific) equipped with nano-ESI source (Proxeon)^56,76^. The samples were separated via a C18 column (250 x 1 mm, 5 μm, 300 Å, Jupiter C18, Phenomenex) with a linear 10-min gradient from 2 % to 40 % solvent B: solvent B: acetonitrile with 0.08% formic acid (FA); solvent A: 0.1% formic acid (FA) at a flow rate of 50 μl/min. Peak areas were determined with Xcalibur (version 2.1; Thermo Fisher Scientific) in the ion chromatograms; linearity was obtained between 0.06-60 pmol cGMP.

### XL-MS Analyses

For XL-MS of ROS preparations, the IP beads loaded with ROS-GC1 were suspended in reaction buffer (20 mM HEPES, 100 mM NaCl, 2 mM EGTA, 2 mM MgCl_2_, 2 mM DDM, 10% (*v*/*v*) glycerol, pH 7.5). The cross-linking reaction (400 μl final volume) was induced by adding 1 mM DSBU^47,59^ (40 mM stock solution in DMSO). After incubating the mixture at room temperature for 60 min, the reaction was quenched with 20 mM Tris-HCl, pH 8.0. Alternatively, the 3D-structure of ROS-GC1 was stabilized by adding glutaraldehyde prior to IP and DSBU cross-linking. According to an established protocol^77^, 0.05 % (*v*/*v*) glutaraldehyde was added to 20 μl of ROS preparation and the solution was kept on ice for 5 min before the reaction was quenched by 100 mM Tris-HCl, pH 8.0 (final concentration).

For XL-MS of HEK293 cell lysates, cells (*see above*) of a confluent 75-cm^2^ flask were resuspended in 500 μl of buffer (20 mM HEPES, 100 mM NaCl, 1 mM TCEP, 0.1 mM EGTA, 2 mM MgCl_2_, 0.15 mM DDM, 10% (*v*/*v*) glycerol, cOmplete EDTA-free protease inhibitor cocktail (Roche), pH 7.5). After incubation on ice for 30 min, cells were lysed using a syringe (0.5 mm tube)^57,69^. The total protein concentration of the cell lysate was determined via Bradford protein assay to be 2 mg/ml. 0.5 μl of benzonase (stock solution 250 units/ml, Sigma Aldrich) were added and the cell lysate was incubated on ice for 30 min before 100 μl-aliquots were prepared. GCAP-2 was added to a final concentration of 1 μM to each aliquot. The cross-linker DSBU (40 mM stock solution in DMSO) or a 1:1 mixture of non-deuterated/deuterated cross-linker DSBU-D_0_/D_12_ (14 mM stock solution in DMSO) was added to a final concentration of 1 mM. The reaction mixtures were incubated at room temperature for 60 min. After quenching the reaction with 20 mM Tris-HCl, pH 8.0, samples were centrifuged at 13,000 x *g* for 20 min. The pellet, representing the HEK293 membrane fraction, was further used for LC/MS analysis^25,78^ (SEC and selected fractions for LC/MS/MS are shown in **Supplementary Fig. 13).**

For XL-MS of the purified ROS-GC1 domain (aa 814-1110), a protein concentration of 5 μM was employed in 50 μl reaction buffer (50 mM HEPES, 150 mM KCl, 1.5 mM MgCl_2_, 2.5 mM TCEP, 10% (v/v) glycerol, 0.05% DDM (*w/v*), 10 mM EGTA, 5 μM GCAP-2, pH 7.2). DSBU was added at 500-fold molar excess and the mixture was incubated at 0°C for 120 min, before the reaction was quenched with 20 mM NH_4_HCO_3_ (final concentration).

### SDS-PAGE and Enzymatic Digestion

For LC/MS/MS analysis of ROS-GC1 from bovine ROS preparations, *in-solution* digestion was performed directly on the magnetic beads used for IP (*see above*) with the filter-aided sample preparation (FASP) protocol^79^. The buffer was exchanged to 50 mM ammonium bicarbonate and samples were solubilized with 20 μl of 0,2% (*v*/*v*) ProteaseMAX surfactant solution. Tryptic digestion was performed according to the manufacturer’s protocol (Promega)^73,74^. After adding 2 μl of trypsin (0.5 μg/μl, Promega), peptide mixtures were incubated at 37°C for 4 hrs. Samples were acidified with 10 μl 10% (*v*/*v*) TFA. After digestion, supernatants from magnetic beads (~ 100 μl) were concentrated to 40 μl and analyzed by LC/MS/MS.

To monitor DSBU cross-linking of HEK293 cell lysates, SDS-PAGE analysis (10% gel) was performed. For *in-solution* digestion of HEK293 cell membranes, the same Protease MAX protocol was applied as for the ROS preparations. Peptide mixtures (~400 μl) were separated via SEC^59,80,81^ and concentrated to a volume of 50 μl each. For each cross-linking experiment, 12 SEC fractions were subjected to LC/MS/MS analysis.

Cross-linked samples of purified ROS-GC1 fragment (aa 814-1110) were analyzed by SDS-PAGE using gradient gels (4-20%, Mini-PROTEAN TGX gels, Biorad). For *in-gel* digestion, selected bands corresponding to ROS-GC1 monomer and dimer were excised. Enzymatic proteolysis with AspN (Promega) and trypsin as well as LC/MS/MS analysis were performed as previously described^82,83^.

### LC/MS/MS

Peptide mixtures originating from IP and SEC fractionation were analyzed by LC/MS/MS using an Ultimate 3000 RSLC nano HPLC system (Thermo Fisher Scientific) coupled to an Orbitrap Fusion Tribrid mass spectrometer equipped with Nanospray Flex ion source (Thermo Fisher Scientific). Peptides originating from *in-solution* digestion of ROS preparations and HEK293 cell membranes were concentrated and desalted on a C18 precolumn (Acclaim PepMap, 300 μM * 5 mm, 5 μm, 100 Å, Thermo Fisher Scientific) with 0.1% TFA at a flow rate of 30 μl/min for 15 minutes. Peptides from ROS preparations were separated by μPAC C18 (50 cm, PharmaFluidics) using a 5-min gradient from 3% to 10% solvent B (80% acetonitrile, 0.08% FA), 10% to 30% B (345 min), 30 to 85% B (5 min), and 85% B (5 min) at a flow rate of 0.3 μl/min.

LC separation of peptides originating from HEK293 cell membranes was performed via self-packed C18 columns (PicoFrits, 75 μm x 50 cm, 15 μm tip, New Objective, self-packed with ReproSil-Pur 120 C18-AQ, 1.9 μm, Dr. Maisch GmbH). A linear 175-min gradient was employed, ranging from 7% to 40% solvent B, followed by a 5-min gradient from 40% to 85% solvent B, and 85% solvent B for 5 min. Peptides originating from *in-gel* digestion of purified ROS-GC1 domain (aa 814-1110) were concentrated, desalted on a C8 column (Acclaim PepMap, 300 μm x 5 mm, 5 μm, 100 Å, Thermo Fisher Scientific), and separated on a C18 column (Acclaim PepMap, 75 μm * 250 mm, 2 μm, 100 Å, Thermo Fisher Scientific) using a 90-min LC gradient as described previously^84^.

For data acquisition in data-dependent MS/MS mode, the most intense signals in the MS full scan within 5 s were isolated (window 2 u) for fragmentation via stepped higher-energy collision-induced dissociation (HCD) using normalized collision energies of 30 ± 3 % (IP and SEC fractions) or 29 ± 5 % (purified ROS-GC1 domain). Dynamic exclusion was enabled (exclusion duration 60 s, deviation ± 2 ppm). For the ROS preparations and HEK293 cell membranes, inclusion lists comprising all theoretical masses and charge states of potential cross-linked products of bovine/human ROS-GC were generated to allow a targeted mass analysis. For the isotope-labeled cross-linker DSBU-D_0_/D_12_, the software ICC-CLASS (isotopically-coded cleavable crosslinking analysis software suite) served to extract D_0_/D_12_ signals^85^. Mass lists of DSBU-D_0_/D_12_ cross-linked products were generated and applied for a second LC/MS/MS analysis of corresponding SEC fractions^86^. Examples of annotated MS/MS spectra for cross-linked products from bovine and human full-length ROS-GC1 are shown in **Supplementary Figs. 14** and **15**, respectively.

### Analysis of Cross-linked Products

MS data were converted into mzML files by Proteome Discoverer 2.0 (Thermo Fisher Scientific) and the MeroX software, version 2.0 beta 5^59–61^, was used to assign cross-linked products. The RISEUP mode in MeroX was used allowing a maximum of three missing indicative ions. All cross-links were confirmed by manual validation. For DSBU-D_0_/D_12_ crosslinked peptides, the D_12_ composition of the cross-linker was added in MeroX.

### Molecular Modeling

According to our novel domain organization, ROS-GC1 IcD possesses three structural regions (**Fig. 1b**), for which suitable templates were identified with HHPRED^87^, namely KHD, αHD and CD. For predicting the 3D-structure of the KHD, templates were identified with HHPRED^87^ and a homology model was generated with MODELLER (Version 9.24) by default^88^. The KHD includes aa 542 – 815, with the exception of the highly flexible loop (aa 618 – 626), for which no suitable template was found. In addition, a loop (aa 688 – 709) was missing in the template (RIP1 kinase domain, 4NEU^89^), but was modeled using a homologous region from the kinase domain structure of PknA from *Mycobacterium tuberculosis* (4OW8)^90^. The dimeric model of the KHD domain was not captured by any template and was generated using the HADDOCK docking server^91^ with its grid implementation^92^. To model the KHD dimer, the resolved intermolecular cross-links were defined as 3 - 30 Å Cα-Cα distance range. Additionally, intermolecular cross-links to Lys-818 were considered by extending the distance range to 35 Å as the unresolved Lys-818 is located three amino acids further to the last resolved residue, Lys-815. Overall, 8 cross-link pairs were used as 16 unambiguous constraints (chain A to B and vice versa) and the guru interface was used to generate 5,000 models in it0, 400 models in it1, and 200 models during water refinement. In addition, C2 symmetry was applied. Statistics of docking are given in **Supplementary Table 9**. The 3D-structures of αHD (aa 823 – 859) and CD (aa 860-1066) were generated as dimers using the soluble guanylate cyclase domain as template (6PAS)^68^. We extended this model to cover the complete αHD (aa 801-859) by using the homologous sequence from the highest ranking template, identified with HHPRED (TAF5-TAF6-TAF9 complex^93^, 6F3T) utilizing MODELLER. To generate the model of the complete ROS-GC1 IcD, overlapping sequences (aa 801 – 816) from the αHD and KHD dimer were superimposed using PyMOL [https://pymol.org/] and optimized with MODELLER. The final ROS-GC1 IcD model was refined in explicit water using the HADDOCK refinement server^94–96^ by default.

## Abbreviations

αHD: α-helical domain
ACN: Acetonitrile
ATP: Adenosine triphosphate
CD: Catalytic domain
CCD: Core catalytic domain
cGMP: Cyclic guanosine monophosphate
CORD: Cone-rod dystrophies
CTE: *C*-terminal extension
DD: Dimerization domain
DDM: *n*-Dodecyl-β-D-maltopyranoside
DMEM: Dulbecco’s Modified Eagle’s Medium
DMSO: Dimethyl sulfoxide
DSBU: Disuccinimidyl dibutyric urea
DTT: Dithiothreitol
EDTA: 2,2’,2”,2ū-(Ethane-1,2-diyldinitrilo)tetraacetic acid
EGTA: Ethylene glycol-bis(2-aminoethylether)-*N,N,N’,N*-tetraacetic acid
ESI: Electrospray ionization
ExtD: Extracellular domain
FA: Formic acid
FASP: Filter-aided sample preparation
GC: Guanylyl cyclase
GCAP: Guanylyl cyclase-activating protein
GTP: Guanosine triphosphate
GMP: Guanosine monophosphate
HCD: Higher energy collision-induced dissociation
HEK: Human embryonic kidney
HEPES: 4-(2-Hydroxyethyl)-1-piperazineethanesulfonic acid
HLT: (His)_8_ tag, lipoyl domain, and tobacco etch virus cleavage site
IAA: Iodoacetamide
IBMX: 3-isobutyl-1-methylxanthine
IcD: Intracellular domain
IPTG: Isopropyl β-D-1-thiogalactopyranoside
JMD: Juxtamembrane domain
KHD: Kinase homology domain
LC: Liquid chromatography
LCA: Leber’s congenital amourosis
LS: Leader sequence
MRM: Multiple reaction monitoring
MS: Mass spectrometry
MS/MS: Tandem mass spectrometry
NPR: Natriuretic peptide receptor
NTA: Nitrilotriacetic acid
ROS: Rod-outer-segment
RP: Reversed phase
SCAD: Surfactant and chaotropic agent-assisted sequential extraction/on-pellet digestion
SDS-PAGE: Sodium dodecyl sulfate-polyacrylamide gel electrophoresis
SEC: Size exclusion chromatography
SHD: Signal helix domain
SRM: Selected reaction monitoring
STaR: Heat-stable enterotoxin receptor
TFA: Trifluoroacetic acid
Th: Thomson
TMD: Transmembrane domain
Tris: Tris(hydroxymethyl)aminomethane
XL-MS: Cross-linking/mass spectrometry

## Acknowledgements

A.S. acknowledges financial support by the DFG (project Si 867/15-2 and RTG 2467, project number 391498659 “Intrinsically Disordered Proteins – Molecular Principles, Cellular Functions, and Diseases”) and the region of Saxony-Anhalt. C.I. was funded by a postdoctoral fellowship of the Alexander-von-Humboldt foundation. P.L.K. acknowledges financial support by the BMBF (ZIK program, grant number 03Z22HN23), the European Regional Development Funds for Saxony-Anhalt (EFRE: ZS/2016/04/78115) and the Martin Luther University Halle-Wittenberg. The authors are indebted to Prof. Daniele Dell’Orco and Giuditta Dal Cortivo for providing ROS-GC1 transfected HEK293 cells and to Dr. Cordelia Schiene-Fischer for providing cell culture facilities.

## Author Contributions

A.R. and D.T. expressed and purified proteins and performed XL-MS experiments. A.R., A.S. and C.I. executed XL-MS data analysis. C.T. and P.L.K. performed computational modeling studies, C.H.I and M.K. executed MS analyses. K.W.K provided bovine ROS preparation samples and gave input to the manuscript. A.S. supervised the work. A.R., A.S., C.T. and P.L.K. wrote the manuscript. All authors approved the intellectual content.

## Supplementary Information

Supplementary Information is provided as a docx file.

## Data Availability

All data generated in this study have been made available. The mass spectrometry data have been deposited to the ProteomeXchange Consortium via the PRIDE partner repository with the dataset identifier PXD021656. Models will be deposited at https://data.sbgrid.org/.

## Materials & Correspondence

Request for materials or correspondence should be addressed to Prof. Dr. Andrea Sinz.

## References

1. Pugh, E.N., Jr. & Lamb, T.D. Phototransduction in vertebrate rods and cones: Molecular mechanisms of amplification, recovery and light adaptation. Handbook of Biological Physics 3, 183–255 (2000).

2. Pugh, E.N., Jr. & Cobbs, W.H. Visual transduction in vertebrate rods and cones: a tale of two transmitters, calcium and cyclic GMP. Vision Res 26, 1613–43 (1986).

3. Yau, K.W. Phototransduction mechanism in retinal rods and cones. The Friedenwald Lecture. Invest Ophthalmol Vis Sci 35, 9–32 (1994).

4. Koch, K.W. Purification and identification of photoreceptor guanylate cyclase. J Biol Chem 266, 8634–7 (1991).

5. Dizhoor, A.M., Lowe, D.G., Olshevskaya, E.V., Laura, R.P. & Hurley, J.B. The human photoreceptor membrane guanylyl cyclase, RetGC, is present in outer segments and is regulated by calcium and a soluble activator. Neuron 12, 1345–52 (1994).

6. Goraczniak, R.M., Duda, T., Sitaramayya, A. & Sharma, R.K. Structural and functional characterization of the rod outer segment membrane guanylate cyclase. Biochem J 302 (Pt 2), 455–61 (1994).

7. Dizhoor, A.M., Olshevskaya, E.V. & Peshenko, I.V. Mg2+/Ca2+ cation binding cycle of guanylyl cyclase activating proteins (GCAPs): role in regulation of photoreceptor guanylyl cyclase. Mol Cell Biochem 334, 117–24 (2010).

8. Koch, K.W. & Dell’Orco, D. Protein and Signaling Networks in Vertebrate Photoreceptor Cells. Front Mol Neurosci 8, 67 (2015).

9. Palczewski, K. et al. Molecular cloning and characterization of retinal photoreceptor guanylyl cyclase-activating protein. Neuron 13, 395–404 (1994).

10. Dizhoor, A.M. & Hurley, J.B. Regulation of photoreceptor membrane guanylyl cyclases by guanylyl cyclase activator proteins. Methods 19, 521–31 (1999).

11. Koch, K.W. & Stryer, L. Highly cooperative feedback control of retinal rod guanylate cyclase by calcium ions. Nature 334, 64–6 (1988).

12. Koch, K.W., Duda, T. & Sharma, R.K. Photoreceptor specific guanylate cyclases in vertebrate phototransduction. Mol Cell Biochem 230, 97–106 (2002).

13. Gray-Keller, M.P. & Detwiler, P.B. The calcium feedback signal in the phototransduction cascade of vertebrate rods. Neuron 13, 849–61 (1994).

14. Hayashi, F. & Yamazaki, A. Polymorphism in purified guanylate cyclase from vertebrate rod photoreceptors. Proc Natl Acad Sci U S A 88, 4746–50 (1991).

15. Margulis, A., Goraczniak, R.M., Duda, T., Sharma, R.K. & Sitaramayya, A. Structural and biochemical identity of retinal rod outer segment membrane guanylate cyclase. Biochem Biophys Res Commun 194, 855–61 (1993).

16. Shyjan, A.W., de Sauvage, F.J., Gillett, N.A., Goeddel, D.V. & Lowe, D.G. Molecular cloning of a retina-specific membrane guanylyl cyclase. Neuron 9, 727–37 (1992).

17. Sharma, R.K. & Duda, T. Membrane guanylate cyclase, a multimodal transduction machine: history, present, and future directions. Front Mol Neurosci 7, 56 (2014).

18. Lange, C., Duda, T., Beyermann, M., Sharma, R.K. & Koch, K.W. Regions in vertebrate photoreceptor guanylyl cyclase ROS-GC1 involved in Ca(2+)-dependent regulation by guanylyl cyclase-activating protein GCAP-1. FEBS Lett 460, 27–31 (1999).

19. Wilson, E.M. & Chinkers, M. Identification of sequences mediating guanylyl cyclase dimerization. Biochemistry 34, 4696–701 (1995).

20. Laura, R.P., Dizhoor, A.M. & Hurley, J.B. The membrane guanylyl cyclase, retinal guanylyl cyclase-1, is activated through its intracellular domain. J Biol Chem 271, 11646–51 (1996).

21. Yu, H. et al. Activation of retinal guanylyl cyclase-1 by Ca2+-binding proteins involves its dimerization. J Biol Chem 274, 15547–55 (1999).

22. Peshenko, I.V., Olshevskaya, E.V. & Dizhoor, A.M. Dimerization Domain of Retinal Membrane Guanylyl Cyclase 1 (RetGC1) Is an Essential Part of Guanylyl Cyclase-activating Protein (GCAP) Binding Interface. J Biol Chem 290, 19584–96 (2015).

23. Ramamurthy, V. et al. Interactions within the coiled-coil domain of RetGC-1 guanylyl cyclase are optimized for regulation rather than for high affinity. J Biol Chem 276, 26218–29 (2001).

24. Saha, S., Biswas, K.H., Kondapalli, C., Isloor, N. & Visweswariah, S.S. The linker region in receptor guanylyl cyclases is a key regulatory module: mutational analysis of guanylyl cyclase C. J Biol Chem 284, 27135–45 (2009).

25. Zägel, P., Dell’Orco, D. & Koch, K.W. The dimerization domain in outer segment guanylate cyclase is a Ca(2)(+)-sensitive control switch module. Biochemistry 52, 5065–74 (2013).

26. Anantharaman, V., Balaji, S. & Aravind, L. The signaling helix: a common functional theme in diverse signaling proteins. Biol Direct 1, 25 (2006).

27. Duda, T., Pertzev, A. & Sharma, R.K. Differential Ca(2+) sensor guanylate cyclase activating protein modes of photoreceptor rod outer segment membrane guanylate cyclase signaling. Biochemistry 51, 4650–7 (2012).

28. Duda, T., Pertzev, A., Makino, C.L. & Sharma, R.K. Bicarbonate and Ca(2+) Sensing Modulators Activate Photoreceptor ROS-GC1 Synergistically. Front Mol Neurosci 9, 5 (2016).

29. Duda, T. et al. Ca(2+) sensor S100beta-modulated sites of membrane guanylate cyclase in the photoreceptor-bipolar synapse. EMBO J 21, 2547–56 (2002).

30. Perrault, I. et al. Retinal-specific guanylate cyclase gene mutations in Leber’s congenital amaurosis. Nat Genet 14, 461–4 (1996).

31. Duda, T. & Koch, K.W. Retinal diseases linked with photoreceptor guanylate cyclase. Mol Cell Biochem 230, 129–38 (2002).

32. Sato, S., Peshenko, I.V., Olshevskaya, E.V., Kefalov, V.J. & Dizhoor, A.M. GUCY2D Cone-Rod Dystrophy-6 Is a “Phototransduction Disease” Triggered by Abnormal Calcium Feedback on Retinal Membrane Guanylyl Cyclase 1. J Neurosci 38, 2990–3000 (2018).

33. Sharon, D., Wimberg, H., Kinarty, Y. & Koch, K.W. Genotype-functional-phenotype correlations in photoreceptor guanylate cyclase (GC-E) encoded by GUCY2D. Prog Retin Eye Res 63, 69–91 (2018).

34. Duda, T. et al. Functional consequences of a rod outer segment membrane guanylate cyclase (ROS-GC1) gene mutation linked with Leber’s congenital amaurosis. Biochemistry 38, 509–15 (1999).

35. Wimberg, H. et al. Photoreceptor Guanylate Cyclase (GUCY2D) Mutations Cause Retinal Dystrophies by Severe Malfunction of Ca(2+)-Dependent Cyclic GMP Synthesis. Front Mol Neurosci 11, 348 (2018).

36. Sinz, A. Chemical cross-linking and mass spectrometry to map three-dimensional protein structures and protein-protein interactions. Mass Spectrom Rev 25, 663–82 (2006).

37. Rappsilber, J. The beginning of a beautiful friendship: cross-linking/mass spectrometry and modelling of proteins and multi-protein complexes. J Struct Biol 173, 530–40 (2011).

38. Piotrowski, C. & Sinz, A. Structural investigation of proteins and protein complexes by chemical cross-Linking/mass spectrometry, in Integrative Structural Biology with Hybrid Methods Vol. 1105 (eds Nakamura, H., Kleywegt, G., Burley, S.K. & Markley, J.L.) 101–121 (Advances in Experimental Medicine and Biology, 2018).

39. Liu, F., Rijkers, D.T., Post, H. & Heck, A.J. Proteome-wide profiling of protein assemblies by cross-linking mass spectrometry. Nat Methods 12, 1179–84 (2015).

40. Liu, F., Lossl, P., Scheltema, R., Viner, R. & Heck, A.J.R. Optimized fragmentation schemes and data analysis strategies for proteome-wide cross-link identification. Nat Commun 8, 15473 (2017).

41. Chavez, J.D. et al. Chemical Crosslinking Mass Spectrometry Analysis of Protein Conformations and Supercomplexes in Heart Tissue. Cell Syst 6, 136–141 e5 (2018).

42. Sinz, A. Crosslinking Mass Spectrometry Goes In-Tissue. Cell Syst 6, 10–12 (2018).

43. Politis, A. et al. A mass spectrometry-based hybrid method for structural modeling of protein complexes. Nat Methods 11, 403–406 (2014).

44. Leitner, A., Faini, M., Stengel, F. & Aebersold, R. Crosslinking and Mass Spectrometry: An Integrated Technology to Understand the Structure and Function of Molecular Machines. Trends Biochem Sci 41, 20–32 (2016).

45. Chavez, J.D. & Bruce, J.E. Chemical cross-linking with mass spectrometry: a tool for systems structural biology. Curr Opin Chem Biol 48, 8–18 (2019).

46. Kastritis, P.L. et al. Capturing protein communities by structural proteomics in a thermophilic eukaryote. Mol Syst Biol 13, 936 (2017).

47. Müller, M.Q., Dreiocker, F., Ihling, C.H., Schafer, M. & Sinz, A. Cleavable cross-linker for protein structure analysis: reliable identification of cross-linking products by tandem MS. Anal Chem 82, 6958–68 (2010).

48. Merkley, E.D. et al. Distance restraints from crosslinking mass spectrometry: mining a molecular dynamics simulation database to evaluate lysine-lysine distances. Protein Sci 23, 747–59 (2014).

49. Iacobucci, C. et al. A cross-linking/mass spectrometry workflow based on MS-cleavable cross-linkers and the MeroX software for studying protein structures and protein-protein interactions. Nat Protoc 13, 2864–2889 (2018).

50. Kettenbach, A.N., Rush, J. & Gerber, S.A. Absolute quantification of protein and post-translational modification abundance with stable isotope-labeled synthetic peptides. Nat Protoc 6, 175–86 (2011).

51. Yim, Y.Y. et al. Quantitative Multiple-Reaction Monitoring Proteomic Analysis of Gbeta and Ggamma Subunits in C57Bl6/J Brain Synaptosomes. Biochemistry 56, 5405–5416 (2017).

52. MacLean, B. et al. Skyline: an open source document editor for creating and analyzing targeted proteomics experiments. Bioinformatics 26, 966–8 (2010).

53. Pino, L.K. et al. The Skyline ecosystem: Informatics for quantitative mass spectrometry proteomics. Mass Spectrom Rev 39, 229–244 (2020).

54. Schilling, B. et al. Platform-independent and label-free quantitation of proteomic data using MS1 extracted ion chromatograms in skyline: application to protein acetylation and phosphorylation. Mol Cell Proteomics 11, 202–14 (2012).

55. Hwang, J.Y. et al. Regulatory modes of rod outer segment membrane guanylate cyclase differ in catalytic efficiency and Ca(2+)-sensitivity. Eur J Biochem 270, 3814–21 (2003).

56. Lange, V., Picotti, P., Domon, B. & Aebersold, R. Selected reaction monitoring for quantitative proteomics: a tutorial. Mol Syst Biol 4, 222 (2008).

57. Dell’Orco, D. & Dal Cortivo, G. Normal GCAPs partly compensate for altered cGMP signaling in retinal dystrophies associated with mutations in GUCA1A. Sci Rep 9, 20105 (2019).

58. Koch, K.W. & Helten, A. Guanylate cyclase-based signaling in photoreceptors and retina, in Signal Transduction in the Retina (eds. Fliesler, S.J. & Kisselev, O.G.) 121–143 (CRC Press, 2008).

59. Götze, M., Iacobucci, C., Ihling, C.H. & Sinz, A. A Simple Cross-Linking/Mass Spectrometry Workflow for Studying System-wide Protein Interactions. Anal Chem 91, 10236–10244 (2019).

60. Götze, M. et al. Automated assignment of MS/MS cleavable cross-links in protein 3Dstructure analysis. J Am Soc Mass Spectrom 26, 83–97 (2015).

61. Götze, M. et al. StavroX--a software for analyzing crosslinked products in protein interaction studies. J Am Soc Mass Spectrom 23, 76–87 (2012).

62. Ravichandran, S., Duda, T., Pertzev, A. & Sharma, R.K. Membrane Guanylate Cyclase catalytic Subdomain: Structure and Linkage with Calcium Sensors and Bicarbonate. Front Mol Neurosci 10, 173 (2017).

63. Venkataraman, V., Duda, T., Ravichandran, S. & Sharma, R.K. Neurocalcin delta modulation of ROS-GC1, a new model of Ca(2+) signaling. Biochemistry 47, 6590–601 (2008).

64. Park, E. et al. Architecture of autoinhibited and active BRAF-MEK1-14-3-3 complexes. Nature 575, 545–550 (2019).

65. Wagner, T. et al. The crystal structure of the catalytic domain of the ser/thr kinase PknA from M. tuberculosis shows an Src-like autoinhibited conformation. Proteins 83, 982–8 (2015).

66. Lorenz, S., Deng, P., Hantschel, O., Superti-Furga, G. & Kuriyan, J. Crystal structure of an SH2-kinase construct of c-Abl and effect of the SH2 domain on kinase activity. Biochem J 468, 283–91 (2015).

67. Wang, J. et al. Reconstitution and structure of a plant NLR resistosome conferring immunity. Science 364(2019).

68. Horst, B.G. et al. Allosteric activation of the nitric oxide receptor soluble guanylate cyclase mapped by cryo-electron microscopy. Elife 8 (2019).

69. Wimberg, H., Janssen-Bienhold, U. & Koch, K.W. Control of the Nucleotide Cycle in Photoreceptor Cell Extracts by Retinal Degeneration Protein 3. Front Mol Neurosci 11, 52 (2018).

70. Sulmann, S., Kussrow, A., Bornhop, D.J. & Koch, K.W. Label-free quantification of calcium-sensor targeting to photoreceptor guanylate cyclase and rhodopsin kinase by backscattering interferometry. Sci Rep 7, 45515 (2017).

71. Schnetkamp, P.P. & Daemen, F.J. Isolation and characterization of osmotically sealed bovine rod outer segments. Methods Enzymol 81, 110–6 (1982).

72. Koch, K.W., Lambrecht, H.G., Haberecht, M., Redburn, D. & Schmidt, H.H. Functional coupling of a Ca2+/calmodulin-dependent nitric oxide synthase and a soluble guanylyl cyclase in vertebrate photoreceptor cells. EMBO J 13, 3312–20 (1994).

73. Saveliev, S.V. et al. Mass spectrometry compatible surfactant for optimized in-gel protein digestion. Anal Chem 85, 907–14 (2013).

74. Pirmoradian, M. et al. Rapid and deep human proteome analysis by single-dimension shotgun proteomics. Mol Cell Proteomics 12, 3330–8 (2013).

75. Adams, K.J. et al. Skyline for Small Molecules: A Unifying Software Package for Quantitative Metabolomics. J Proteome Res 19, 1447–1458 (2020).

76. Calvo, E., Camafeita, E., Fernandez-Gutierrez, B. & Lopez, J.A. Applying selected reaction monitoring to targeted proteomics. Expert Rev Proteomics 8, 165–73 (2011).

77. Subbotin, R.I. & Chait, B.T. A pipeline for determining protein-protein interactions and proximities in the cellular milieu. Mol Cell Proteomics 13, 2824–35 (2014).

78. Zägel, P. & Koch, K.W. Dysfunction of outer segment guanylate cyclase caused by retinal disease related mutations. Front Mol Neurosci 7, 4 (2014).

79. Wisniewski, J.R., Zougman, A., Nagaraj, N. & Mann, M. Universal sample preparation method for proteome analysis. Nat Methods 6, 359–62 (2009).

80. Rampler, E. et al. Comprehensive Cross-Linking Mass Spectrometry Reveals Parallel Orientation and Flexible Conformations of Plant HOP2-MND1. J Proteome Res 14, 5048–62 (2015).

81. Herzog, F. et al. Structural probing of a protein phosphatase 2A network by chemical crosslinking and mass spectrometry. Science 337, 1348–52 (2012).

82. Rehkamp, A. et al. Molecular Details of Retinal Guanylyl Cyclase 1/GCAP-2 Interaction. Front Mol Neurosci 11, 330 (2018).

83. Lossl, P. & Sinz, A. Combining Amine-Reactive Cross-Linkers and Photo-Reactive Amino Acids for 3D-Structure Analysis of Proteins and Protein Complexes. Methods Mol Biol 1394, 109–127 (2016).

84. Iacobucci, C., Piotrowski, C., Rehkamp, A., Ihling, C.H. & Sinz, A. The First MS-Cleavable, Photo-Thiol-Reactive Cross-Linker for Protein Structural Studies. J Am Soc Mass Spectrom 30, 139–148 (2019).

85. Petrotchenko, E.V. & Borchers, C.H. ICC-CLASS: isotopically-coded cleavable crosslinking analysis software suite. BMC Bioinformatics 11, 64 (2010).

86. Petrotchenko, E.V., Makepeace, K.A., Serpa, J.J. & Borchers, C.H. Analysis of protein structure by cross-linking combined with mass spectrometry. Methods Mol Biol 1156, 447–63 (2014).

87. Zimmermann, L. et al. A Completely Reimplemented MPI Bioinformatics Toolkit with a New HHpred Server at its Core. J Mol Biol 430, 2237–2243 (2018).

88. Webb, B. & Sali, A. Comparative Protein Structure Modeling Using MODELLER. Curr Protoc Bioinformatics 54, 5 6 1-5 6 37 (2016).

89. Harris, P.A. et al. Discovery of Small Molecule RIP1 Kinase Inhibitors for the Treatment of Pathologies Associated with Necroptosis. ACS Med Chem Lett 4, 1238–43 (2013).

90. Ravala, S.K. et al. Evidence that phosphorylation of threonine in the GT motif triggers activation of PknA, a eukaryotic-type serine/threonine kinase from Mycobacterium tuberculosis. FEBS J 282, 1419–31 (2015).

91. van Zundert, G.C.P. et al. The HADDOCK2.2 Web Server: User-Friendly Integrative Modeling of Biomolecular Complexes. J Mol Biol 428, 720–725 (2016).

92. Wassenaar, T.A. et al. WeNMR: Structural biology on the grid. Journal of Grid Computing 10, 743–767 (2012).

93. Antonova, S.V. et al. Chaperonin CCT checkpoint function in basal transcription factor TFIID assembly. Nat Struct Mol Biol 25, 1119–1127 (2018).

94. Kastritis, P.L. & Bonvin, A.M. Are scoring functions in protein-protein docking ready to predict interactomes? Clues from a novel binding affinity benchmark. J Proteome Res 9, 2216–25 (2010).

95. Karaca, E., Melquiond, A.S., de Vries, S.J., Kastritis, P.L. & Bonvin, A.M. Building macromolecular assemblies by information-driven docking: introducing the HADDOCK multibody docking server. Mol Cell Proteomics 9, 1784–94 (2010).

96. Kastritis, P.L., Rodrigues, J.P., Folkers, G.E., Boelens, R. & Bonvin, A.M. Proteins feel more than they see: fine-tuning of binding affinity by properties of the non-interacting surface. J Mol Biol 426, 2632–52 (2014).

