## Supplementary Information for "First 3D-Structural Data of Full-length Rod-Outer-Segment Guanylyl Cyclase 1 in Bovine Retina by Cross-linking/Mass Spectrometry"

^*^Address correspondence to:

**Table of Content**

Optimized Digestion Protocol for ROS-GC1

Supplementary Fig. 1. Absolute Quantification of Bovine ROS-GC1 in ROS Preparation

Supplementary Fig. 2. Calibration Curves for Determining ROS-GC1 Activity

Supplementary Fig. 3. Catalytic Activity of Human ROS-GC1 in the Absence of Calcium

Supplementary Fig. 4. Background GTP Metabolism in ROS Preparation

Supplementary Fig. 5. Fragmentation of the MS-Cleavable Cross-linker DSBU

Supplementary Fig. 6. SDS-PAGE Analysis of Cross-linked HEK293 Cell Lysate

Supplementary Fig. 7. DSBU Cross-links in Human ROS-GC1 Dimer

Supplementary Fig. 8. SDS-PAGE Analysis of Cross-linking Reactions between Bovine ROS-GC1 Fragment (aa 814-1110) and GCAP-2

Supplementary Fig. 9. DSBU Cross-links Identified in Bovine ROS-GC1 Fragment (aa 814-1110)

Supplementary Fig. 10. MS/MS Spectra of Cross-links of Bovine ROS-GC1 Fragment (aa 814-1110) Dimer

Supplementary Fig. 11. Comparison of DSBU Cross-links Identified in Bovine (a) and Human (b) ROS-GC1 Dimer

Supplementary Fig. 12. Exploring Torsion Angle Space of αHD and *C*-Terminus of CD 🡪 12

Supplementary Fig. 13. Size Exclusion Chromatogram of Peptide Mixture from HEK293 Cell Lysate after Cross-linking with DSBU and In-solution Digestion

Supplementary Fig. 14. MS/MS Spectra of Cross-linked Products from Bovine, Full-length ROS-GC1

Supplementary Fig. 15. MS/MS Spectra of Cross-linked Products from Human, Full-length ROS-GC1

Supplementary Table 1. DSBU Cross-links in Bovine, Full-length ROS-GC1 in ROS-Preparation

Supplementary Table 2. DSBU Cross-links of Bovine, Full-length ROS-GC1 in ROS-Preparation after Glutaraldehyde Fixiation

Supplementary Table 3. DSBU Cross-links in Human, Full-length ROS-GC1

Supplementary Table 4. DSBU Cross-links in Human, Full-length ROS-GC1

Supplementary Table 5. DSBU-D_0_/D_12_ Cross-links in Human, Full-length ROS-GC1

Supplementary Table 6. Unique DSBU Cross-linking Sites in Human, Full-length ROS-GC1

Supplementary Table 7. Cross-linking Sites in Bovine ROS-GC1 Fragment (aa 814-1110)

Supplementary Table 8. DSBU Cross-links in Bovine ROS-GC1 Fragment (aa 814-1110)

Supplementary Table 9. HADDOCK Statistics for C2 Binary Docking of KHD

Supplementary Table 10. Cross-link Distances in Bovine ROS-GC1 (IcD Model)

Supplementary Table 11. Cross-link Distances in Human ROS-GC1 (IcD Model)

Supplementary Table 12. Cross-link distances in Bovine ROS-GC1 Fragment (aa 814-1110, IcD model)

*Optimized Digestion Protocol for ROS-GC1*

The digestion protocol for ROS-GC1 was optimized in HEK293 cell lysates and then applied to bovine ROS-GC1 from ROS preparation. In summary, we compared three different digestion protocols, filter-aided sample preparation (FASP)^1^, surfactant and chaotropic agent-assisted sequential extraction/on-pellet digestion (SCAD)^2^, and the ProteaseMaX workflow^3,4^. Highly reproducible sequence coverage of ~40 to 50% was obtained for human ROS-GC1 derived from the HEK cell lysate by applying the ProteaseMaX protocol. For bovine ROS-GC1 from ROS preparation a combination of FASP and ProteaseMaX protocols proved to yield maximum sequence coverage (see *Methods*). An additional SEC enrichment step of cross-linked peptides greatly reduced sample complexity and allowed identifying even low-abundant human ROS-GC1 cross-links from HEK293 membrane preparations (**Supplementary Fig. 13**)^5^. In-depth LC/MS/MS analysis delivered a total of 30 unique cross-linking sites from full-length, bovine ROS-GC1 from ROS preparation. For full-length, human ROS-GC1 overexpressed in HEK293 cells, 27 cross-links were obtained. Two exemplary MS/MS spectra allowing the unambiguous identification of cross-linked amino acids are presented in **Supplementary Fig. 14** (bovine ROS-GC1) and **Supplementary Fig. 15** (human ROS-GC1).

*Supplementary References*

1.	Wisniewski, J.R., Zougman, A., Nagaraj, N. & Mann, M. Universal sample preparation method for proteome analysis. *Nat Methods* **6**, 359-62 (2009).

2.	Ravichandran, S., Duda, T., Pertzev, A. & Sharma, R.K. Membrane Guanylate Cyclase catalytic Subdomain: Structure and Linkage with Calcium Sensors and Bicarbonate. *Front Mol Neurosci* **10**, 173 (2017).

3.	Pirmoradian, M. et al. Rapid and deep human proteome analysis by single-dimension shotgun proteomics. *Mol Cell Proteomics* **12**, 3330-8 (2013).

4.	Saveliev, S.V. et al. Mass spectrometry compatible surfactant for optimized in-gel protein digestion. *Anal Chem* **85**, 907-14 (2013).

5.	Götze, M., Iacobucci, C., Ihling, C.H. & Sinz, A. A Simple Cross-Linking/Mass Spectrometry Workflow for Studying System-wide Protein Interactions. *Anal Chem* **91**, 10236-10244 (2019).

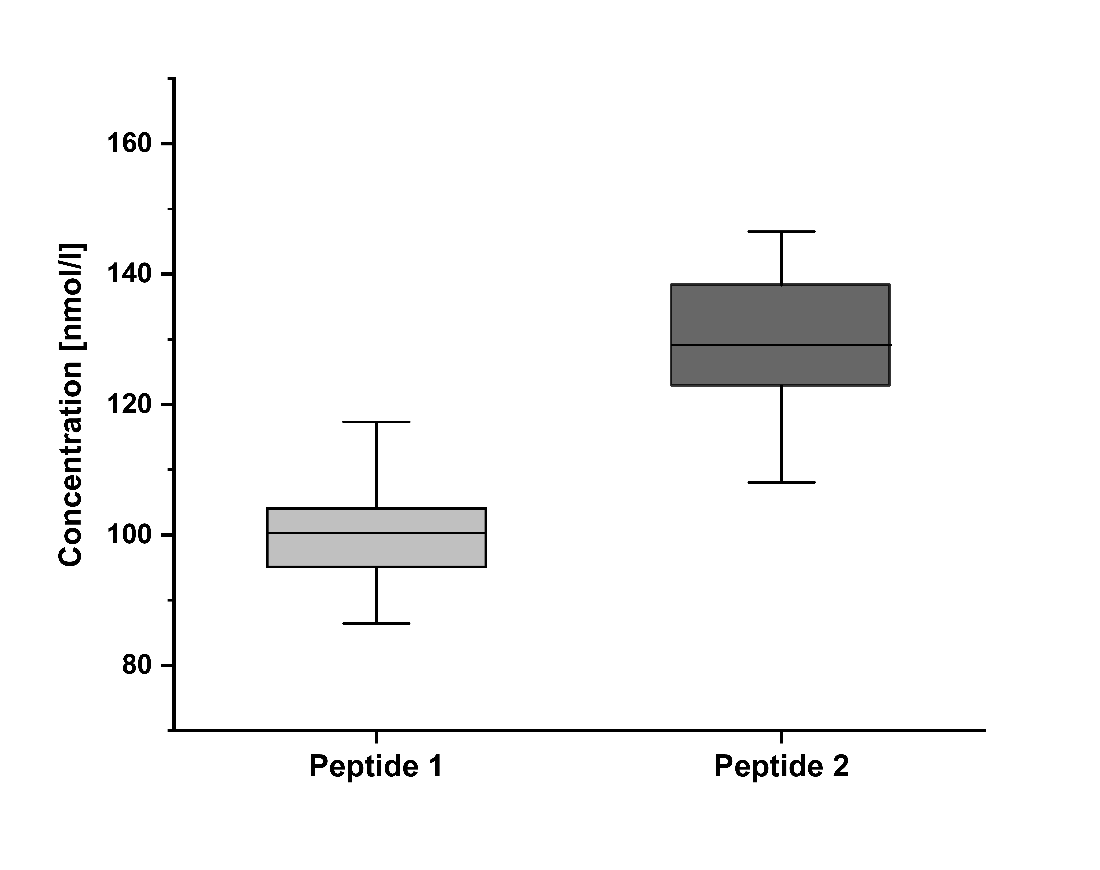

Supplementary Fig. 1. Absolute Quantification of Bovine ROS-GC1 in ROS Preparation. Peptides with isotope-labeled, C-terminal Arg (peptide 1 _162_AAGTTAPVVTPAADALYALLR_182_ and peptide 2 _1028_STVQILSALNEGFLTEVR_1045_) were used for absolute quantification of ROS-GC1 in the ROS preparation. For nomenclature of ROS-GC1 peptides, please see Fig. 1. Intensity ratios were determined between the peptide that had been generated during ROS-GC1 digestion and the externally added, isotope-labeled peptide. Data analysis was performed with Skyline 20.1.0.155. ROS-GC1 concentrations were determined to be 100.3 ± 5.8 nmol/l (based on peptide 1) and 129.1 ±5.0 nmol/l (based on peptide 2).

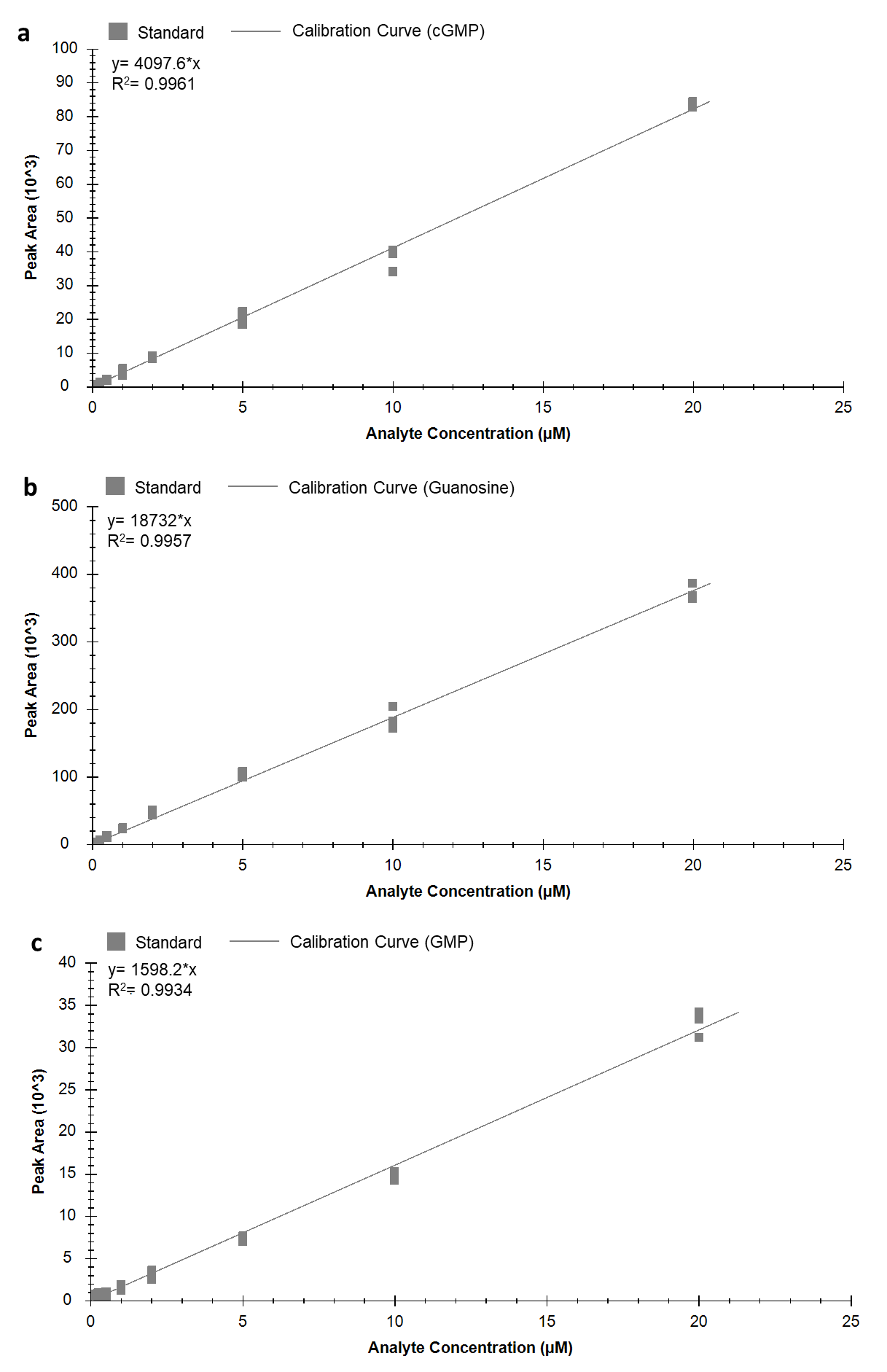

Supplementary Fig. 2. Calibration Curves for Determining ROS-GC1 Activity. a) cGMP, b) guanosine, c) GMP. A linear response was obtained for concentrations between 0.1 µM to 20 µM.

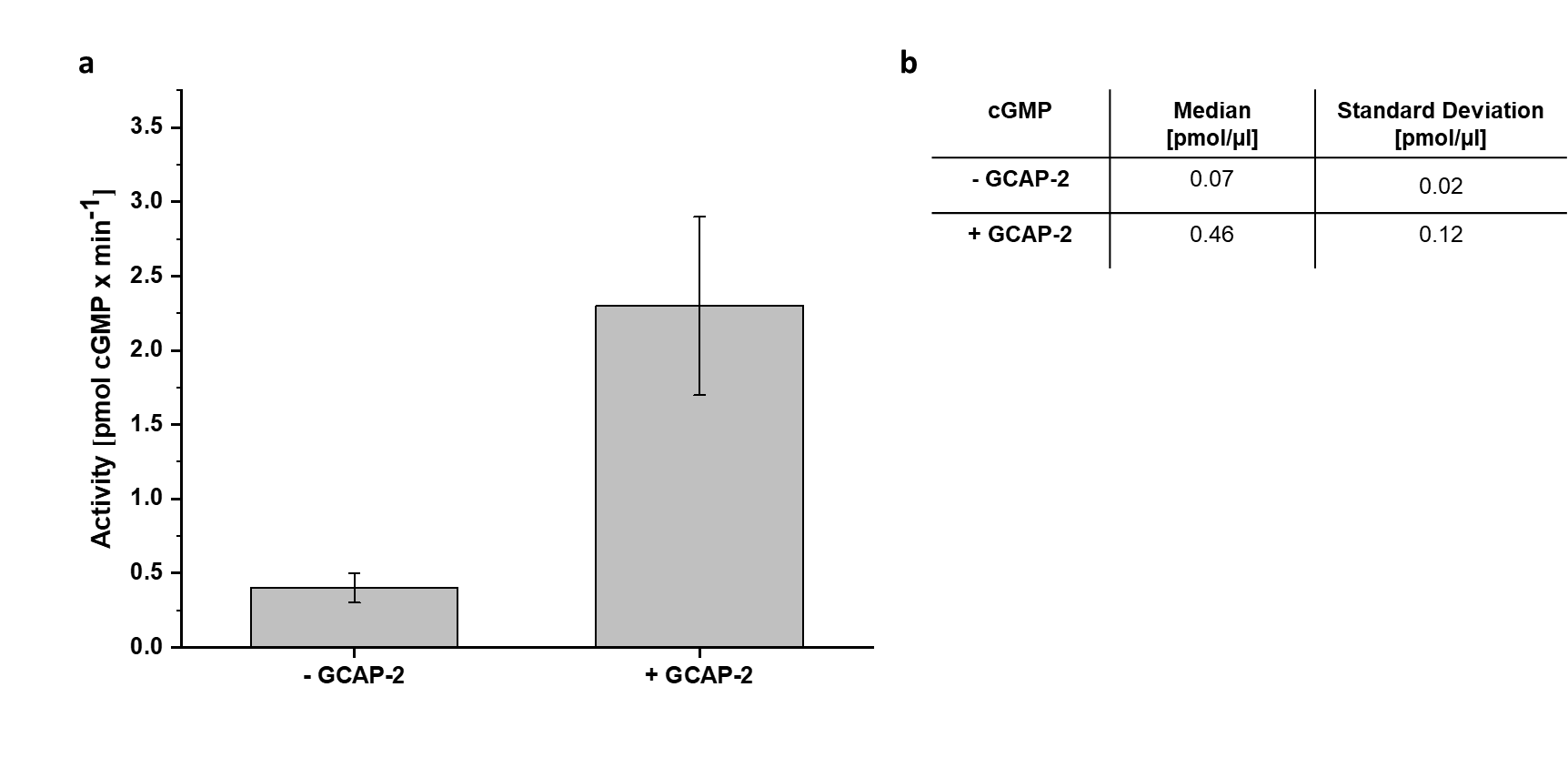

**Supplementary Fig. 3.** **Catalytic Activity of Human ROS-GC1 in the Absence of Calcium**. **a**) ROS-GC1 activity; **b**) cGMP concentration (- and + GCAP-2). Human ROS-GC1 from HEK 293 cell membranes was incubated for 10 minutes in the presence (1 µM) and absence of GCAP-2. Quantification of cGMP, GMP, and guanosine was performed by mass spectrometric MRM analysis, peak areas were determined with Skyline 20.1.0.155. ROS-GC1 activity was determined to be 0.4 ± 0.1 pmol/min (- GCAP-2) and 2.3 ± 0.6 pmol/min (+ GCAP-2) based on cGMP concentration; no residual enzymatic activity was detectable in control experiments.

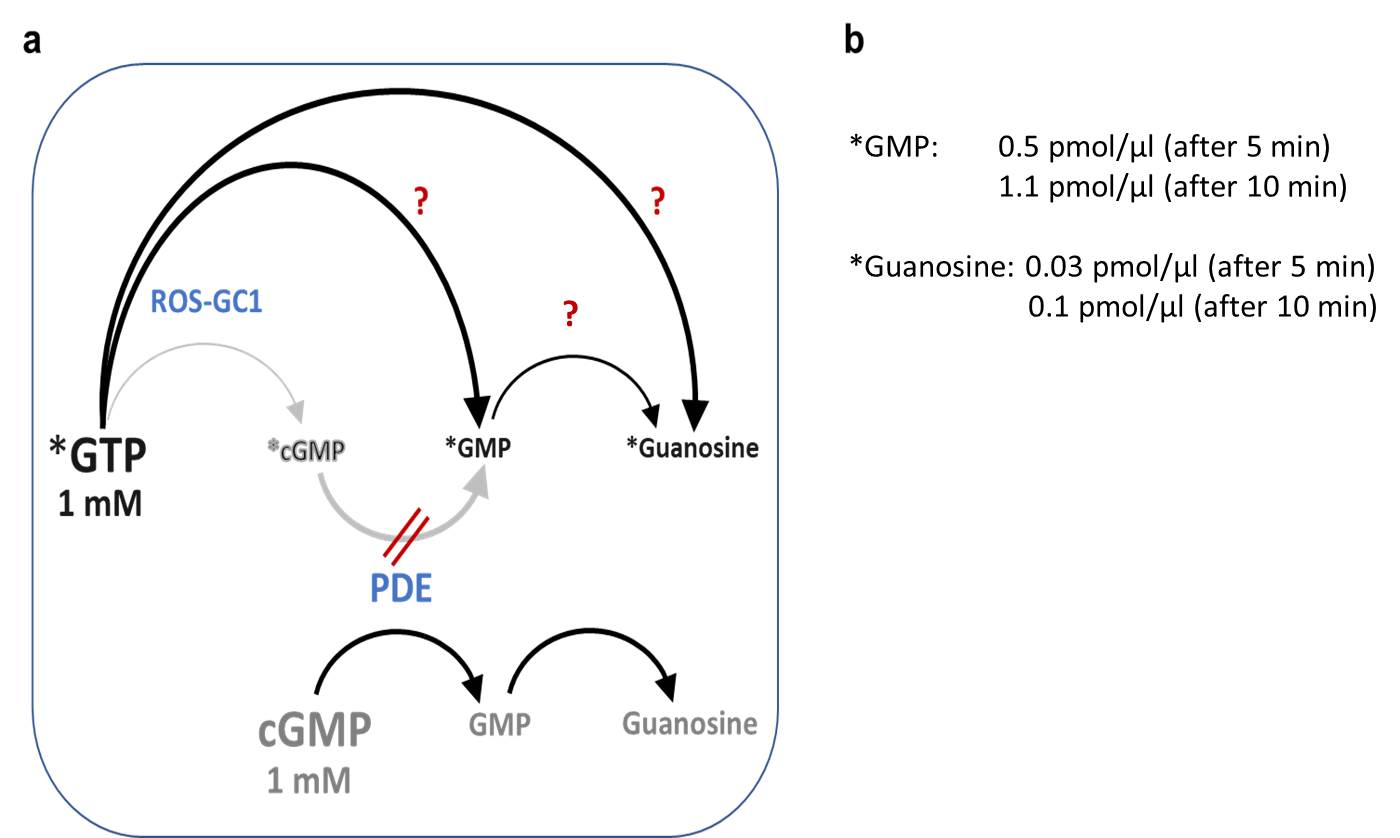

Supplementary Fig. 4. Background GTP Metabolism in ROS Preparation. a) Overview of GTP conversion reactions; PDE: phosphodiesterases; isotope-labeled species (^13^C and ^15^N) are marked with an asterisk; b) Background levels of *GMP and *Guanosine. *GMP and *Guanosine levels were determined in the presence of 1 mM *GTP and 1 mM cGMP. An excess of 1 mM cGMP competes with *cGMP conversion at the PDE, so *cGMP will not be converted to *GMP. Therefore, *GMP and *Guanosine can only be generated from *GTP by ROS-GC1-independent processes (marked with ?). These background levels were subtracted from GMP and Guanosine levels in all experiments without labeling and PDE competition. Conducting experiments in this manner was required as PDE activity could not be completely inhibited. We calculated cGMP levels that are exclusively generated by active ROS-GC1 by adding the GMP and Guanosine levels and subtracting background levels from isotope-labeled *GMP and *Guanosine. All levels of GTP metabolites were determined by MRM mass spectrometry.

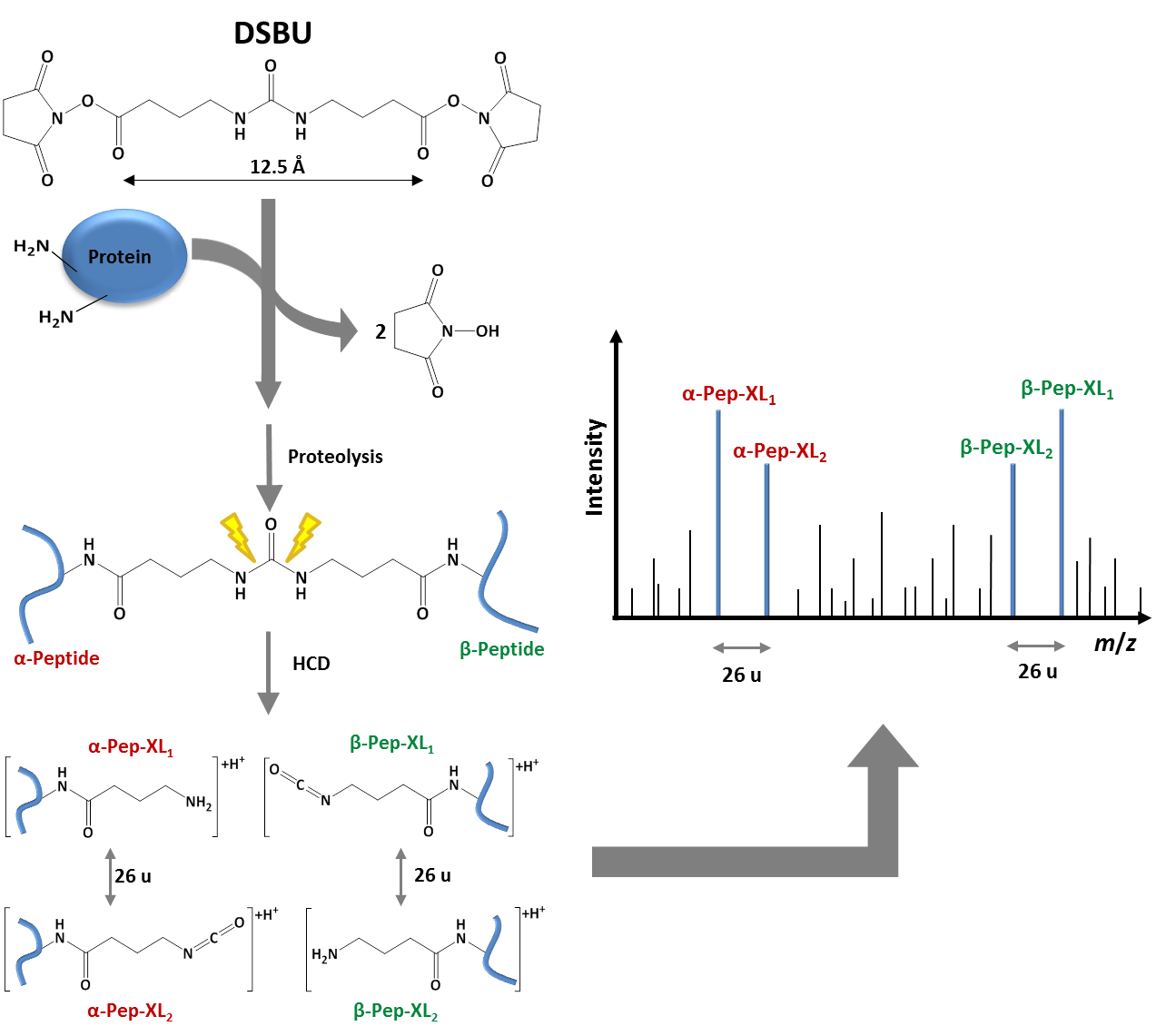

Supplementary Fig. 5. Fragmentation of the MS-Cleavable Cross-linker DSBU. DSBU reacts preferentially with lysine residues in proteins. After enzymatic proteolysis, the urea group of DSBU is fragmented by collisional activation in the mass spectrometer (HCD: higher collision-induced dissociation); fragmentation sites are indicated in yellow. Fragments are generated with a characteristic mass difference of ~ 26 u facilitating the automated identification of cross-linked products by customized software.

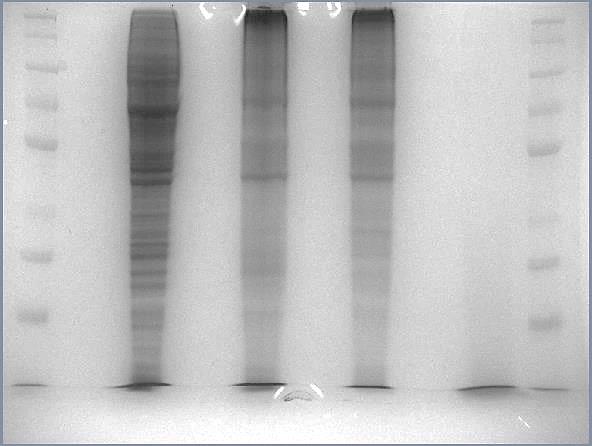

**+ DSBU**

**40 kDa**

**50 kDa**

**70 kDa**

**100 kDa**

**140 kDa**

**260 kDa**

**35 kDa**

**25 kDa**

**- DSBU**

**Supplementary Fig. 6. SDS-PAGE Analysis of Cross-linked HEK293 Cell Lysate.** Cross-linking was performed in the presence or absence (+/- DSBU). Coomassie staining was applied.

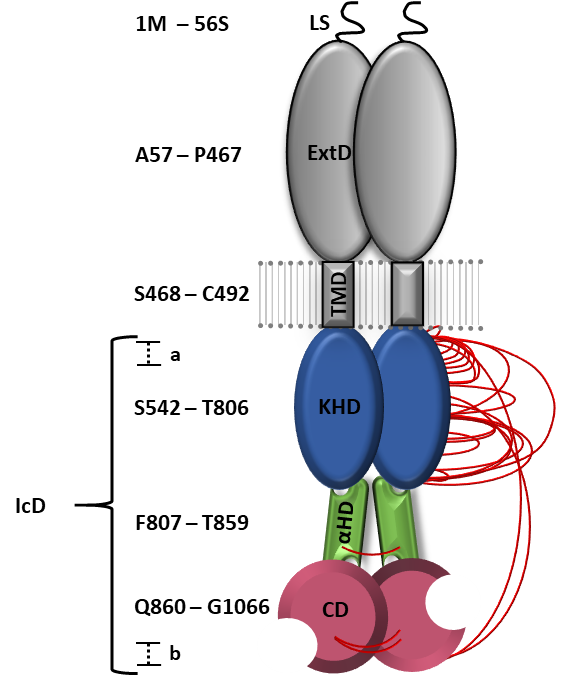

Supplementary Fig. 7. DSBU Cross-links in Human ROS-GC1 Dimer. The domain organization of ROS-GC1 was revised based on the XL-MS and computational modeling experiments described herein. Cross-links are presented in red.

**M**

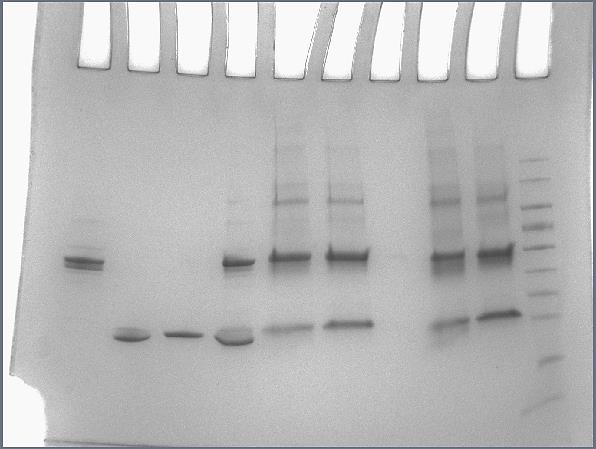

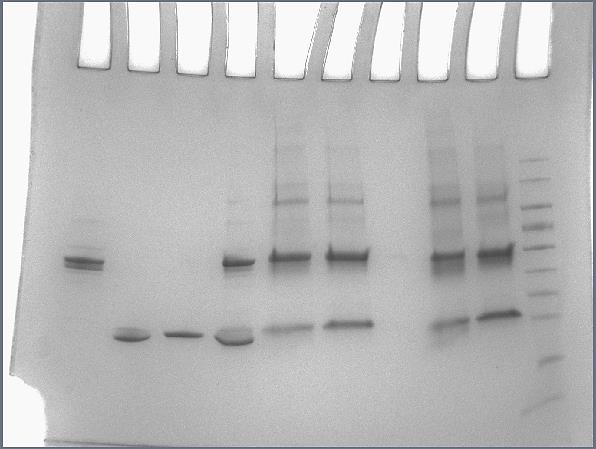

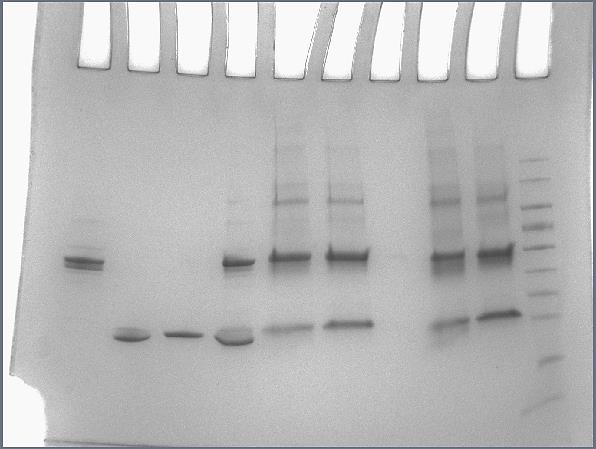

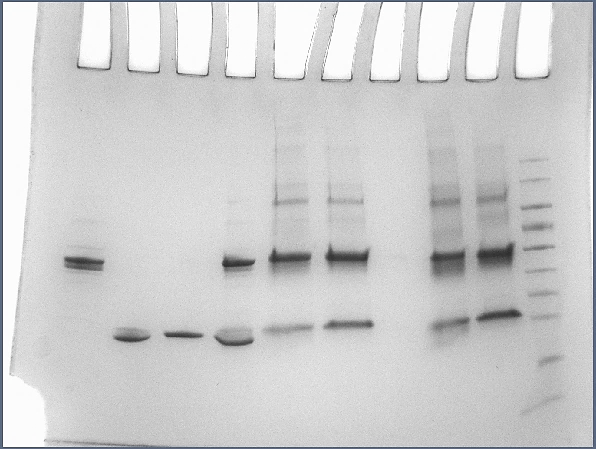

**25 kDa**

**15 kDa**

**35 kDa**

**40 kDa**

**55 kDa**

**70 kDa**

**100 kDa**

**130 kDa**

**170 kDa**

**MW**

**D**

**1**

**2**

**3**

**Supplementary Fig. 8.** **SDS-PAGE Analysis of Cross-linking Reactions between Bovine ROS-GC1 Fragment (aa 814-1110) and GCAP-2.** *Lane 1*: ROS-GC1 fragment without cross-linker, *lane 2*: GCAP-2 without cross-linker, *lane 3*: DSBU cross-linking reaction mixture between ROS-GC fragment and GCAP-2. MW: molecular weight marker, M: monomer , D: dimer of ROS-GC fragment. For *in-gel* digestion, the boxed gel bands were excised and subjected to LC/MS/MS analysis; Coomassie staining was applied.

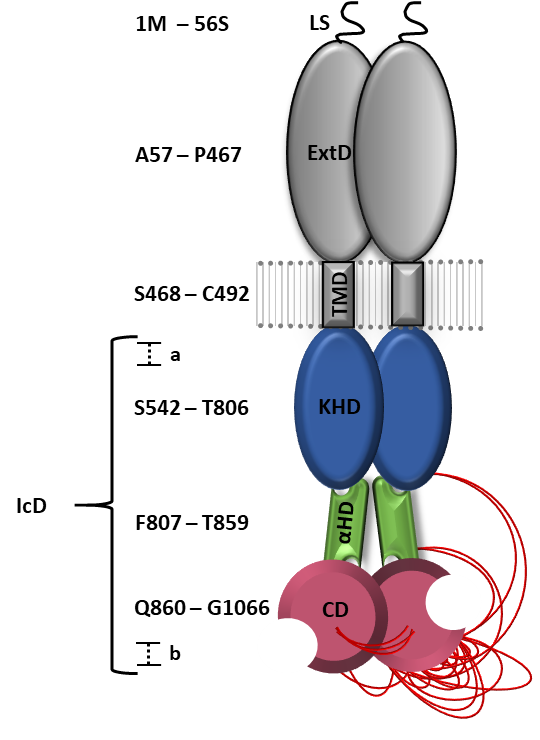

Supplementary Fig. 9. DSBU Cross-links Identified in Bovine ROS-GC1 Fragment (aa 814-1110). Cross-links were identified in the gel band (Supplementary Fig. 8) of the dimer. The domain organization of ROS-GC1 was revised based on the XL-MS and computational modeling experiments described herein. Cross-links are presented in red.

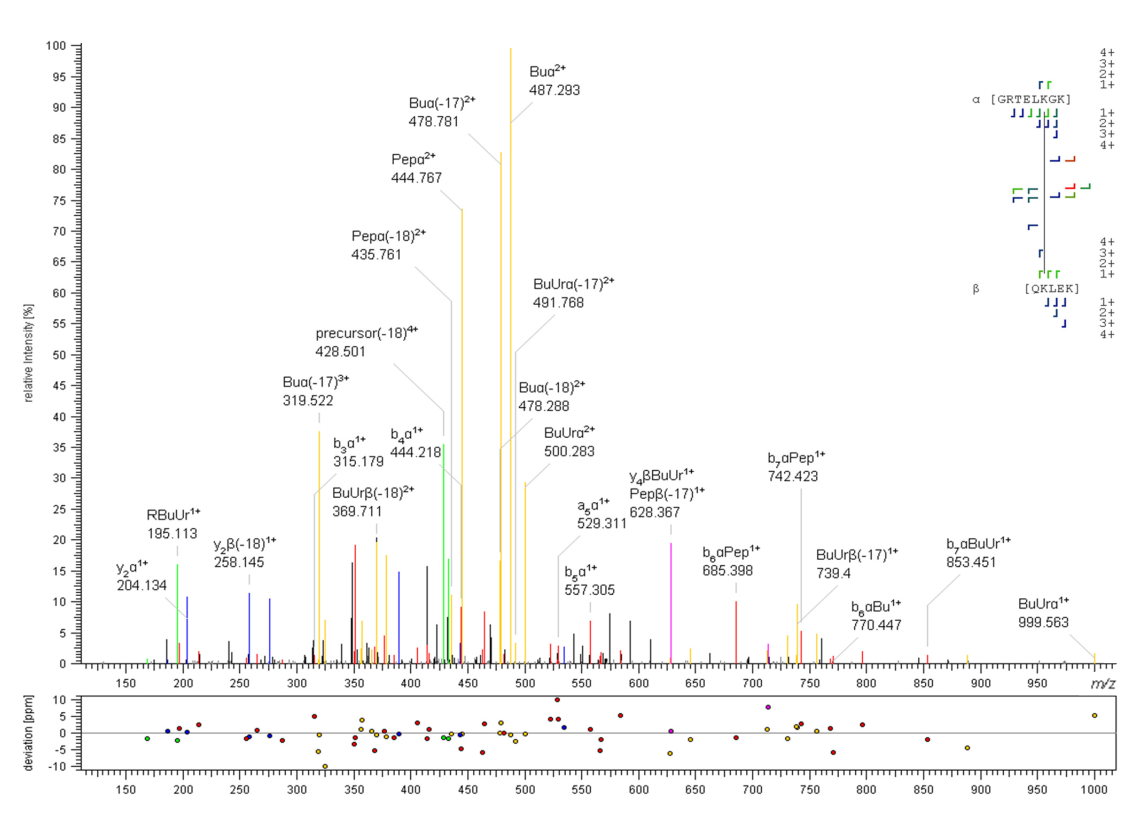

**b**

**a**

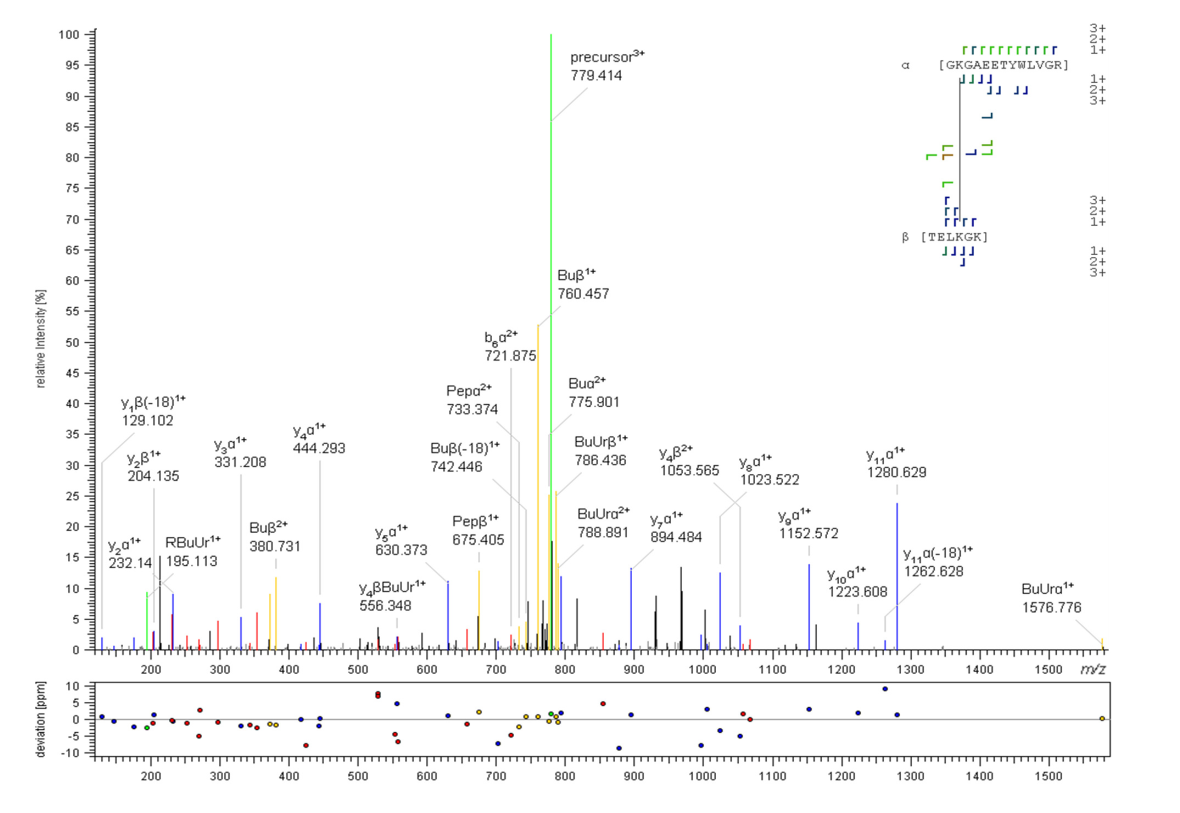

Supplementary Fig. 10. MS/MS Spectra of Cross-links of Bovine ROS-GC1 Fragment (aa 814-1110) Dimer. HCD-MS/MS spectra were automatically annotated with MeroX; a) A cross-link (4+ charged precursor ion at *m*/*z* 433.005) was identified between K1098 (aa 1097-1101) and K1051 (aa 1046-1053); b) a cross-link (4+ charged precursor ion at *m*/*z* 779.414) between K1053 (aa 1052-1064) and K1051 (aa 1048-1053) indicates an interprotein cross-link between two ROS-GC1 monomers. Precursor ions are shown in green, fragment ions of DSBU are shown in yellow, b- and y-type ions of the peptide backbone are shown in red and blue.

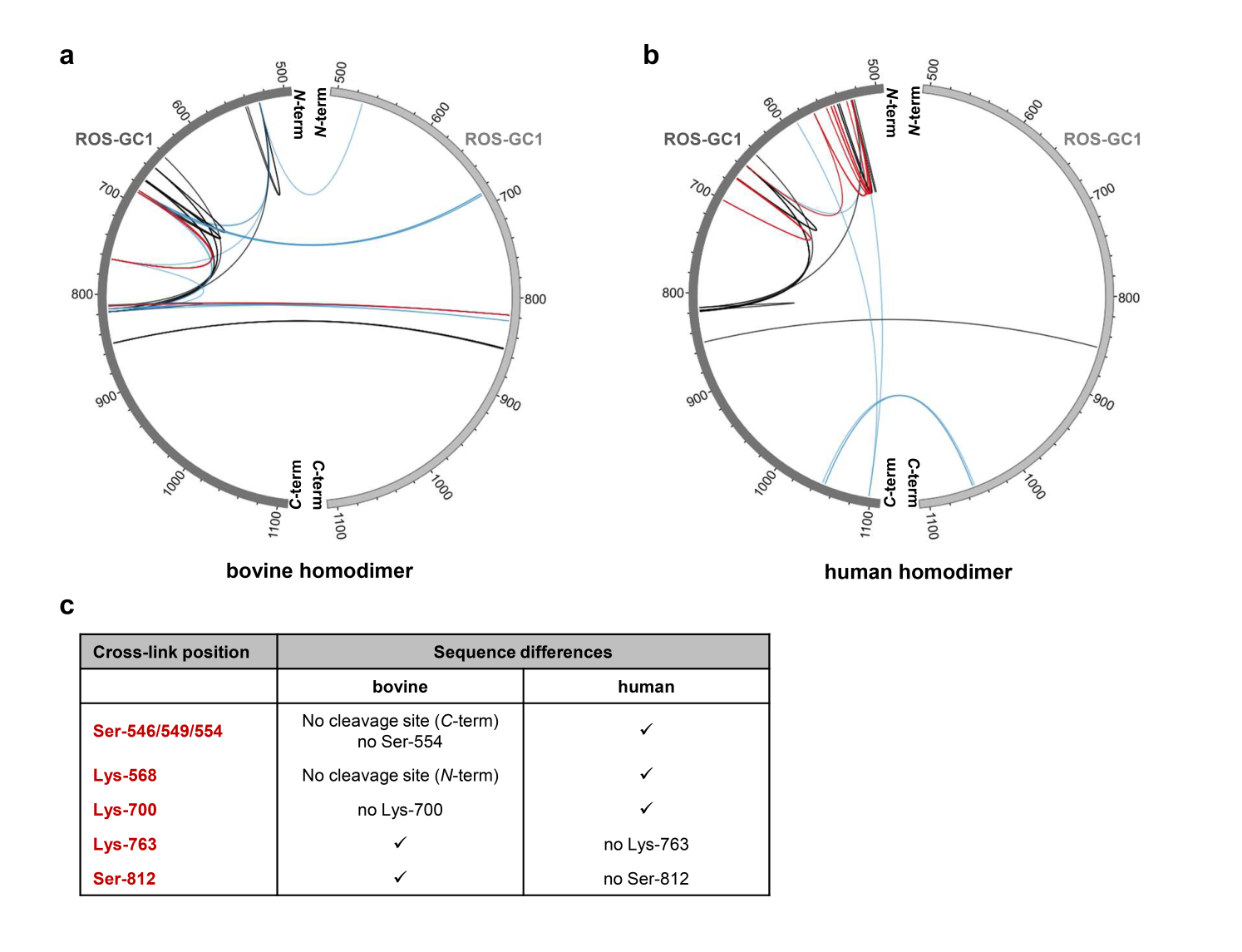

Supplementary Fig. 11. Comparison of DSBU Cross-links Identified in Bovine (a) and Human (b) ROS-GC1 Dimer. For comparison, numbering is given according to bovine ROS-GC1. Cross-links identified in bovine and human ROS-GC1 are shown in black, cross-links identified only in bovine or human ROS-GC1 are shown in blue, and crosslinks identified only in bovine or human ROS-GC1 dependent on different sequences are displayed in red. The Circos software was used for generating circular plots; c) cross-linking sites existing only in bovine or human ROS-GC1 as (i) Lys or Arg are missing as cleavage sites for trypsin for generating the specific cross-linked peptides or (ii) Lys or Ser are missing as cross-linking sites.

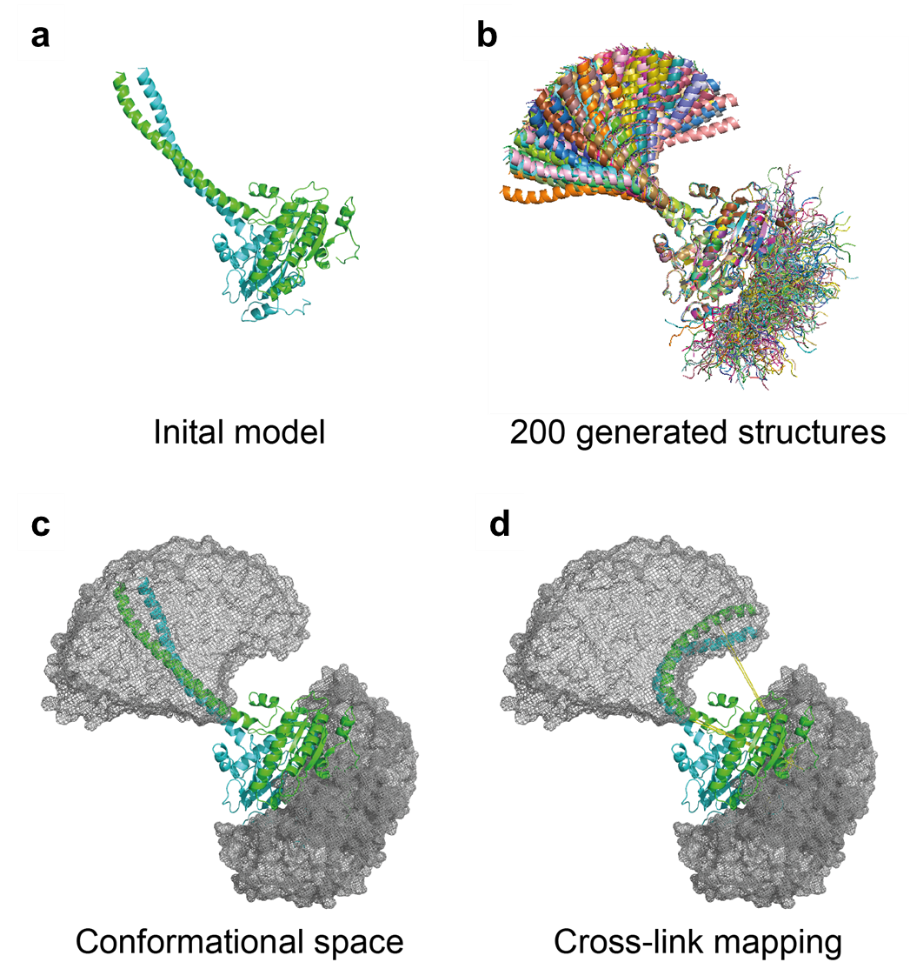

Supplementary Fig. 12. Exploring Torsion Angle Space of αHD and *C*-Terminus of CD. (a) Initial model, derived from the native IcD structure; b) Superposition of 200 generated models after adding 12 residues at the *C*-terminus by de-novo modelling; c) conformational space of αHD and CTE shown as mesh density; the initial model is superimposed; d) mapping of cross-links. Cross-links between αHD and CD are still violated, indicating even more refolding than captured by the remodeling approach.

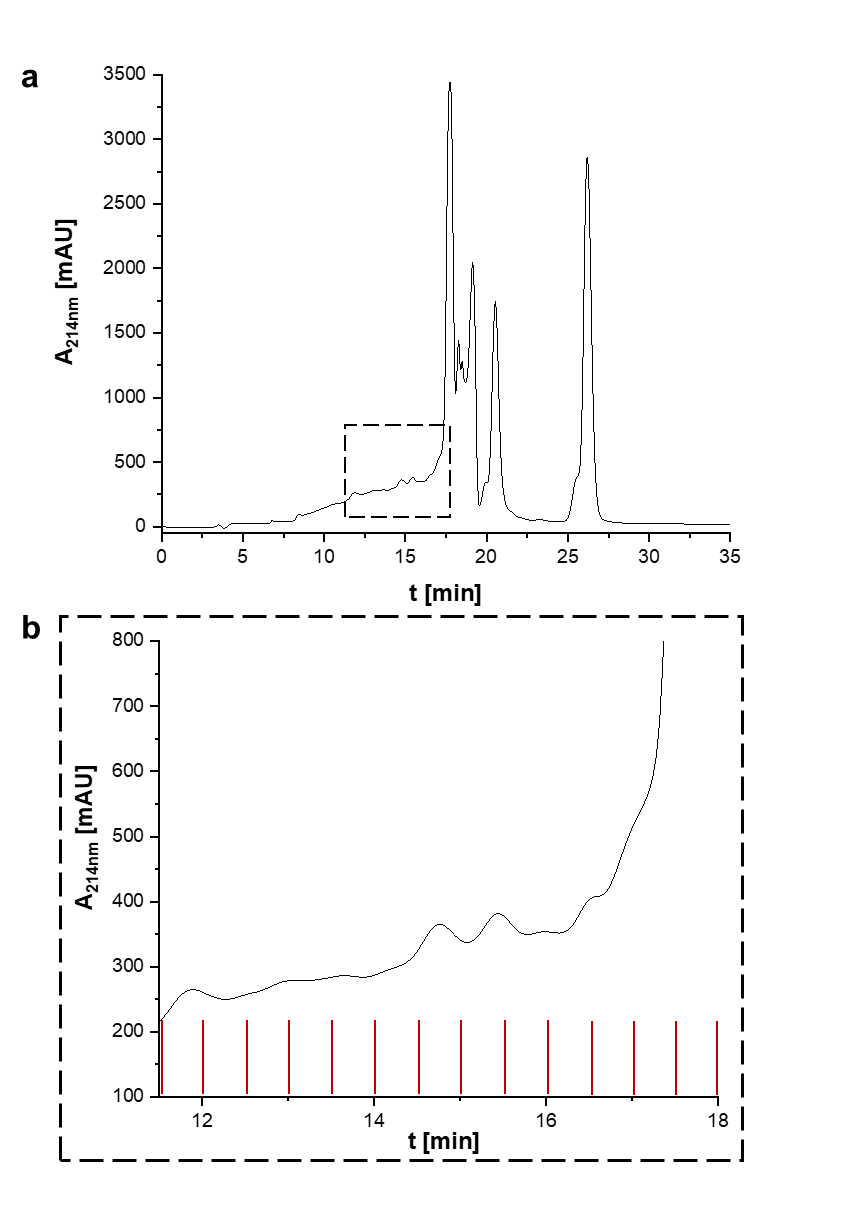

**Supplementary Fig. 13.** **Size Exclusion Chromatogram of Peptide Mixture from HEK293 Cell Lysate after Cross-linking with DSBU and *In-solution* Digestion**; **a**) SEC was performed with a Superdex 30 Increase 13/300 GL column; **b**) selected 500-µl fractions (dashed box in a) were analyzed by LC/MS/MS.

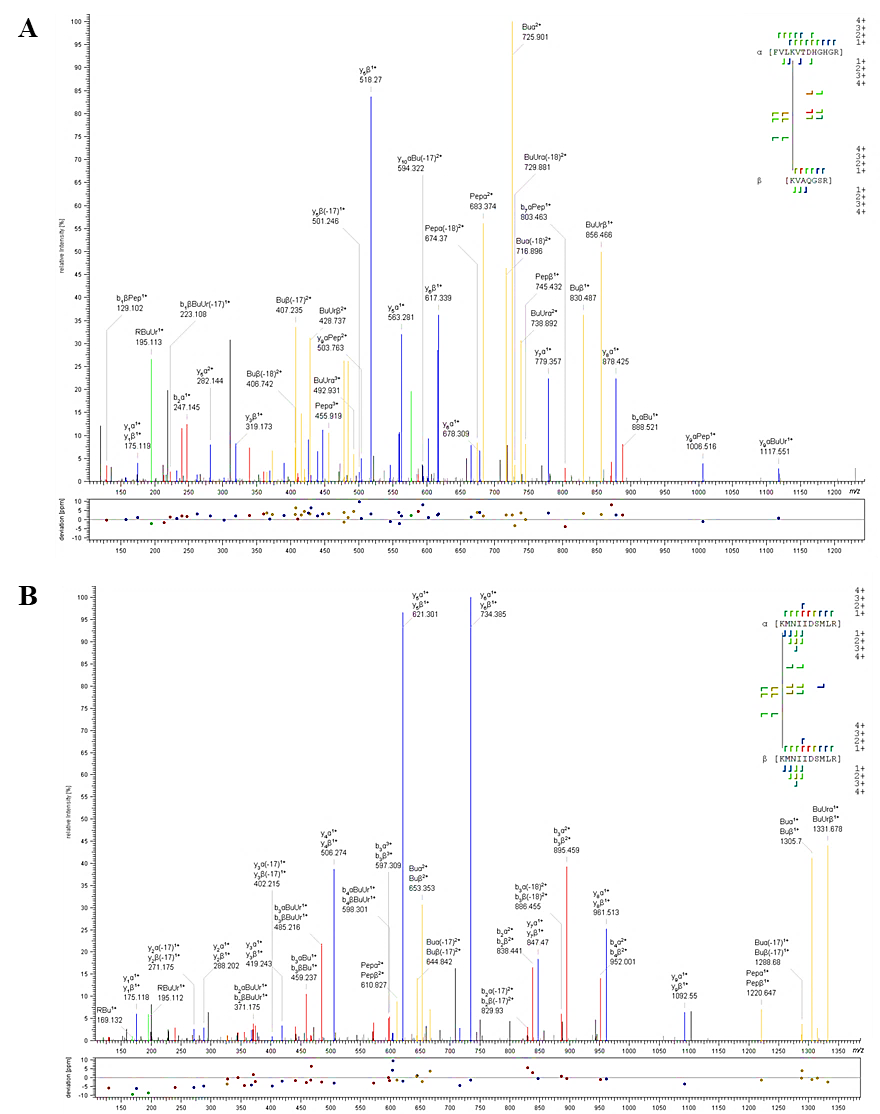

**a**

**b**

Supplementary Fig. 14. MS/MS Spectra of Cross-linked Products from Bovine, Full-length ROS-GC1. HCD-MS/MS spectra were automatically annotated with MeroX; a) A cross-link (4+ charged precursor ion at *m*/*z* 577.317) was identified between K686 (aa 683-694) and K527 (aa 527-533); b) A cross-link (4+ charged precursor ion at *m*/*z* 659.850) between K818 and K818 of identical peptides (aa 818-827) indicates an interprotein connection between two ROS-GC1 monomers. Precursor ions are shown in green, fragment ions of DSBU are shown in yellow, b- and y-type ions of the peptide backbone are shown in red and blue.

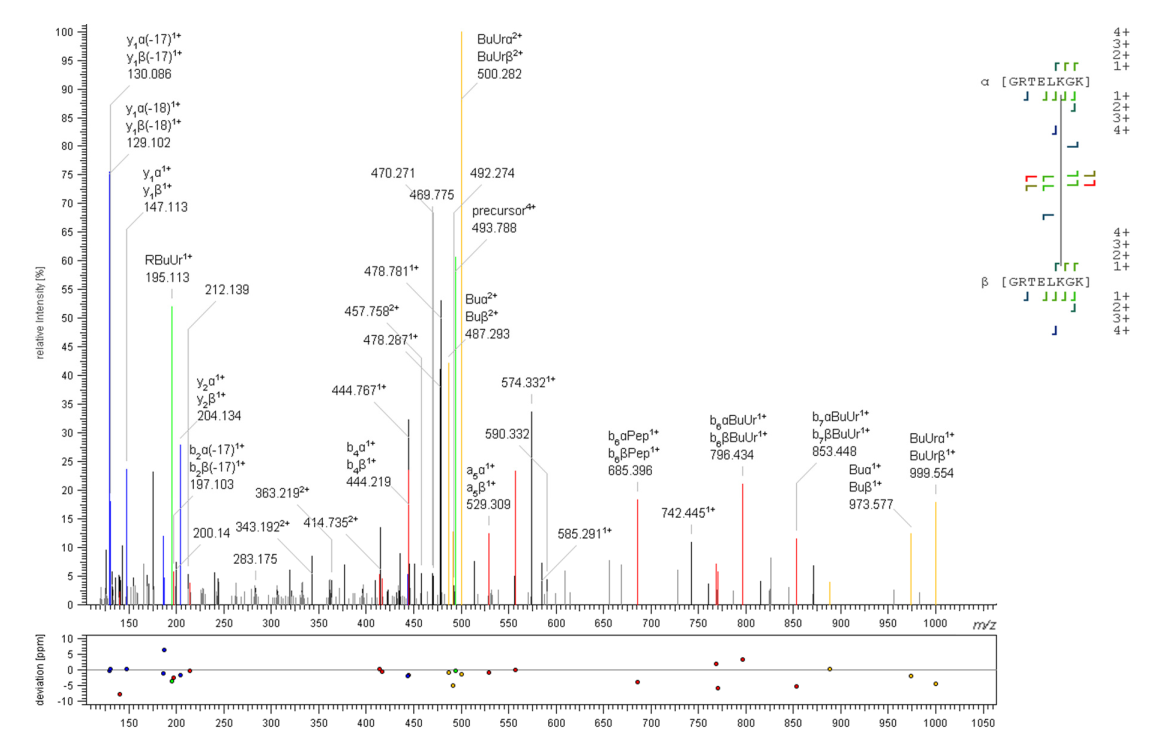

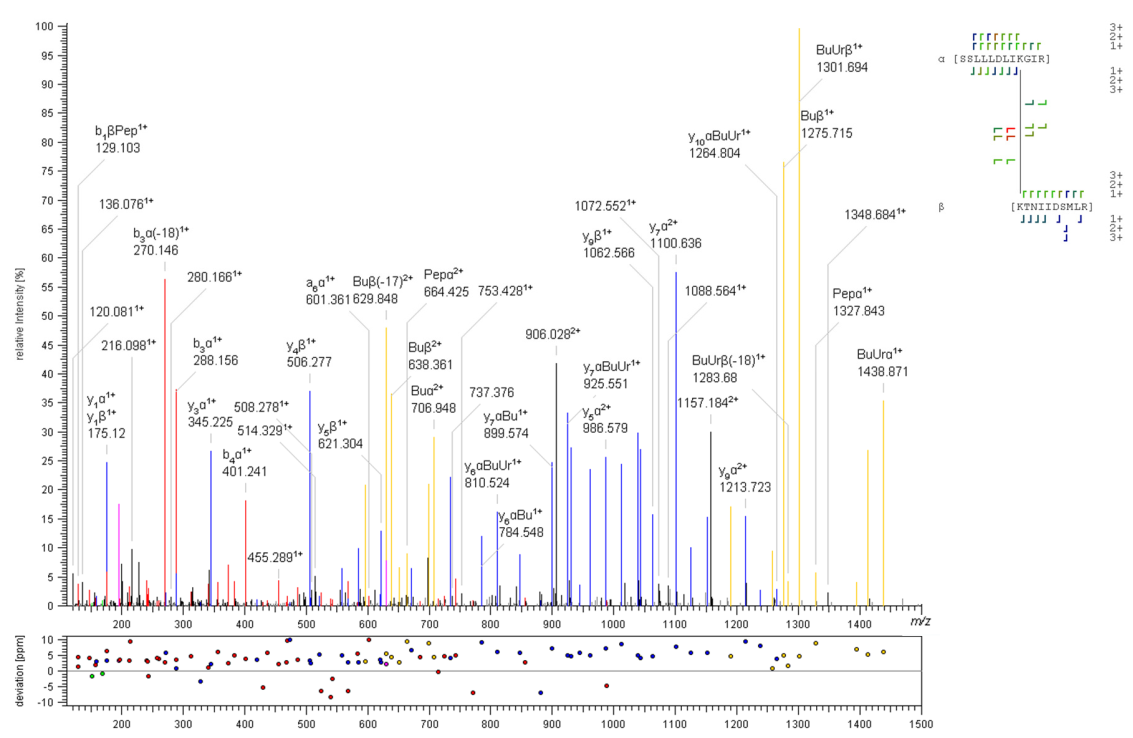

**a**

**b**

**Supplementary Fig. 15. MS/MS Spectra of Cross-linked Products from Human, Full-length ROS-GC1.** HCD-MS/MS spectra were automatically annotated with MeroX; **a)** A cross-link (3+ charged precursor ion at *m*/*z* 905.196) was identified between K652 (aa 644-655) and K813 of (aa 813-822); **b)** A cross-link (4+ charged precursor ion at *m*/*z* 493.787) between K1046 and K1046 of identical peptides (aa 1041-1048) indicates an interprotein cross-link between two ROS-GC1 monomers. Precursor ions are shown in green, fragment ions of DSBU are shown in yellow, b- and y-type ions of the peptide backbone are shown in red and blue.

Supplementary Table 1. DSBU Cross-links in Bovine, Full-length ROS-GC1 in ROS-Preparation. If a cross-linking site is ambiguous, all potential cross-linked amino acids are listed. The cross-linking reactions were conducted in the absence of calcium.

| **DSBU** | | | | | | | |
| --- | --- | --- | --- | --- | --- | --- | --- |
|  | **[M+H]^+^** |  | **deviation** | **cross-linked** |  | **cross-linked** |  |
| ***m/z*** | **theor.** | **z** | **(ppm)** | **peptide (1)** | **site (1)** | **peptide (2)** | **site (2)** |
| 659.851 | 2636.375 | 4 | 2.19 | [KMNIIDSMLR] | K818 | [KMNIIDSMLR] | K818 |
| 586.117 | 2926.555 | 5 | 0.94 | [FVLKVTDHGHGR] | K686 | [FVLKVTDHGHGR] | K686 |
| 659.852 | 2636.375 | 4 | 4.32 | [KMNIIDSMLR] | K818 | [KMNIIDSMLR] | K818 |
| 586.117 | 2926.555 | 5 | 0.94 | [FVLKVTDHGHGR] | K686 | [FVLKVTDHGHGR] | K686 |
| 586.117 | 2926.555 | 5 | 0.94 | [FVLKVTDHGHGR] | K686 | [FVLKVTDHGHGR] | K686 |
| 488.599 | 2926.555 | 6 | 0.87 | [FVLKVTDHGHGR] | K686/T688 | [FVLKVTDHGHGR] | K686/T688 |
| 659.85 | 2636.375 | 4 | 1.73 | [KMNIIDSMLR] | K818 | [KMNIIDSMLR] | K818 |
| 488.599 | 2926.555 | 6 | 0.87 | [FVLKVTDHGHGR] | K686/T688 | [FVLKVTDHGHGR] | K686/T688 |
| 661.358 | 2642.404 | 4 | 2.59 | [TFELFKSINK] | K811/S812 | [KMNIIDSMLR] | K818 |
| 661.355 | 2642.404 | 4 | -2.22 | [TFELFKSINK] | K811/S812 | [KMNIIDSMLR] | K818 |
| 661.357 | 2642.404 | 4 | 1.29 | [TFELFKSINK] | K811 | [KMNIIDSMLR] | K818 |
| 661.358 | 2642.404 | 4 | 2.59 | [TFELFKSINK] | K811/S812 | [KMNIIDSMLR] | K818 |
| 661.358 | 2642.404 | 4 | 2.59 | [TFELFKSINK] | K811/S812 | [KMNIIDSMLR] | K818 |
| 661.357 | 2642.404 | 4 | 1.02 | [TFELFKSINK] | K811 | [KMNIIDSMLR] | K818 |
| 586.117 | 2926.555 | 5 | 1.04 | [FVLKVTDHGHGR] | K686/T688 | [FVLKVTDHGHGR] | K686/T688 |
| 661.358 | 2642.404 | 4 | 2.59 | [TFELFKSINK] | K811/S812 | [KMNIIDSMLR] | K818 |
| 661.358 | 2642.404 | 4 | 2.59 | [TFELFKSINK] | K811/S812 | [KMNIIDSMLR] | K818 |
| 661.358 | 2642.404 | 4 | 2.59 | [TFELFKSINK] | K811/S812 | [KMNIIDSMLR] | K818 |
| 881.475 | 2642.404 | 3 | 2.38 | [TFELFKSINK] | K811 | [KMNIIDSMLR] | K818 |
| 881.473 | 2642.404 | 3 | -0.11 | [TFELFKSINK] | K811 | [KMNIIDSMLR] | K818 |
| 661.358 | 2642.404 | 4 | 2.59 | [TFELFKSINK] | K811/S812 | [KMNIIDSMLR] | K818 |
| 659.85 | 2636.375 | 4 | 0.62 | [KMNIIDSMLR] | K818 | [KMNIIDSMLR] | K818 |
| 837.94 | 3348.736 | 4 | 0.6 | [SAPYAMLELTPEEVVKR] | K763 | [KMNIIDSMLR] | K818 |
| 837.94 | 3348.736 | 4 | 0.6 | [SAPYAMLELTPEEVVKR] | K763 | [KMNIIDSMLR] | K818 |
| 586.118 | 2926.555 | 5 | 2.19 | [FVLKVTDHGHGR] | K686/T688 | [FVLKVTDHGHGR] | K686/T688 |
| 413.635 | 2064.146 | 5 | 0.29 | [FVLKVTDHGHGR] | K686/T688 | [LKSR] | K673 |
| 577.318 | 2306.248 | 4 | 0.78 | [FVLKVTDHGHGR] | K686/T688 | [KVAQGSR] | K527 |
| 577.318 | 2306.248 | 4 | 0.78 | [FVLKVTDHGHGR] | K686/T688 | [KVAQGSR] | K27 |
| 837.944 | 3348.736 | 4 | 4.9 | [SAPYAMLELTPEEVVKR] | K763 | [KMNIIDSMLR] | K818 |
| 661.359 | 2642.404 | 4 | 4.43 | [TFELFKSINK] | K811/S812 | [KMNIIDSMLR] | K818 |
| 767.916 | 3068.65 | 4 | -2.4 | [TFELFKSINKGR] | K815 | [GRKMNIIDSMLR] | K818 |
| 767.916 | 3068.65 | 4 | -2.95 | [TFELFKSINKGR] | K815 | [GRKMNIIDSMLR] | S824 |
| 767.916 | 3068.65 | 4 | -2.4 | [TFELFKSINKGR] | K815 | [GRKMNIIDSMLR] | K818 |
| 767.916 | 3068.65 | 4 | -2.16 | [TFELFKSINKGR] | K815 | [GRKMNIIDSMLR] | K818/S824 |
| 767.918 | 3068.65 | 4 | 0.39 | [TFELFKSINKGR] | K815 | [GRKMNIIDSMLR] | K818 |
| 767.916 | 3068.65 | 4 | -2.16 | [TFELFKSINKGR] | K815 | [GRKMNIIDSMLR] | K818 |
| 837.938 | 3348.736 | 4 | -1.37 | [SAPYAMLELTPEEVVKR] | K763 | [KMNIIDSMLR] | K818 |
| 767.918 | 3068.65 | 4 | 0.39 | [TFELFKSINKGR] | K815 | [GRKMNIIDSMLR] | K818 |
| 767.918 | 3068.65 | 4 | 0.39 | [TFELFKSINKGR] | K815 | [GRKMNIIDSMLR] | K818 |
| 767.918 | 3068.65 | 4 | 0.39 | [TFELFKSINKGR] | K815 | [GRKMNIIDSMLR] | K818 |
| 767.917 | 3068.65 | 4 | -0.73 | [TFELFKSINKGR] | K815 | [GRKMNIIDSMLR] | K818 |
| 767.916 | 3068.65 | 4 | -2.16 | [TFELFKSINKGR] | K815 | [GRKMNIIDSMLR] | K818/S824 |
| 767.918 | 3068.65 | 4 | 0.39 | [TFELFKSINKGR] | K815 | [GRKMNIIDSMLR] | K818 |
| 837.938 | 3348.736 | 4 | -1.37 | [SAPYAMLELTPEEVVKR] | K763 | [KMNIIDSMLR] | K818 |
| 663.85 | 2652.37 | 4 | 2.81 | [KMNIIDSmLR] | K818 | [KMNIIDSMLR] | K818 |
| 732.395 | 2926.555 | 4 | 1.71 | [FVLKVTDHGHGR] | K686 | [FVLKVTDHGHGR] | K686 |
| 767.918 | 3068.65 | 4 | 0.39 | [TFELFKSINKGR] | K815 | [GRKMNIIDSMLR] | K818 |
| 837.938 | 3348.736 | 4 | -1.37 | [SAPYAMLELTPEEVVKR] | K763 | [KMNIIDSMLR] | K818 |
| 699.572 | 3493.826 | 5 | 1.07 | [SAPYAMLELTPEEVVKR] | K763 | [FVLKVTDHGHGR] | K686 |
| 767.918 | 3068.65 | 4 | 0.39 | [TFELFKSINKGR] | K815 | [GRKMNIIDSMLR] | K818 |
| 697.378 | 2090.121 | 3 | -1.23 | [KMNIIDSMLR] | K818 | [SINKGR] | K815 |
| 699.572 | 3493.826 | 5 | 1.07 | [SAPYAMLELTPEEVVKR] | K763 | [FVLKVTDHGHGR] | T688 |
| 697.38 | 2090.121 | 3 | 1.49 | [KMNIIDSMLR] | K818 | [SINKGR] | K815 |
| 699.571 | 3493.826 | 5 | -0.67 | [SAPYAMLELTPEEVVKR] | K763 | [FVLKVTDHGHGR] | K686 |
| 767.915 | 3068.65 | 4 | -3.67 | [TFELFKSINKGR] | K815 | [GRKMNIIDSMLR] | K818 |
| 697.375 | 2090.121 | 3 | -4.73 | [KMNIIDSMLR] | K818 | [SINKGR] | K815 |
| 661.359 | 2642.404 | 4 | 3.79 | [TFELFKSINK] | K811/S812 | [KMNIIDSMLR] | K818 |
| 697.379 | 2090.121 | 3 | 0.7 | [KMNIIDSMLR] | K818 | [SINKGR] | K815 |
| 767.917 | 3068.65 | 4 | -0.81 | [TFELFKSINKGR] | K815 | [GRKMNIIDSMLR] | K818 |
| 640.359 | 1919.056 | 3 | 3.4 | [KMNIIDSMLR] | K818 | [LKSR] | K673/S674 |
| 699.574 | 3493.826 | 5 | 3.96 | [SAPYAMLELTPEEVVKR] | K763 | [FVLKVTDHGHGR] | K686 |
| 640.359 | 1919.056 | 3 | 3.4 | [KMNIIDSMLR] | K818 | [LKSR] | K673/S674 |
| 733.107 | 2197.303 | 3 | 2.38 | [SSLLLDLIKGIR] | K657 | [SINKGR] | K815 |
| 697.379 | 2090.121 | 3 | 0.7 | [KMNIIDSMLR] | K818 | [SINKGR] | K815 |
| 697.375 | 2090.121 | 3 | -4.73 | [KMNIIDSMLR] | K818 | [SINKGR] | K815 |
| 676.085 | 2026.238 | 3 | 1.55 | [SSLLLDLIKGIR] | K657 | [LKSR] | K673/S674 |
| 767.914 | 3068.65 | 4 | -4.7 | [TFELFKSINKGR] | K815 | [GRKMNIIDSMLR] | K818 |
| 733.109 | 2197.303 | 3 | 4.29 | [SSLLLDLIKGIR] | K657 | [SINKGR] | S812/K815 |
| 653.591 | 2611.344 | 4 | -0.88 | [KMNIIDSMLR] | K818 | [DIKLDWMFK] | K642 |
| 516.792 | 2064.146 | 4 | 0.6 | [FVLKVTDHGHGR] | K686 | [LKSR] | K673 |
| 767.918 | 3068.65 | 4 | 0.39 | [TFELFKSINKGR] | K815 | [GRKMNIIDSMLR] | K818 |
| 640.36 | 1919.056 | 3 | 4.45 | [KMNIIDSMLR] | K818 | [LKSR] | K673/S674 |
| 516.792 | 2064.146 | 4 | 0.84 | [FVLKVTDHGHGR] | K686 | [LKSR] | K673 |
| 767.916 | 3068.65 | 4 | -2.16 | [TFELFKSINKGR] | K815 | [GRKMNIIDSMLR] | K818 |
| 767.918 | 3068.65 | 4 | 0.39 | [TFELFKSINKGR] | K815 | [GRKMNIIDSMLR] | K818 |
| 640.36 | 1919.056 | 3 | 4.35 | [KMNIIDSMLR] | K818 | [LKSR] | K673/S674 |
| 697.379 | 2090.121 | 3 | 0.26 | [KMNIIDSMLR] | K818 | [SINKGR] | K815 |
| 676.085 | 2026.238 | 3 | 0.65 | [SSLLLDLIKGIR] | K657 | [LKSR] | K673/S674 |
| 733.107 | 2197.303 | 3 | 2.38 | [SSLLLDLIKGIR] | K657 | [SINKGR] | K815 |
| 721.058 | 2161.158 | 3 | 1.32 | [KMNIIDSMLR] | K818 | [KVAQGSR] | K527 |
| 516.792 | 2064.146 | 4 | 0.48 | [FVLKVTDHGHGR] | K686/T688 | [LKSR] | K673 |
| 516.792 | 2064.146 | 4 | 0.48 | [FVLKVTDHGHGR] | T688 | [LKSR] | K673 |
| 661.356 | 2642.404 | 4 | -0.28 | [TFELFKSINK] | K811/S812 | [KMNIIDSMLR] | K818 |
| 697.379 | 2090.121 | 3 | 0.26 | [KMNIIDSMLR] | K818 | [SINKGR] | K815 |
| 640.359 | 1919.056 | 3 | 2.35 | [KMNIIDSMLR] | K818 | [LKSR] | K673/S674 |
| 767.918 | 3068.65 | 4 | 0.39 | [TFELFKSINKGR] | K815 | [GRKMNIIDSMLR] | K818 |
| 721.059 | 2161.158 | 3 | 2 | [KMNIIDSMLR] | K818 | [KVAQGSR] | K527 |
| 721.059 | 2161.158 | 3 | 2 | [KMNIIDSMLR] | K818 | [KVAQGSR] | K527 |
| 699.574 | 3493.826 | 5 | 3.96 | [SAPYAMLELTPEEVVKR] | K763 | [FVLKVTDHGHGR] | K686/T688 |
| 767.917 | 3068.65 | 4 | -1.28 | [TFELFKSINKGR] | K815 | [GRKMNIIDSMLR] | K818/S824 |
| 661.356 | 2642.404 | 4 | -0.28 | [TFELFKSINK] | K811/S812 | [KMNIIDSMLR] | K818 |
| 767.918 | 3068.65 | 4 | 0.39 | [TFELFKSINKGR] | K815 | [GRKMNIIDSMLR] | K818 |
| 767.918 | 3068.65 | 4 | 0.39 | [TFELFKSINKGR] | K815 | [GRKMNIIDSMLR] | S824 |
| 702.712 | 2106.116 | 3 | 2.82 | [KmNIIDSMLR] | K818 | [SINKGR] | K815 |
| 676.083 | 2026.238 | 3 | -1.34 | [SSLLLDLIKGIR] | K657 | [LKSR] | K673/S674 |
| 676.083 | 2026.238 | 3 | -1.34 | [SSLLLDLIKGIR] | K657 | [LKSR] | K673/S674 |
| 702.712 | 2106.116 | 3 | 2.82 | [KmNIIDSMLR] | K818 | [SINKGR] | K815 |
| 640.358 | 1919.056 | 3 | 1.3 | [KMNIIDSMLR] | K818 | [LKSR] | K673/S674 |
| 767.914 | 3068.65 | 4 | -4.7 | [TFELFKSINKGR] | K815 | [GRKMNIIDSMLR] | K818 |
| 516.792 | 2064.146 | 4 | 0.6 | [FVLKVTDHGHGR] | K686 | [LKSR] | K673 |
| 516.792 | 2064.146 | 4 | 0.6 | [FVLKVTDHGHGR] | K686 | [LKSR] | K673 |
| 733.105 | 2197.303 | 3 | -0.96 | [SSLLLDLIKGIR] | K657 | [SINKGR] | K815 |
| 699.57 | 3493.826 | 5 | -1.29 | [SAPYAMLELTPEEVVKR] | K763 | [FVLKVTDHGHGR] | K686 |
| 697.379 | 2090.121 | 3 | 0.87 | [KMNIIDSMLR] | K818 | [SINKGR] | K815 |
| 697.379 | 2090.121 | 3 | 0.87 | [KMNIIDSMLR] | K818 | [SINKGR] | K815 |
| 697.376 | 2090.121 | 3 | -3.07 | [KMNIIDSMLR] | K818 | [SINKGR] | K815 |
| 696.125 | 2781.465 | 4 | 4.18 | [FVLKVTDHGHGR] | K686/T688 | [KMNIIDSMLR] | K818 |
| 719.139 | 2873.519 | 4 | 4.84 | [SAPYAMLELTPEEVVKR] | K763 | [KVAQGSR] | K527 |
| 719.138 | 2873.519 | 4 | 3.65 | [SAPYAMLELTPEEVVKR] | K763 | [KVAQGSR] | K527 |
| 516.792 | 2064.146 | 4 | 0.37 | [FVLKVTDHGHGR] | K686/T688 | [LKSR] | K673 |
| 719.139 | 2873.519 | 4 | 4.84 | [SAPYAMLELTPEEVVKR] | K763 | [KVAQGSR] | K527 |
| 653.594 | 2611.344 | 4 | 3.7 | [KMNIIDSMLR] | K818 | [DIKLDWMFK] | K642 |
| 659.85 | 2636.375 | 4 | 0.53 | [KMNIIDSMLR] | K818 | [KMNIIDSMLR] | K818 |
| 601.325 | 2402.272 | 4 | 2.54 | [KAHDAVLTLTR] | K308 | [HBPLGGSVR] | S325 |
| 697.378 | 2090.121 | 3 | -1.23 | [KMNIIDSMLR] | K818 | [SINKGR] | K815 |
| 697.376 | 2090.121 | 3 | -3.07 | [KMNIIDSMLR] | K818 | [SINKGR] | K815 |
| 516.792 | 2064.146 | 4 | 0.84 | [FVLKVTDHGHGR] | K686/T688 | [LKSR] | K673 |
| 653.594 | 2611.344 | 4 | 3.7 | [KMNIIDSMLR] | K818 | [DIKLDWMFK] | K642 |
| 697.38 | 2090.121 | 3 | 1.49 | [KMNIIDSMLR] | K818 | [SINKGR] | K815 |
| 601.325 | 2402.272 | 4 | 2.13 | [KAHDAVLTLTR] | K308 | [HBPLGGSVR] | S325 |
| 697.378 | 2090.121 | 3 | -1.23 | [KMNIIDSMLR] | K818 | [SINKGR] | K815 |
| 697.376 | 2090.121 | 3 | -3.07 | [KMNIIDSMLR] | K818 | [SINKGR] | K815 |
| 697.38 | 2090.121 | 3 | 2.1 | [KMNIIDSMLR] | K818 | [SINKGR] | K815 |
| 699.57 | 3493.826 | 5 | -1.29 | [SAPYAMLELTPEEVVKR] | K763 | [FVLKVTDHGHGR] | K686/T688 |
| 697.38 | 2090.121 | 3 | 2.1 | [KMNIIDSMLR] | K818 | [SINKGR] | K815 |
| 719.134 | 2873.519 | 4 | -1.36 | [SAPYAMLELTPEEVVKR] | K763 | [KVAQGSR] | K527 |
| 601.325 | 2402.272 | 4 | 2.03 | [KAHDAVLTLTR] | K308 | [HBPLGGSVR] | S325 |
| 696.122 | 2781.465 | 4 | -0.12 | [FVLKVTDHGHGR] | K686/T688 | [KMNIIDSMLR] | K818 |

Supplementary Table 2. DSBU Cross-links of Bovine, Full-length ROS-GC1 in ROS-Preparation after Glutaraldehyde Fixiation. Cross-linking experiments were conducted in the absence of calcium. If a cross-linking site is ambiguous, all potential cross-linked amino acids are listed.

| **DSBU** | | | | | | | |
| --- | --- | --- | --- | --- | --- | --- | --- |
|  | **[M+H]^+^** |  | **deviation** | **cross-linked** |  | **cross-linked** |  |
| ***m/z*** | **theor.** | **z** | **(ppm)** | **peptide (1)** | **site (1)** | **peptide (2)** | **site (2)** |
| 659.85 | 2636.375 | 4 | 1.64 | [KMNIIDSMLR] | K818 | [KMNIIDSMLR] | K818 |
| 413.635 | 2064.146 | 5 | 0.36 | [FVLKVTDHGHGR] | K686/T688 | [LKSR] | K673 |
| 879.464 | 2636.375 | 3 | 1.09 | [KMNIIDSMLR] | K818 | [KMNIIDSMLR] | K818 |
| 881.474 | 2642.404 | 3 | 1.34 | [TFELFKSINK] | K811/S812 | [KMNIIDSMLR] | K818 |
| 837.94 | 3348.736 | 4 | 1.04 | [SAPYAMLELTPEEVVKR] | K763 | [KMNIIDSMLR] | K818 |
| 697.03 | 2089.077 | 3 | -0.16 | [TEELELEKQK] | K851 | [QKTDR] | K853/T854 |
| 697.031 | 2089.077 | 3 | 0.89 | [TEELELEKQK] | K851 | [QKTDR] | K853/T854 |
| 697.379 | 2090.121 | 3 | 1.14 | [KMNIIDSMLR] | K818 | [SINKGR] | K815 |
| 767.916 | 3068.65 | 4 | -1.84 | [TFELFKSINKGR] | K815 | [GRKMNIIDSMLR] | K818/S824 |
| 562.651 | 1685.941 | 3 | -0.59 | [KVAQGSR] | K527 | [KVAQGSR] | K527 |
| 697.379 | 2090.121 | 3 | 1.14 | [KMNIIDSMLR] | K818 | [SINKGR] | K815 |
| 697.032 | 2089.077 | 3 | 1.59 | [TEELELEKQK] | K851 | [QKTDR] | K853/T854 |
| 539.629 | 1616.871 | 3 | 0.1 | [KVAQGSR] | K527 | [SISDVR] | K540/K542 |
| 837.94 | 3348.736 | 4 | 1.04 | [SAPYAMLELTPEEVVKR] | K763 | [KMNIIDSMLR] | K818 |
| 837.943 | 3348.736 | 4 | 3.88 | [SAPYAMLELTPEEVVKR] | K763 | [KMNIIDSMLR] | K818 |
| 697.379 | 2090.121 | 3 | 1.05 | [KMNIIDSMLR] | K818 | [SINKGR] | K815 |
| 697.379 | 2090.121 | 3 | 1.05 | [KMNIIDSMLR] | K818 | [SINKGR] | K815 |
| 879.464 | 2636.375 | 3 | 0.46 | [KMNIIDSMLR] | K818 | [KMNIIDSMLR] | K818 |
| 697.379 | 2090.121 | 3 | 0.09 | [KMNIIDSMLR] | K818 | [SINKGR] | K815 |
| 697.38 | 2090.121 | 3 | 1.57 | [KMNIIDSMLR] | K818 | [SINKGR] | K815 |
| 697.379 | 2090.121 | 3 | 0.35 | [KMNIIDSMLR] | K818 | [SINKGR] | K815 |
| 697.38 | 2090.121 | 3 | 1.57 | [KMNIIDSMLR] | K818 | [SINKGR] | S812/K815 |
| 539.629 | 1616.871 | 3 | 0.1 | [KVAQGSR] | K527 | [SISDVR] | K540/K542 |
| 697.031 | 2089.077 | 3 | 0.8 | [TEELELEKQK] | K851 | [QKTDR] | K853/T854 |
| 697.378 | 2090.121 | 3 | 0 | [KMNIIDSMLR] | K818 | [SINKGR] | K815 |
| 767.917 | 3068.65 | 4 | -1.2 | [TFELFKSINKGR] | K815 | [GRKMNIIDSMLR] | K818/S824 |
| 837.943 | 3348.736 | 4 | 3.88 | [SAPYAMLELTPEEVVKR] | K763 | [KMNIIDSMLR] | K818 |
| 562.652 | 1685.941 | 3 | 0.5 | [KVAQGSR] | K527 | [KVAQGSR] | K527 |
| 697.03 | 2089.077 | 3 | -0.16 | [TEELELEKQK] | K851 | [QKTDR] | K853 |
| 697.38 | 2090.121 | 3 | 1.84 | [KMNIIDSMLR] | K818 | [SINKGR] | K815 |
| 697.38 | 2090.121 | 3 | 1.84 | [KMNIIDSMLR] | K818 | [SINKGR] | K815 |
| 562.652 | 1685.941 | 3 | 0.93 | [KVAQGSR] | K527 | [KVAQGSR] | K527 |
| 767.917 | 3068.65 | 4 | -1.68 | [TFELFKSINKGR] | K815 | [GRKMNIIDSMLR] | K818/S824 |
| 792.079 | 2374.221 | 3 | 0.41 | [ERTEELELEKQK] | K851 | [QKTDR] | K853/T854 |
| 523.287 | 2090.121 | 4 | 1.92 | [KMNIIDSMLR] | K818 | [SINKGR] | K815 |
| 661.357 | 2642.404 | 4 | 0.46 | [TFELFKSINK] | K811/S812 | [KMNIIDSMLR] | K818 |
| 562.652 | 1685.941 | 3 | 0.06 | [KVAQGSR] | K527 | [KVAQGSR] | K527 |
| 562.652 | 1685.941 | 3 | 0.06 | [KVAQGSR] | K527 | [KVAQGSR] | K527 |
| 562.652 | 1685.941 | 3 | 0.93 | [KVAQGSR] | K527 | [KVAQGSR] | K527 |
| 879.464 | 2636.375 | 3 | 0.46 | [KMNIIDSMLR] | K818 | [KMNIIDSMLR] | K818 |
| 697.378 | 2090.121 | 3 | 0 | [KMNIIDSMLR] | K818 | [SINKGR] | S812/K815 |
| 562.652 | 1685.941 | 3 | 0.5 | [KVAQGSR] | K527 | [KVAQGSR] | K527 |
| 697.032 | 2089.077 | 3 | 1.59 | [TEELELEKQK] | K851 | [QKTDR] | K853/T854 |
| 697.031 | 2089.077 | 3 | 0.8 | [TEELELEKQK] | K851 | [QKTDR] | K853/T854 |
| 697.031 | 2089.077 | 3 | 0.89 | [TEELELEKQK] | K851 | [QKTDR] | K853/T854 |
| 523.284 | 2090.121 | 4 | -2.75 | [KMNIIDSMLR] | K818 | [SINKGR] | K815 |
| 879.464 | 2636.375 | 3 | 0.32 | [KMNIIDSMLR] | K818 | [KMNIIDSMLR] | K818 |
| 879.464 | 2636.375 | 3 | 0.32 | [KMNIIDSMLR] | K818 | [KMNIIDSMLR] | K818 |

Supplementary Table 3. DSBU Cross-links in Human, Full-length ROS-GC1. If a cross-linking site is ambiguous, all potential cross-linked amino acids are listed. The cross-linking reactions were conducted in the presence of GCAP-2 and in the absence of calcium. The human nomenclature is applied for ROS-GC1.

| **DSBU** | | | | | | | |
| --- | --- | --- | --- | --- | --- | --- | --- |
|  | **[M+H]^+^** |  | **deviation** | **cross-linked** |  | **cross-linked** |  |
| ***m/z*** | **theor.** | **z** | **(ppm)** | **peptide (1)** | **site (1)** | **peptide (2)** | **site (2)** |
| 440.508 | 1759.007 | 4 | 0.76 | [GRTELKGK] | K1046 | [TELKGK] | K1046 |
| 587.007 | 1759.007 | 4 | 0.22 | [GRTELKGK] | K1046 | [TELKGK] | K1046 |
| 607.332 | 3032.628 | 4 | 0.94 | [IILTVDDITFLHPHGGTSR] | T519/S520 | [KVAQGSR] | K522 |
| 607.334 | 3032.628 | 5 | 4.16 | [IILTVDDITFLHPHGGTSR] | T519/S520 | [KVAQGSR] | K522 |
| 1011.548 | 3032.628 | 4 | 0.83 | [IILTVDDITFLHPHGGTSR] | T519/S520 | [KVAQGSR] | K522 |
| 522.54 | 2087.139 | 4 | 0.04 | [KTNIIDSMLR] | K813 | [NINKGR] | K810 |
| 696.385 | 2087.139 | 3 | 0.1 | [KTNIIDSMLR] | K813 | [NINKGR] | K810 |
| 538.655 | 1613.945 | 4 | 3.93 | [KVAQGSR] | K522 | [RKLEK] | K1093 |
| 538.655 | 1613.945 | 3 | 3.93 | [KVAQGSR] | K522 | [RKLEK] | K1093 |
| 550.286 | 1648.844 | 5 | 0.3 | [KVAQGSR] | K522 | [SMSDIR] | S537 |
| 550.286 | 1648.844 | 3 | -0.58 | [KVAQGSR] | K522 | [SMSDIR] | S535/537 |
| 404.243 | 1613.949 | 4 | 0.86 | [KVAQGSR] | S527 | [VWLKK] | K563 |
| 538.655 | 1613.949 | 4 | 0.87 | [KVAQGSR] | S527 | [VWLKK] | K563 |
| 538.654 | 1613.949 | 4 | -0.61 | [KVAQGSR] | K522 | [VWLKK] | K563 |
| 404.242 | 1613.949 | 4 | -0.73 | [KVAQGSR] | K522 | [VWLKK] | K563 |
| 404.242 | 1613.949 | 4 | -0.73 | [KVAQGSR] | K522 | [VWLKK] | K563 |
| 742.869 | 2968.451 | 3 | 1.01 | [SGPSQHLDSPNIGVYEGDR] | S541/544 | [KVAQGSR] | K522 |
| 507.315 | 2026.238 | 3 | 0.73 | [SSLLLDLIKGIR] | K652 | [LKSR] | K668 |
| 556.833 | 2224.314 | 3 | -0.75 | [SSLLLDLIKGIR] | K652 | [NINKGR] | K810 |
| 556.834 | 2224.314 | 3 | 0.35 | [SSLLLDLIKGIR] | K562 | [NINKGR] | K810 |
| 549.844 | 2196.348 | 3 | 2.99 | [SSLLLDLIKGIR] | K652 | [VWLKK] | K563 |
| 387.226 | 1545.885 | 3 | -0.86 | [TELKGK] | K1046 | [TELKGK] | K1046 |

**Supplementary Table 4.** **DSBU Cross-links in Human, Full-length ROS-GC1.** If a cross-linking site is ambiguous, all potential cross-linked amino acids are listed. The cross-linking reactions were conducted in the absence of GCAP-2 and calcium. The human nomenclature is applied for ROS-GC1.

| **DSBU** | | | | | | | |
| --- | --- | --- | --- | --- | --- | --- | --- |
|  | **[M+H]^+^** |  | **deviation** | **cross-linked** |  | **cross-linked** |  |
| ***m/z*** | **theor.** | **z** | **(ppm)** | **peptide (1)** | **site (1)** | **peptide (2)** | **site (2)** |
| 538.656 | 1613.949 | 3 | 2.57 | [KVAQGSR] | K522 | [VWLKK] | K563 |
| 507.316 | 2026.238 | 4 | 1.64 | [SSLLLDLIKGIR] | K652 | [LKSR] | K668 |
| 556.835 | 2224.314 | 4 | 1.89 | [SSLLLDLIKGIR] | K652 | [NINKGR] | K810 |
| 567.841 | 2268.34 | 4 | 0.19 | [SSLLLDLIKGIR] | K652 | [KVAQGSR] | K522 |
| 538.655 | 1613.949 | 3 | 0.3 | [KVAQGSR] | K522 | [VWLKK] | K653 |
| 742.87 | 2968.451 | 4 | 2.74 | [SGPSQHLDSPNIGVYEGDR] | S541 | [KVAQGSR] | K522 |
| 990.156 | 2968.451 | 3 | 0.98 | [SGPSQHLDSPNIGVYEGDR] | S544 | [KVAQGSR] | K522 |
| 404.243 | 1613.949 | 4 | 0.56 | [KVAQGSR] | S527 | [VWLKK] | K563 |
| 840.452 | 2519.336 | 3 | 1.56 | [GKGAEDTFWLVGR] | K1048 | [GRTELKGK] | K1043/K1046 |
| 493.787 | 1972.13 | 4 | -0.87 | [GRTELKGK] | K1046 | [GRTELKGK] | K1046 |
| 696.386 | 2087.139 | 3 | 1.51 | [KTNIIDSMLR] | K813 | [NINKGR] | K810 |
| 697.031 | 2089.077 | 3 | 0.54 | [TEELELEKQK] | K846 | [QKTDR] | K848 |
| 607.331 | 3032.628 | 5 | -0.87 | [IILTVDDITFLHPHGGTSR] | T519/S520 | [KVAQGSR] | K522 |
| 493.788 | 1972.13 | 4 | 0.55 | [GRTELKGK] | K1046 | [GRTELKGK] | K1046 |

Supplementary Table 5. DSBU-D_0_/D_12_ Cross-links in Human, Full-length ROS-GC1. If a cross-linking site is ambiguous, all potential cross-linked amino acids are listed. The cross-linking reactions were conducted in the presence of GCAP-2 and in the absence of calcium. The human nomenclature is applied for ROS-GC1. Cross-links with DSBU-D_0_ are labeled as light (L) version, those with DSBU-D_12_ as heavy (H) version.

| **DSBU** | | | | | | | | |
| --- | --- | --- | --- | --- | --- | --- | --- | --- |
|  | **[M+H]^+^** |  | **deviation** | **cross-linked** |  | **cross-linked** |  | **DSBU** |
| ***m/z*** | **theor.** | **z** | **(ppm)** | **peptide (1)** | **site (1)** | **peptide (2)** | **site (2)** | **version** |
| 506.277 | 3032.628 | 6 | 0.21 | [IILTVDDITFLHPHGGTSR] | T519/S520 | [KVAQGSR] | K522 | (L) |
| 508.29 | 3044.703 | 6 | 0.25 | [IILTVDDITFLHPHGGTSR] | T519/S520 | [KVAQGSR] | K522 | (H) |
| 607.332 | 3032.628 | 5 | 1.34 | [IILTVDDITFLHPHGGTSR] | T519/S520 | [KVAQGSR] | K522 | (L) |
| 607.332 | 3032.628 | 5 | 1.64 | [IILTVDDITFLHPHGGTSR] | T519/S520 | [KVAQGSR] | K522 | (L) |
| 607.333 | 3032.628 | 5 | 2.05 | [IILTVDDITFLHPHGGTSR] | T519/S520 | [KVAQGSR] | K522 | (L) |
| 607.334 | 3032.628 | 5 | 3.66 | [IILTVDDITFLHPHGGTSR] | T519/S520 | [KVAQGSR] | K522 | (L) |
| 596.913 | 2980.526 | 5 | 2.83 | [SGPSQHLDSPNIGVYEGDR] | S541/S544/S549 | [KVAQGSR] | K522 | (H) |
| 745.888 | 2980.526 | 4 | 0.97 | [SGPSQHLDSPNIGVYEGDR] | S541/S544/S549 | [KVAQGSR] | K522 | (H) |
| 742.869 | 2968.451 | 4 | 1.42 | [SGPSQHLDSPNIGVYEGDR] | S541/S544 | [KVAQGSR] | K522 | (L) |
| 745.885 | 2980.526 | 4 | -2.15 | [SGPSQHLDSPNIGVYEGDR] | S541/S544 | [KVAQGSR] | K522 | (H) |
| 745.887 | 2980.526 | 4 | -0.1 | [SGPSQHLDSPNIGVYEGDR] | S541 | [KVAQGSR] | K522 | (H) |
| 519.07 | 2073.257 | 4 | 1.49 | [TAFSKLQELR] | K8583 | [RKLEK] | K1093 | (H) |
| 519.07 | 2073.257 | 4 | 0.9 | [TAFSKLQELR] | K8583 | [RKLEK] | K1093 | (H) |
| 519.072 | 2073.257 | 4 | 3.73 | [TAFSKLQELR] | K8583 | [RKLEK] | K1093 | (H) |
| 696.386 | 2087.139 | 3 | 1.51 | [KTNIIDSMLR] | K813 | [NINKGR] | K810 | (L) |
| 700.412 | 2099.214 | 3 | 3.04 | [KTNIIDSMLR] | K813 | [NINKGR] | K810 | (H) |
| 700.412 | 2099.214 | 3 | 2.77 | [KTNIIDSMLR] | K813 | [NINKGR] | K810 | (H) |
| 700.411 | 2099.214 | 3 | 1.82 | [KTNIIDSMLR] | K813 | [NINKGR] | K810 | (H) |
| 522.541 | 2087.139 | 4 | 0.86 | [KTNIIDSMLR] | K813 | [NINKGR] | K810 | (L) |
| 696.385 | 2087.139 | 3 | 1.24 | [KTNIIDSMLR] | K813 | [NINKGR] | K810 | (L) |
| 700.411 | 2099.214 | 3 | 1.47 | [KTNIIDSMLR] | K813 | [NINKGR] | K810 | (H) |
| 522.541 | 2087.139 | 4 | 1.21 | [KTNIIDSMLR] | K813 | [NINKGR] | K810 | (L) |
| 696.382 | 2087.139 | 3 | -3.14 | [KTNIIDSMLR] | K813 | [NINKGR] | K810 | (L) |
| 533.547 | 2131.165 | 4 | 1.25 | [KTNIIDSMLR] | K813 | [KVAQGSR] | K522 | (L) |
| 730.099 | 2188.281 | 3 | 0.81 | [LLEAQKVLPEPPR] | K695 | [LKSR] | K668/S669 | (L) |
| 547.826 | 2188.281 | 4 | 1.2 | [LLEAQKVLPEPPR] | K695 | [LKSR] | K668/S669 | (L) |
| 550.845 | 2200.357 | 4 | 1.36 | [LLEAQKVLPEPPR] | K695 | [LKSR] | K668/S669 | (H) |
| 547.826 | 2188.281 | 4 | 1.2 | [LLEAQKVLPEPPR] | K695 | [LKSR] | K668/S669 | (L) |
| 730.1 | 2188.281 | 3 | 1.9 | [LLEAQKVLPEPPR] | K695 | [LKSR] | K668/S669 | (L) |
| 909.219 | 2725.64 | 3 | 1.18 | [SSLLLDLIKGIR] | K652 | [KTNIIDSMLR] | K813/T814 | (H) |
| 686.164 | 2741.635 | 4 | -0.54 | [SSLLLDLIKGIR] | K652 | [KTNIIDSmLR] | K813/T814 | (H) |
| 905.197 | 2713.564 | 3 | 3.97 | [SSLLLDLIKGIR] | K652 | [KTNIIDSMLR] | K813 | (L) |
| 905.193 | 2713.564 | 3 | 0.26 | [SSLLLDLIKGIR] | K652 | [KTNIIDSMLR] | K813 | (L) |
| 682.166 | 2725.64 | 4 | 1.53 | [SSLLLDLIKGIR] | K652 | [KTNIIDSMLR] | K813 | (H) |
| 682.167 | 2725.64 | 4 | 2.61 | [SSLLLDLIKGIR] | K652 | [KTNIIDSMLR] | K813 | (H) |
| 679.148 | 2713.564 | 4 | 2.48 | [SSLLLDLIKGIR] | K652 | [KTNIIDSMLR] | K813 | (L) |
| 905.194 | 2713.564 | 3 | 1.61 | [SSLLLDLIKGIR] | K652 | [KTNIIDSMLR] | K813 | (L) |
| 905.195 | 2713.564 | 3 | 2.02 | [SSLLLDLIKGIR] | K652 | [KTNIIDSMLR] | K813 | (L) |
| 909.221 | 2725.64 | 3 | 3.26 | [SSLLLDLIKGIR] | K652 | [KTNIIDSMLR] | K813 | (H) |
| 679.147 | 2713.564 | 4 | 0.33 | [SSLLLDLIKGIR] | K652 | [KTNIIDSMLR] | K813 | (L) |
| 905.195 | 2713.564 | 3 | 2.49 | [SSLLLDLIKGIR] | K652 | [KTNIIDSMLR] | K813 | (L) |
| 679.148 | 2713.564 | 4 | 2.21 | [SSLLLDLIKGIR] | K652 | [KTNIIDSMLR] | K813 | (L) |
| 679.147 | 2713.564 | 4 | 0.96 | [SSLLLDLIKGIR] | K652 | [KTNIIDSMLR] | K813 | (L) |
| 682.168 | 2725.64 | 4 | 3.32 | [SSLLLDLIKGIR] | K652 | [KTNIIDSMLR] | K813 | (H) |
| 905.197 | 2713.564 | 3 | 4.04 | [SSLLLDLIKGIR] | K652 | [KTNIIDSMLR] | K813 | (L) |
| 556.834 | 2224.314 | 4 | 0.79 | [SSLLLDLIKGIR] | K652 | [NINKGR] | K810 | (L) |
| 556.834 | 2224.314 | 4 | 0.79 | [SSLLLDLIKGIR] | K652 | [NINKGR] | K810 | (L) |
| 556.835 | 2224.314 | 4 | 1.56 | [SSLLLDLIKGIR] | K652 | [NINKGR] | K810 | (L) |
| 559.853 | 2236.389 | 4 | 0.29 | [SSLLLDLIKGIR] | K652 | [NINKGR] | K810 | (H) |
| 746.135 | 2236.389 | 3 | 0.79 | [SSLLLDLIKGIR] | K652 | [NINKGR] | K810 | (H) |
| 746.135 | 2236.389 | 3 | 1.12 | [SSLLLDLIKGIR] | K652 | [NINKGR] | K810 | (H) |
| 746.137 | 2236.389 | 3 | 3.41 | [SSLLLDLIKGIR] | K652 | [NINKGR] | K810 | (H) |
| 746.137 | 2236.389 | 3 | 3.41 | [SSLLLDLIKGIR] | K652 | [NINKGR] | K810 | (H) |
| 742.112 | 2224.314 | 3 | 3.54 | [SSLLLDLIKGIR] | K652 | [NINKGR] | K810 | (L) |
| 742.11 | 2224.314 | 3 | 1.48 | [SSLLLDLIKGIR] | K652 | [NINKGR] | K810 | (L) |
| 746.135 | 2236.389 | 3 | 1.12 | [SSLLLDLIKGIR] | K652 | [NINKGR] | K810 | (H) |
| 746.133 | 2236.389 | 3 | -1.5 | [SSLLLDLIKGIR] | K652 | [NINKGR] | K810 | (H) |
| 556.835 | 2224.314 | 4 | 2.54 | [SSLLLDLIKGIR] | K652 | [NINKGR] | K810 | (L) |
| 556.836 | 2224.314 | 4 | 4.74 | [SSLLLDLIKGIR] | K652 | [NINKGR] | K810 | (L) |
| 742.11 | 2224.314 | 3 | 0.99 | [SSLLLDLIKGIR] | K652 | [NINKGR] | K810 | (L) |
| 510.334 | 2038.314 | 4 | -0.65 | [SSLLLDLIKGIR] | K652 | [LKSR] | K668/S669 | (H) |
| 702.711 | 2106.116 | 3 | 1.38 | [EIKLDWMFK] | K637 | [NINKGR] | K810 | (L) |
| 554.312 | 1660.919 | 3 | 1.03 | [KVAQGSR] | K522 | [SMSDIR] | S535/S537 | (H) |
| 550.286 | 1648.844 | 3 | -0.25 | [KVAQGSR] | K522 | [SMSDIR] | S535 | (L) |
| 404.243 | 1613.949 | 4 | -0.05 | [KVAQGSR] | K522 | [VWLKK] | K563 | (L) |
| 407.262 | 1626.024 | 4 | 0.7 | [KVAQGSR] | K522 | [VWLKK] | K563 | (H) |
| 542.68 | 1626.024 | 3 | 0.26 | [KVAQGSR] | K522 | [VWLKK] | K563 | (H) |
| 542.679 | 1626.024 | 3 | -0.87 | [KVAQGSR] | K522 | [VWLKK] | K563 | (H) |
| 538.654 | 1613.949 | 3 | 0.19 | [KVAQGSR] | K522 | [VWLKK] | K563 | (L) |
| 395.232 | 1972.13 | 5 | 0.85 | [GRTELKGK] | K1046 | [GRTELKGK] | K1046 | (L) |

Supplementary Table 6. Unique DSBU Cross-linking Sites in Human, Full-length ROS-GC1. The numbering of cross-linked amino acids is compared for bovine and human ROS-GC1. For the nomenclature of ROS-GC1 domains, please see Fig. 1b. If a cross-linking site is ambiguous, all potential cross-linked amino acids are listed.

| **Cross-linking site 1** | | **Cross-linking site2** | | **Domains** |
| --- | --- | --- | --- | --- |
| **bovine** | **human** | **bovine** | **human** |  |
| 524/525 | 519/520 | 527 | 522 | KHD - KHD |
| 527 | 522 | 540 | 535 | KHD - KHD |
| 527 | 522 | 540/542 | 535/537 | KHD - KHD |
| 527 | 522 | 542 | 537 | KHD - KHD |
| 527 | 522 | 546 | 541 | KHD - KHD |
| 527 | 522 | 546/549 | 541/544 | KHD - KHD |
| 527 | 522 | 549 | 544 | KHD - KHD |
| 527 | 522 | 546/549/554 | 541/544/549 | KHD - KHD |
| 527 | 522 | 568 | 563 | KHD - KHD |
| 527 | 522 | 657 | 652 | KHD - KHD |
| 532 | 527 | 568 | 563 | KHD - KHD |
| 568 | 563 | 657 | 652 | KHD - KHD |
| 657 | 652 | 673 | 668 | KHD - KHD |
| 657 | 652 | 673/674 | 668/669 | KHD - KHD |
| 673/674 | 668/669 | 700 | 695 | KHD - KHD |
| 527 | 522 | 818 | 813 | KHD - αHD |
| 642 | 637 | 815 | 810 | KHD - αHD |
| 657 | 652 | 815 | 810 | KHD - αHD |
| 657 | 652 | 818 | 813 | KHD - αHD |
| 657 | 652 | 819 | 814 | KHD - αHD |
| 657 | 652 | 818/819 | 813/814 | KHD - αHD |
| 815 | 810 | 818 | 813 | αHD -αHD |
| 527 | 522 | 1098 | 1093 | KHD - CD |
| 588 | 583 | 1098 | 1093 | KHD - CD |
| 851 | 846 | 853 | 848 | αHD -αHD |
| 1051 | 1046 | 1051 | 1046 | CD - CD |
| 1048/1051 | 1043/1046 | 1053 | 1048 | CD - CD |

**Supplementary Table 7. Cross-linking Sites in Bovine ROS-GC1 Fragment (aa 814-1110).** For the nomenclature of ROS-GC1 domains, please see **Fig. 1b**. Cross-links derived from ROS-GC1 monomeric gel bands are marked as intraprotein (intra) cross-links. Cross-links between identical or overlapping sequences originating from dimeric gel bands are indicated as interprotein (inter) cross-links between two ROS-GC1 monomers. If a cross-linking site is ambiguous, all potential cross-linked amino acids are listed.

| **Cross-linking site 1** | **Cross-linking site 2** | **Domains** | **Type of cross-link** |
| --- | --- | --- | --- |
| 818 | 1051 | αHD - CD | intra |
| 853/854 | 1048/1051 | CD - CD | intra |
| 1048 | 1098 | CD - CD | intra |
| 1051 | 1098 | CD - CD | intra |
| 1053 | 1073 | CD - CD | intra |
| 1053 | 1098 | CD - CD | intra |
| 1053 | 1110 | CD - CD | intra |
| 1069 | 1051 | CD - CD | intra |
| 1069 | 1098 | CD - CD | intra |
| 1073 | 1051 | CD - CD | intra |
| 1073 | 1098 | CD - CD | intra |
| 1073 | 1110 | CD - CD | intra |
| 1101 | 1051 | CD - CD | intra |
| 1110 | 1048 | CD - CD | intra |
| 1110 | 1051 | CD - CD | intra |
| 1110 | 1098 | CD - CD | intra |
| 1051 | 1051 | CD - CD | inter |
| 1053 | 1051 | CD - CD | inter |
| 1053 | 1048/1051 | CD - CD | inter |
| 1101 | 1098 | CD - CD | inter |
| 1110 | 1110 | CD - CD | inter |
| 818 | 1051 | αHD - CD |  |
| 818 | 1098 | αHD - CD |  |
| 853/854 | 1048/1051 | CD - CD |  |
| 853/854 | 1110 | CD - CD |  |
| 1048 | 1098 | CD - CD |  |
| 1051 | 1098 | CD - CD |  |
| 1053 | 1098 | CD - CD |  |
| 1053 | 1110 | CD - CD |  |
| 1069 | 1048 | CD - CD |  |
| 1069 | 1051 | CD - CD |  |
| 1073 | 1051 | CD - CD |  |
| 1073 | 1110 | CD - CD |  |
| 1101 | 1051 | CD - CD |  |
| 1110 | 1048/1051 | CD - CD |  |
| 1110 | 1051 | CD - CD |  |
| 1110 | 1098 | CD - CD |  |

**Supplementary Table 8.** **DSBU Cross-links in Bovine ROS-GC1 Fragment (aa 814-1110).** All cross-links were obtained in the presence of GCAP-2 and in the absence of claium. If a cross-linking site is ambiguous, all potential cross-linked amino acids are listed.

| **DSBU** | | | | | | | | |
| --- | --- | --- | --- | --- | --- | --- | --- | --- |
|  | **[M+H]^+^** |  | **deviation** | **cross-linked** |  | **cross-linked** |  |  |
| ***m/z*** | **theor.** | **z** | **(ppm)** | **peptide (1)** | **site (1)** | **peptide (2)** | **site (2)** |  |
| 407.028 | 2031.109 | 5 | 0.32 | [ARPGQFSGK} | 1110 | [GRTELKGK] | 1051 | intra |
| 508.533 | 2031.109 | 4 | -0.42 | [ARPGQFSGK} | 1110 | [GRTELKGK] | 1048 | intra |
| 486.775 | 1944.077 | 4 | 0.02 | [ARPGQFSGK} | 1110 | [RQKLEK] | 1098 | intra |
| 455.252 | 1817.987 | 4 | -1 | [ARPGQFSGK} | 1110 | [TELKGK] | 1051 | intra |
| 596.664 | 1787.976 | 3 | -0.15 | [ARPGQFSGK} | 1110 | [QKLEK] | 1098 | intra |
| 358.401 | 1787.976 | 5 | -0.2 | [ARPGQFSGK} | 1110 | [QKLEK] | 1098 | intra |
| 447.749 | 1787.976 | 4 | -1.11 | [ARPGQFSGK} | 1110 | [QKLEK] | 1098 | intra |
| 447.749 | 1787.976 | 4 | -0.84 | [ARPGQFSGK} | 1110 | [QKLEK] | 1098 | intra |
| 458.456 | 2288.247 | 5 | 0.91 | [EKARPGQFSGK} | 1101 | [GRTELKGK] | 1051 | intra |
| 492.035 | 1965.117 | 4 | 0.53 | [GFNKPIPKPP] | 1073 | [TELKGK] | 1051 | intra |
| 728.645 | 2911.558 | 4 | -0.07 | [GKGAEETYWLVGR] | 1053 | [RGFNKPIPKPP] | 1073 | intra |
| 919.158 | 2755.457 | 3 | 1.21 | [GKGAEETYWLVGR] | 1053 | [GFNKPIPKPP] | 1073 | intra |
| 870.115 | 2608.327 | 3 | 1.65 | [GKGAEETYWLVGR] | 1053 | [ARPGQFSGK} | 1110 | intra |
| 577.311 | 2306.214 | 4 | 2.79 | [GKGAEETYWLVGR] | 1053 | [QKLEK] | 1098 | intra |
| 433.493 | 1730.951 | 4 | -0.11 | [GRTELKGK] | 1048/1051 | [QKTDR] | 853/854 | intra |
| 433.005 | 1728.997 | 4 | 0.11 | [GRTELKGK] | 1051 | [QKLEK] | 1098 | intra |
| 433.005 | 1728.997 | 4 | -0.03 | [GRTELKGK] | 1051 | [QKLEK] | 1098 | intra |
| 461.657 | 2304.253 | 5 | 0.98 | [KMNIIDSMLR] | 818 | [GRTELKGK] | 1051 | intra |
| 523.539 | 2091.13 | 4 | 2.55 | [KMNIIDSMLR] | 818 | [TELKGK] | 1051 | intra |
| 530.897 | 2650.458 | 5 | 0.34 | [RGFNKPIPKPP] | 1073 | [EKARPGQFSGK} | 1110 | intra |
| 479.47 | 2393.32 | 5 | -0.22 | [RGFNKPIPKPP] | 1073 | [ARPGQFSGK} | 1110 | intra |
| 599.085 | 2393.32 | 4 | -0.75 | [RGFNKPIPKPP] | 1073 | [ARPGQFSGK} | 1110 | intra |
| 599.085 | 2393.32 | 4 | -0.96 | [RGFNKPIPKPP] | 1073 | [ARPGQFSGK} | 1110 | intra |
| 584.341 | 2334.34 | 4 | 1.47 | [RGFNKPIPKPP] | 1073 | [GRTELKGK] | 1051 | intra |
| 584.342 | 2334.34 | 4 | 2.2 | [RGFNKPIPKPP] | 1073 | [GRTELKGK] | 1051 | intra |
| 584.341 | 2334.34 | 4 | 1.47 | [RGFNKPIPKPP] | 1069 | [GRTELKGK] | 1051 | intra |
| 584.342 | 2334.34 | 4 | 2.2 | [RGFNKPIPKPP] | 1069 | [GRTELKGK] | 1051 | intra |
| 467.674 | 2334.34 | 5 | -0.18 | [RGFNKPIPKPP] | 1069 | [GRTELKGK] | 1051 | intra |
| 467.674 | 2334.34 | 5 | -0.8 | [RGFNKPIPKPP] | 1069 | [GRTELKGK] | 1051 | intra |
| 467.674 | 2334.34 | 5 | -0.18 | [RGFNKPIPKPP] | 1073 | [GRTELKGK] | 1051 | intra |
| 467.674 | 2334.34 | 5 | -0.18 | [RGFNKPIPKPP] | 1073 | [GRTELKGK] | 1051 | intra |
| 531.06 | 2121.218 | 4 | -0.21 | [RGFNKPIPKPP] | 1069 | [TELKGK] | 1051 | intra |
| 523.557 | 2091.207 | 4 | 0.28 | [RGFNKPIPKPP] | 1069 | [QKLEK] | 1098 | intra |
| 523.557 | 2091.207 | 4 | 0.05 | [RGFNKPIPKPP] | 1069 | [QKLEK] | 1098 | intra |
| 523.557 | 2091.207 | 4 | -0.42 | [RGFNKPIPKPP] | 1073 | [QKLEK] | 1098 | intra |
| 523.557 | 2091.207 | 4 | -0.54 | [RGFNKPIPKPP] | 1073 | [QKLEK] | 1098 | intra |
| 459.273 | 1834.07 | 4 | 0.78 | [RGFNKPIPKPP] | 1069 | [QKL] | 1098 | intra |
| 505.965 | 1515.874 | 3 | 4.13 | [TELKGK] | 1051 | [QKLEK] | 1098 | intra |
| 379.724 | 1515.874 | 4 | -0.43 | [TELKGK] | 1051 | [QKLEK] | 1098 | intra |
| 695.128 | 2777.483 | 4 | 2.31 | [TELKGKGAEETYWLVGR] | 1048 | [QKLEK] | 1098 | intra |
| 418.823 | 2090.089 | 5 | 0.9 | [ARPGQFSGK} | 1110 | [ARPGQFSGK} | 1110 | inter |
| 418.824 | 2090.089 | 5 | 0.63 | [ARPGQFSGK} | 1110 | [ARPGQFSGK} | 1110 | inter |
| 523.278 | 2090.089 | 4 | 1.18 | [ARPGQFSGK} | 1110 | [ARPGQFSGK} | 1110 | inter |
| 523.278 | 2090.089 | 4 | -0.11 | [ARPGQFSGK} | 1110 | [ARPGQFSGK} | 1110 | inter |
| 587.562 | 2347.227 | 4 | 0.07 | [EKARPGQFSGK} | 1110 | [ARPGQFSGK} | 1110 | inter |
| 638.093 | 2549.347 | 4 | 0.77 | [GKGAEETYWLVGR] | 1053 | [GRTELKGK] | 1048/1051 | inter |
| 638.092 | 2549.347 | 4 | -0.38 | [GKGAEETYWLVGR] | 1053 | [GRTELKGK] | 1051 | inter |
| 779.414 | 2336.224 | 3 | 0.96 | [GKGAEETYWLVGR] | 1053 | [TELKGK] | 1051 | inter |
| 779.414 | 2336.224 | 3 | 1.35 | [GKGAEETYWLVGR] | 1053 | [TELKGK] | 1051 | inter |
| 395.231 | 1972.13 | 5 | -1 | [GRTELKGK] | 1051 | [GRTELKGK] | 1051 | inter |
| 395.232 | 1972.13 | 5 | -0.15 | [GRTELKGK] | 1051 | [GRTELKGK] | 1051 | inter |
| 493.787 | 1972.13 | 4 | -1.21 | [GRTELKGK] | 1051 | [GRTELKGK] | 1051 | inter |
| 493.788 | 1972.13 | 4 | -0.5 | [GRTELKGK] | 1051 | [GRTELKGK] | 1051 | inter |
| 440.507 | 1759.007 | 4 | -0.07 | [GRTELKGK] | 1051 | [TELKGK] | 1051 | inter |
| 440.508 | 1759.007 | 4 | 1.18 | [GRTELKGK] | 1051 | [TELKGK] | 1051 | inter |
| 587.007 | 1759.007 | 3 | -0.1 | [GRTELKGK] | 1051 | [TELKGK] | 1051 | inter |
| 492.868 | 2460.311 | 5 | 1.13 | [LEKARPGQFSGK} | 1110 | [ARPGQFSGK} | 1110 | inter |
| 615.834 | 2460.311 | 4 | 0.82 | [LEKARPGQFSGK} | 1110 | [ARPGQFSGK} | 1110 | inter |
| 615.834 | 2460.311 | 4 | 0.92 | [LEKARPGQFSGK} | 1110 | [ARPGQFSGK} | 1110 | inter |
| 540.305 | 2158.198 | 4 | 0.8 | [LEKARPGQFSGK} | 1101 | [QKLEK] | 1098 | inter |
| 544.099 | 2716.464 | 5 | 0.41 | [QKLEKARPGQFSGK} | 1110 | [ARPGQFSGK} | 1110 | inter |
| 544.099 | 2716.464 | 5 | -0.04 | [QKLEKARPGQFSGK} | 1110 | [ARPGQFSGK} | 1110 | inter |
| 387.226 | 1545.885 | 4 | -2.83 | [TELKGK] | 1051 | [TELKGK] | 1051 | inter |
| 515.965 | 1545.885 | 3 | -2.43 | [TELKGK] | 1051 | [TELKGK] | 1051 | inter |
| 508.533 | 2031.109 | 4 | 0.24 | [ARPGQFSGK} | 1110 | [GRTELKGK] | 1048/1051 | intra or inter |
| 407.027 | 2031.109 | 5 | -1.1 | [ARPGQFSGK} | 1110 | [GRTELKGK] | 1051 | intra or inter |
| 407.028 | 2031.109 | 5 | 0.32 | [ARPGQFSGK} | 1110 | [GRTELKGK] | 1051 | intra or inter |
| 510.952 | 1530.839 | 3 | 1.67 | [ARPGQFSGK} | 1110 | [QKL] | 1098 | intra or inter |
| 596.663 | 1787.976 | 3 | -0.77 | [ARPGQFSGK} | 1110 | [QKLEK] | 1098 | intra or inter |
| 596.664 | 1787.976 | 3 | 0.46 | [ARPGQFSGK} | 1110 | [QKLEK] | 1098 | intra or inter |
| 596.665 | 1787.976 | 3 | 1.9 | [ARPGQFSGK} | 1110 | [QKLEK] | 1098 | intra or inter |
| 596.664 | 1787.976 | 3 | -0.05 | [ARPGQFSGK} | 1110 | [QKLEK] | 1098 | intra or inter |
| 596.664 | 1787.976 | 3 | 0.98 | [ARPGQFSGK} | 1110 | [QKLEK] | 1098 | intra or inter |
| 447.749 | 1787.976 | 4 | -1.52 | [ARPGQFSGK} | 1110 | [QKLEK] | 1098 | intra or inter |
| 447.749 | 1787.976 | 4 | -1.05 | [ARPGQFSGK} | 1110 | [QKLEK] | 1098 | intra or inter |
| 447.749 | 1787.976 | 4 | -0.98 | [ARPGQFSGK} | 1110 | [QKLEK] | 1098 | intra or inter |
| 447.749 | 1787.976 | 4 | -2 | [ARPGQFSGK} | 1110 | [QKLEK] | 1098 | intra or inter |
| 506.939 | 1518.802 | 3 | 0.32 | [ARPGQFSGK} | 1110 | [QKT] | 853/854 | intra or inter |
| 506.94 | 1518.802 | 3 | 1.1 | [ARPGQFSGK} | 1110 | [QKT] | 853/854 | intra or inter |
| 455.252 | 1817.987 | 4 | -1.07 | [ARPGQFSGK} | 1110 | [TELKGK] | 1051 | intra or inter |
| 606.667 | 1817.987 | 3 | -0.02 | [ARPGQFSGK} | 1110 | [TELKGK] | 1051 | intra or inter |
| 606.667 | 1817.987 | 3 | 0.39 | [ARPGQFSGK} | 1110 | [TELKGK] | 1051 | intra or inter |
| 606.667 | 1817.987 | 3 | 0.39 | [ARPGQFSGK} | 1110 | [TELKGK] | 1051 | intra or inter |
| 436.454 | 2178.239 | 5 | 0.73 | [GFNKPIPKPP] | 1069 | [GRTELKGK] | 1048 | intra or inter |
| 492.035 | 1965.117 | 4 | 0.91 | [GFNKPIPKPP] | 1073 | [TELKGK] | 1051 | intra or inter |
| 652.837 | 2608.327 | 4 | 0.15 | [GKGAEETYWLVGR] | 1053 | [ARPGQFSGK} | 1110 | intra or inter |
| 652.837 | 2608.327 | 4 | 0.15 | [GKGAEETYWLVGR] | 1053 | [ARPGQFSGK} | 1110 | intra or inter |
| 683.698 | 2049.076 | 3 | 1.86 | [GKGAEETYWLVGR] | 1053 | [QKL] | 1098 | intra or inter |
| 769.41 | 2306.214 | 3 | 1.02 | [GKGAEETYWLVGR] | 1053 | [QKLEK] | 1098 | intra or inter |
| 769.41 | 2306.214 | 3 | 0.55 | [GKGAEETYWLVGR] | 1053 | [QKLEK] | 1098 | intra or inter |
| 433.005 | 1728.997 | 4 | 0.32 | [GRTELKGK] | 1051 | [QKLEK] | 1098 | intra or inter |
| 433.005 | 1728.997 | 4 | -0.24 | [GRTELKGK] | 1051 | [QKLEK] | 1098 | intra or inter |
| 433.493 | 1730.951 | 4 | -0.46 | [GRTELKGK] | 1051 | [QKTDR] | 853/854 | intra or inter |
| 516.036 | 2061.119 | 4 | 2.03 | [KMNIIDSMLR] | 818 | [QKLEK] | 1098 | intra or inter |
| 687.713 | 2061.119 | 3 | 2.16 | [KMNIIDSMLR] | 818 | [QKLEK] | 1098 | intra or inter |
| 697.715 | 2091.13 | 3 | 0.4 | [KMNIIDSMLR] | 818 | [TELKGK] | 1051 | intra or inter |
| 697.717 | 2091.13 | 3 | 3.03 | [KMNIIDSMLR] | 818 | [TELKGK] | 1051 | intra or inter |
| 697.717 | 2091.13 | 3 | 2.68 | [KMNIIDSMLR] | 818 | [TELKGK] | 1051 | intra or inter |
| 547.808 | 2188.208 | 4 | 0.2 | [LEKARPGQFSGK} | 1101 | [TELKGK] | 1051 | intra or inter |
| 479.47 | 2393.32 | 5 | 0.22 | [RGFNKPIPKPP] | 1073 | [ARPGQFSGK} | 1110 | intra or inter |
| 479.47 | 2393.32 | 5 | 0.03 | [RGFNKPIPKPP] | 1073 | [ARPGQFSGK} | 1110 | intra or inter |
| 599.085 | 2393.32 | 4 | -0.24 | [RGFNKPIPKPP] | 1073 | [ARPGQFSGK} | 1110 | intra or inter |
| 467.674 | 2334.34 | 5 | 0.34 | [RGFNKPIPKPP] | 1073 | [GRTELKGK] | 1051 | intra or inter |
| 467.674 | 2334.34 | 5 | -0.64 | [RGFNKPIPKPP] | 1069 | [GRTELKGK] | 1051 | intra or inter |
| 467.674 | 2334.34 | 5 | 2.04 | [RGFNKPIPKPP] | 1069 | [GRTELKGK] | 1051 | intra or inter |
| 584.342 | 2334.34 | 4 | 2.31 | [RGFNKPIPKPP] | 1073 | [GRTELKGK] | 1051 | intra or inter |
| 531.061 | 2121.218 | 4 | 1.4 | [RGFNKPIPKPP] | 1069 | [TELKGK] | 1051 | intra or inter |
| 531.061 | 2121.218 | 4 | 1.51 | [RGFNKPIPKPP] | 1069 | [TELKGK] | 1051 | intra or inter |
| 505.963 | 1515.874 | 3 | 0.93 | [TELKGK] | 1051 | [QKLEK] | 1098 | intra or inter |
| 505.964 | 1515.874 | 3 | 1.59 | [TELKGK] | 1051 | [QKLEK] | 1098 | intra or inter |
| 379.724 | 1515.874 | 4 | 0.06 | [TELKGK] | 1051 | [QKLEK] | 1098 | intra or inter |
| 379.724 | 1515.874 | 4 | 0.94 | [TELKGK] | 1051 | [QKLEK] | 1098 | intra or inter |
| 695.128 | 2777.483 | 4 | 1.79 | [TELKGKGAEETYWLVGR] | 1048 | [QKLEK] | 1098 | intra or inter |

**Supplementary Table 9. HADDOCK Statistics for C2 Binary Docking of KHD.**

| HADDOCK score (a.u.) | -194.7 ± 0.7 |
| --- | --- |
| Cluster size (N) | 200 |
| RMSD from the overall lowest-energy structure (Å) | 0.5 ± 0.3 |
| Van der Waals energy (kcal.mol^-1^) | -95.9 ± 5.0 |
| Electrostatic energy (kcal.mol^-1^) | -323.6 ± 26.5 |
| Desolvation energy (kcal.mol^-1^) | -46.8 ± 3.7 |
| Restraints violation energy (a.u.) | 126.7 ± 23.19 |
| Buried Surface Area (Å^2^) | 3027.9 ± 97.6 |

Supplementary Table 10. Cross-link Distances in Bovine ROS-GC1 (IcD Model). The shortest distance was picked for all possible combinations (A-A, A-B, B-A, B-B). Chain combinations are given in columns 2 and 3. The given distances are calculated as Euclidean distances between Cα atoms of the residues involved. Cross-links are classified by a maximum distance threshold of 30 Å.

| ID | Chain/Residue 1 | Chain/Residue 2 | Euclidean Distance [Å] | Classification |
| --- | --- | --- | --- | --- |
| 1 | A/642 | A/818 | 15.27 | satisfied |
| 2 | A/657 | A/673 | 15.82 | satisfied |
| 3 | A/657 | A/674 | 12.6 | satisfied |
| 4 | A/657 | B/815 | 15.38 | satisfied |
| 5 | A/657 | B/815 | 15.38 | satisfied |
| 6 | B/673 | B/686 | 5.91 | satisfied |
| 7 | B/673 | B/688 | 7.43 | satisfied |
| 8 | A/673 | A/818 | 31.23 | violated |
| 9 | A/674 | A/818 | 30.1 | violated |
| 10 | B/686 | B/763 | 25.24 | satisfied |
| 11 | B/688 | B/763 | 23.28 | satisfied |
| 12 | A/686 | A/818 | 30.56 | violated |
| 13 | A/688 | A/818 | 36.15 | violated |
| 14 | B/763 | A/818 | 36.73 | violated |
| 15 | A/811 | B/818 | 8.2 | satisfied |
| 16 | A/686 | B/686 | 25.88 | satisfied |
| 17 | B/686 | A/688 | 28.47 | satisfied |
| 18 | A/811 | B/818 | 8.2 | satisfied |
| 19 | A/812 | B/818 | 9.8 | satisfied |
| 20 | B/815 | A/818 | 17.88 | satisfied |
| 21 | A/815 | B/824 | 16.47 | satisfied |
| 22 | A/818 | B/818 | 14.69 | satisfied |
| 23 | A/851 | B/853 | 11.37 | satisfied |
| 24 | A/851 | B/854 | 15.17 | satisfied |

Supplementary Table 11. Cross-link Distances in Human ROS-GC1 (IcD Model). The shortest distance was picked for all possible combinations (A-A, A-B, B-A, B-B). Chain combinations are given in columns 2 and 3. The given distances are calculated as Euclidean distances between Cα atoms of the residues involved. Cross-links are classified by a maximum distance threshold of 30 Å.

| ID | Chain/Residue 1 | Chain/Residue 2 | Euclidean Distance [Å] | Classification |
| --- | --- | --- | --- | --- |
| 1 | A/568 | A/657 | 27.29 | satisfied |
| 2 | B/642 | B/815 | 10.76 | satisfied |
| 3 | A/657 | A/673 | 15.82 | satisfied |
| 4 | A/657 | A/674 | 12.6 | satisfied |
| 5 | A/657 | B/815 | 15.38 | satisfied |
| 6 | B/657 | A/818 | 21.13 | satisfied |
| 7 | B/657 | A/819 | 19.64 | satisfied |
| 8 | A/673 | A/700 | 25.35 | satisfied |
| 9 | A/674 | A/700 | 26.12 | satisfied |
| 10 | B/815 | A/818 | 17.88 | satisfied |
| 11 | A/851 | B/853 | 11.37 | satisfied |
| 12 | A/1051 | B/1051 | 24.43 | satisfied |
| 13 | A/1048 | A/1053 | 28.47 | satisfied |
| 14 | A/1051 | B/1053 | 22.41 | satisfied |

Supplementary Table 12. Cross-link Distances in Bovine ROS-GC1 Fragment (aa 814-1110, IcD model). The ROS-GC1 fragment comprises αHD and CD. The shortest distance was picked for all possible combinations (A-A, A-B, B-A, B-B). Chain combinations are given in columns 2 and 3. The given distances are calculated as Euclidean distances between Cα atoms of the residues involved. Cross-links are classified by a maximum distance threshold of 30 Å.

| ID | Chain/Residue 1 | Chain/Residue 2 | Euclidean Distance [Å] | Classification |
| --- | --- | --- | --- | --- |
| 1 | A/818 | A/1051 | 92.20 | violated |
| 2 | B/853 | B/1048 | 46.00 | violated |
| 3 | A/853 | A/1051 | 41.81 | violated |
| 4 | B/854 | B/1048 | 44.27 | violated |
| 5 | A/854 | B/1051 | 40.31 | violated |
| 6 | A/1051 | B/1051 | 24.43 | satisfied |
| 7 | B/1053 | A/1051 | 22.41 | satisfied |
| 8 | B/1053 | A/1048 | 28.47 | satisfied |
